# Robustness vs productivity during evolutionary community assembly: short-term synergies and long-term trade-offs

**DOI:** 10.1101/2022.10.14.512255

**Authors:** Vasco J. Lepori, Nicolas Loeuille, Rudolf P. Rohr

## Abstract

The realization that evolutionary feedbacks need to be considered to fully grasp ecological dynamics has sparked interest in the effect of evolution on community properties like coexistence and productivity. However, little is known about the evolution of community robustness and productivity along diversification processes in species-rich systems. We leverage the recent structural approach to coexistence together with adaptive dynamics to study such properties and their relationships in a general trait-based model of competition on a niche axis. We show that the effects of coevolution on coexistence are two-fold and contrasting depending on the timescale considered. In the short term, evolution of niche differentiation strengthens coexistence, while long-term diversification leads to niche packing and decreased robustness. Moreover, we find that co-evolved communities tend to be on average more robust and more productive than non-evolutionary assemblages. We illustrate how our theoretical predictions echo in observed empirical patterns and the implications of our results for empiricists and applied ecologists. We suggest that some of our results such as the improved robustness of Evolutionary Stable Communities could be tested experimentally in suitable model systems.

## 1 Introduction

Ecology and evolutionary biology have long developed as separate disciplines [1], in spite of efforts throughout the years to better grasp the feedbacks that link variations in the biotic environment (inter- and intraspecific interactions) and evolutionary trajectories [2]. A series of empirical demonstrations of evolutionary changes happening over relatively short time-scales in different systems [3–5] fostered a renewed interest in the interface between ecology and evolution, an area of research now named eco-evolutionary dynamics [6]. At the same time, theoretical advances in the field of adaptive dynamics have shown that the feedback loop between ecological interactions and evolutionary change in characters can give rise to complex dynamics beyond simple optimization of growth rates, such as disruptive selection and evolutionary branching [7,8]. Yet, most ecological models for prediction and decision-making do away with evolution, considering it either negligible or too complex to grapple with [9,10]; and understanding the effects of evolution on the maintenance of species diversity largely remains an open question.

At the population level, evolutionary adaptation may save species from extinction under specific conditions, a phenomenon named evolutionary rescue [11]. But accounting for density- or frequency-dependent selection can open up scenarios where a species evolves towards self-extinction [12], a case dubbed evolutionary suicide in the adaptive dynamics literature [13]. In two-species scenarios, there can be both evolution towards stronger niche differentiation or one species can push the other to extinction [14]. Recently, Pastore et al. studied the effect of co-evolution of niche positions in a model of two competing species in the framework of Chesson’s coexistence theory [15–17]. Their work shows that evolution tends to have a negative effect on coexistence by increasing competitive imbalance, an outcome matching the experimental results of Hart et al. [18]. On the other hand, the quantitative genetics model of Barabás and D’Andrea suggest that communities where niche position and variation are heritable tend to exhibit more robust coexistence [19].

Hence it is not clear that evolution, which is driven by differences in individual fitness, will generally lead to improvement or even optimization of emergent properties at a larger organizational scale (here, communities). Brannström et al. highlighted what they call the dual nature of evolution in rich competitive communities [20], which can lead to the increase of diversity through generation of polymorphism and speciation but also to competitive exclusion and evolutionary murder. Classical ecological theory also predicts that stability in arbitrarily large systems may be difficult: in randomly interacting communities local dynamical stability decreased with richness [21], but adding non-random interactions such as adaptive foraging or eco-evo dynamics can counter this effect [22]. Loeuille showed that this effect is context-dependent in randomly assembled competitive and trophic communities, as evolution stabilizes moderately rich communities but destabilizes more species-rich systems [23].

Importantly, dynamical stability - the return to equilibrium after a state perturbation - is only half the picture [24], the other half being feasibility - the existence of an equilibrium state where all species have positive abundances, and its structural stability to small changes in the parameters [25,26]. The structural framework recently developed by Saavedra et al. allows us to quantify the tolerance to perturbations in the parameter space of intrinsic growth rates on an ecological timescale, such as those due to environmental perturbation, and disentangle the mechanisms underlying robustness [27]. The study of feasibility of competing species on a niche axis is also tightly linked to what classical theory calls species packing, a concept which dates back to Hutchinson’s work on species niches [28,29] and was mathematically formalized by MacArthur [30] who studied the limit in similarity between species, i.e., how close species can be on the niche axis while still coexisting. Case extended this approach to study limiting similarity under the constraint imposed by evolution [31].

If species interactions are important drivers of evolutionary change, we also expect that communities arising from co-evolution, for which Edwards et al. [32] suggested the term Evolutionary Stable Communities (ESCs), should comprise a highly non-random subset of all possible combinations of species. This has been advocated as early as in Rummel and Roughgarden [33], and more recently Aubree et al. showed that coevolved communities were generally more productive, more stable, and more resistant to invasion than collections which were randomly assembled from the species pool [34].

Here, we aim to study how evolution of niche positions and the emergence of polymorphism through evolutionary branching in a classical model of competition [35–37] interplay with coexistence and productivity constraints in diversifying species-rich systems. We use the structural stability approach to coexistence theory [27,38] to assess community robustness. Our aim is to go beyond previous works by 1) explicitly linking eco-evolutionary dynamics and structural coexistence metrics not only at the ESC but along evolutionary trajectories; 2) discussing how these links vary in contrasting ways depending on the time scale that is considered (rapid trait evolution vs diversification); 3) Uncovering and explaining the emergence of positive or negative correlations among various community properties, particularly diversity, productivity and coexistence, along evolutionary trajectories.

To do so, we follow the structural indicators of niche and fitness difference. These indicators aim to be species-rich analogs to the niche and fitness difference of Modern Coexistence Theory [27], but they are not one-to-one equivalents. For a given niche difference, small fitness differences increase robustness through equalizing effects; similarly, for a given fitness difference, niche differences increase robustness due to stabilizing effects. However, as is widely recognized, these two metrics do not exist independently [17,39] and we find they are insufficient to paint a complete picture of extinction risks. Therefore, we make use of a structural metric which quantifies the community resistance in the face of perturbations that would cause loss of one or more of its constituent species [38]. We track all three metrics along evolutionary trajectories. Finally, we consider changes in total community productivity, and contrast it with measures of coexistence. We compare these numbers to those of non-evolutionary communities of the same size.

We expect evolution to cause character displacement along the resource axis [35,40,41], which should first result in an increase of niche differentiation. At the same time, if divergence of niche positions results in a decrease in the growth rate of phenotypes situated further from the resource optimum, this should generate greater fitness differences. Increasing richness in a given resource space should on the other hand increase niche overlap [30] as well as fitness differences [42]. We therefore expect that effects of evolution on coexistence and productivity may vary depending on the environmental and temporal scales considered.

## 2 Materials and methods

### 2.1 Ecological dynamics

We study a niche-based model of competition based on generalized Lotka-Volterra dynamics following the *α*-*r* parametrization, whose advantages have been discussed by Mallet [43]. Each phenotype is defined by its position *µ*_*i*_ on a niche axis representing resources. Position on the niche axis affects both the intrinsic growth rates *r*(*µ*_*i*_) and the strength *α*(*µ*_*i*_, *µ*_*j*_) of density-dependent competition interactions between types. Versions of trait-based generalized Lotka-Volterra models and their eco-evolutionary dynamics have a long history in the literature [37,44–46], and have been shown to readily lead to the evolutionary emergence of polymorphism [35,36,47]. We use this behavior to explore the conditions of coexistence throughout the diversification process. Population dynamics of a type *i* then follow:

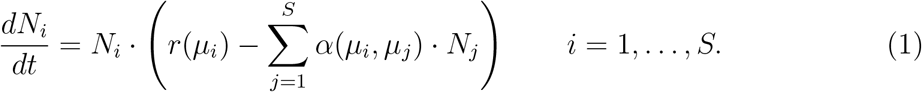

We assume a Gaussian function for *r*(*µ*_*i*_), with fecundity decreasing with distance from a resource optimum *µ*_*R*_

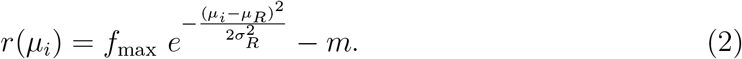

while including a small density-independent intrinsic mortality *m* to prevent phenotypes with unreasonably large or small trait values from persisting at virtually nil abundances, giving rise to the evolution of an infinite number of morphs whose coexistence has been shown to be structurally unstable, an outcome known as the problem of continuous coexistence [48–50]. The parameter *f*_max_ represents the maximum fecundity rate at the resource optimum, while *σ*_*R*_ depicts the width of resource availability on the niche axis. In line with previous work [40], we suppose that competition strength is defined by the similarity among types. It then follows a Gaussian-like function centered in *µ*_*i*_, so that *α*(*µ*_*i*_, *µ*_*j*_) = *α*(*µ*_*j*_, *µ*_*i*_), and interaction strength reaches its maximum *α*_max_ for *µ*_*i*_ = *µ*_*j*_:

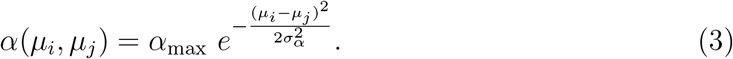

The parameter *σ*_*α*_ controls the width of the niche. This model is conceptually a size *S*-extension of the model used by Doebeli and Dieckmann [35,44]. The form of the competition function presents the agreeable property of being dissipative *sensu* Volterra [25,51]. This implies that ecological dynamics possess one and only one equilibrium point. Moreover, if this point is a feasible equilibrium 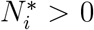 for all *i*, this equilibrium will be globally stable, i.e., all ecological dynamics will converge to it regardless of initial abundances. Note that a feasible equilibrium must fulfill 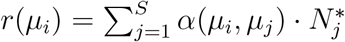 for each *i*.

### 2.2 Evolutionary dynamics

We study evolution within the adaptive dynamics framework [7,52], thereby accounting for both frequency- and density-dependent selection. The adaptive dynamics framework assumes clonal reproduction where mutations are infinitesimally small and rare, and a separation of timescales where ecology is assumed faster than evolution (but see Meszéna et al. on relaxing this assumption [36]), implying that advantageous mutations always go to fixation. The evolution of a quantitative trait is determined by the invasion fitness function, defined as the *per capita* growth rate of a rare mutant of traits 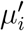 in a resident population of *S* phenotypes with traits *µ*_1_, …, *µ*_*S*_ at their ecological equilibrium [46]:

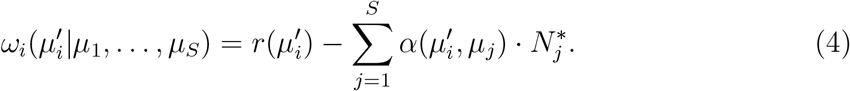

Note, that each phenotype *i* has its invasion fitness function *ω*_*i*_. The evolution over time of niche positions is determined by the canonical equation of adaptive dynamics [47]:

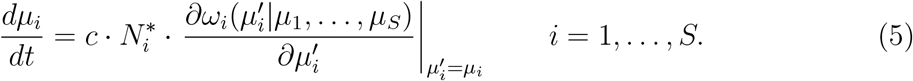

The partial derivatives

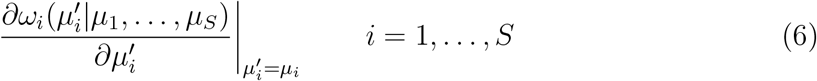

correspond to selection gradients determining the direction of co-evolution of the *S* niche positions. The coefficient *c* incorporates the per-capita mutation rate and the mutational variance, which we assume to be equal for all phenotypes. Without loss of generality, we can set *c* = 1 by rescaling the time axis. The selection gradients are also multiplied by the equilibrium abundances since larger population sizes lead to higher total chance of mutation [53]. We integrate the canonical equation (Equ. 5) numerically until an evolutionary singular strategy is reached, i.e, a set of traits 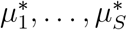, at which all selection gradients vanish

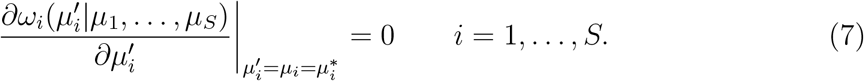

We start with a single monomorphic population and once we reach an evolutionary singular strategy, we evaluate the evolutionary stability condition (invasibility of the strategy) for each phenotype [7,52]. When the second derivative of the invasion fitness function (Equ. 4) with respect to the mutant trait is positive at the singular strategy the strategy is invadable [8]. Evolutionary branching ensues, and we introduce a new phenotype at a small distance of the singularity, augmenting the dimensionality of the system. We continue the simulation until all strategies are evolutionary stable, forming an Evolutionary Stable Community (ESC) *sensu Edwards et al*. [29]. The algorithm is detailed in the Supplementary Materials S1.

### 2.3 Structural coexistence metrics

Along co-evolutionary trajectories where *S ≥* 2, we assess coexistence metrics. Within the structural approach, coexistence in species-rich systems is quantified by structural niche differences Ω and structural fitness differences *θ*. The structural niche difference Ω quantifies the probability of coexistence given the interactions, and is mathematically defined as the solid angle of the domain of intrinsic growth rates leading to coexistence given the interaction strength *α*(*µ*_*i*_, *µ*_*j*_), defining the feasibility domain (Figure 1). In turn, the structural fitness difference *θ* quantifies the deviation from the center of the feasibility domain, i.e., to what extent one phenotype dominates the system. The fitness difference *θ* depends on both the intrinsic growth rates vector **r**(*µ*_*i*_) and the competition strength *α*(*µ*_*i*_, *µ*_*j*_). To compare structural values of niche difference across communities of different sizes *S*, we consider the standardized metric 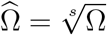 [54].

**Figure 1:**
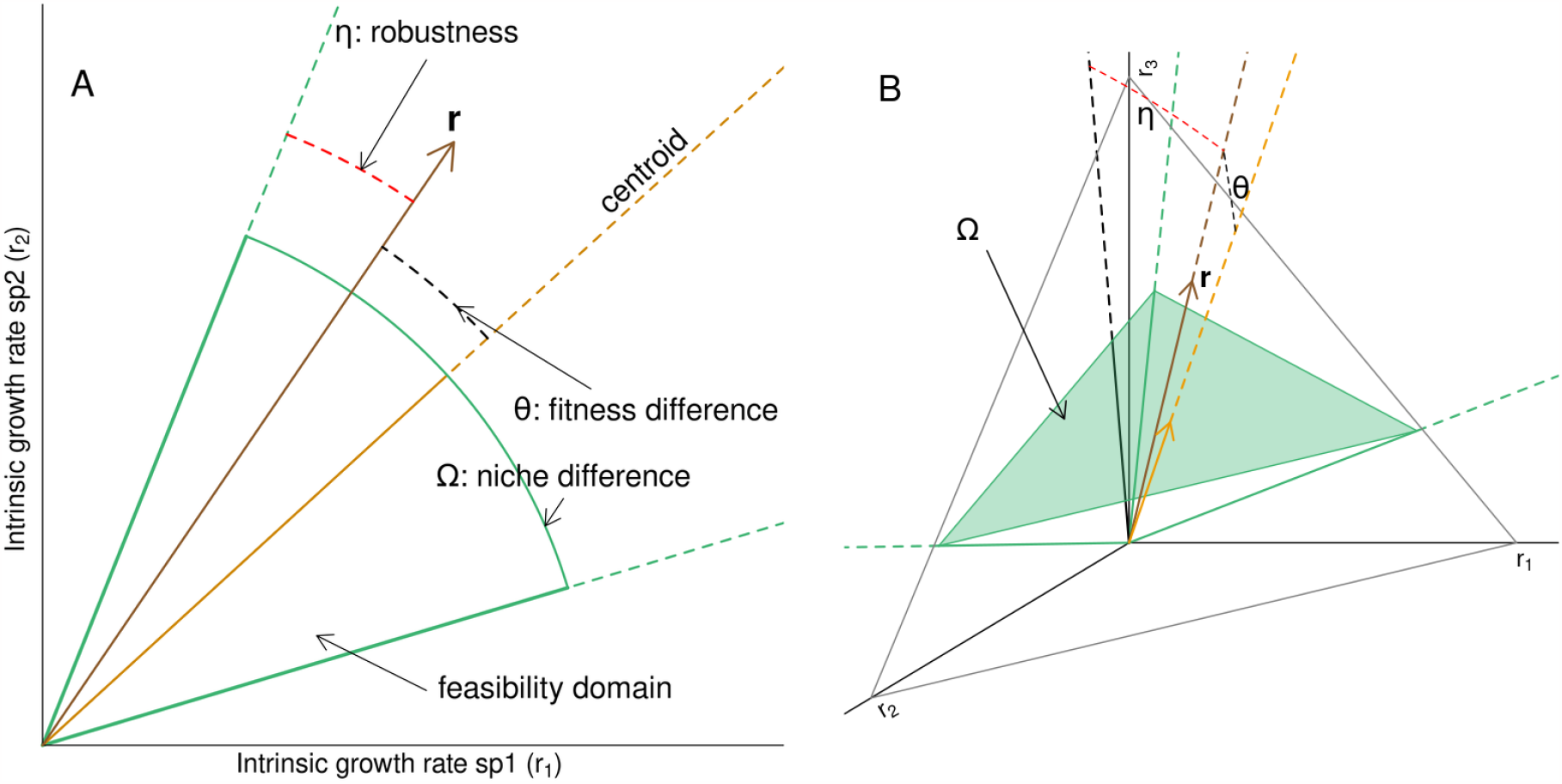
Geometric representation of the structural coexistence metrics for a 2-species system (panel A) and a 3-species system (panel B). On both panels, the green cone, called domain of feasibility, determines the set of intrinsic growth rates leading to a positive equilibrium for each species, i.e., to coexistence. The structural niche difference Ω is given by the measure of this solid angle, while the structural fitness difference *θ* measures the deviation of the vector of intrinsic growth rates (**r**) from the centroid of the cone. Finally, *η* – the structural measure of resistance – is given by the smallest angle between the vector of intrinsic growth rates and the border of the feasibility domain. It gives an indication of the fragility of the community with respect to perturbations on **r**. Note that we illustrated the structural metrics for *S* = 2 and *S* = 3, but they are effectively defined and computed for *S*-rich systems (though they become hard to represent).

### 2.4 Structural robustness metric

While niche and fitness differences allow for partitioning coexistence effects, they do not give a clear indication of the absolute strength of coexistence. For example, if both Ω and *θ* increase, we do not know whether coexistence is favored or weakened. Following Medeiros et al. [38] we utilize a complementary measure which we call *η*, the smallest angle between the vector of intrinsic growth rates **r** and the border of the feasibility domain as a measure of resistance to perturbations. This angle gives an indication of the fragility of the community with respect to perturbations on **r**, i.e., how close the community is to losing of one or more of its components. Like the other metrics, resistance is considered on an ecological timescale, with respect to perturbations that would impact growth rates through resources or the environment (rather than through trait change). Figure 1 provides representations of Ω, *θ*, and *η* in a system of *S* = 2, *S* = 3. The mathematics of how *η* is computed are detailed in the Supplementary Materials S2.

### 2.5 Productivity metrics

Besides coexistence metrics, we also measure the evolution of productivity using the proxy of total community biomass (or abundance) 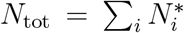 is [55]. Biomass diversity effects (overyielding of polymorphic communities compared to monocultures) can be due to complementarity in resource use, as well as selection for high-yield phenotypes (high carrying capacity *K*_*i*_ = *r*_*i*_*/α*_ii_) [56]. Both of these mechanisms can be easily captured under our model and are expected to vary over coevolutionary time.

### 2.6 Randomizations

We follow coexistence metrics and productivity along co-evolutionary trajectories and branching points. However, this does not tell us whether those properties are generally improved or even maximized by evolution. To this end, we generate between 4’000 and 128’000 (depending on *S*) communities for each singular strategy (branching point or stable strategy) of each simulation, by sampling uniformly sets of niche positions *µ*_*i*_, conditioned on the resulting community being feasible (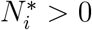 for all *i* = 1, …, *S*). We then compare the metrics (structural niche and fitness difference, resistance, and total biomass) of the sampled communities with the eco-evolutionary trajectories. We use two different rules to define the range of niche positions from which we sample. For the first rule, we sample communities within the range of niche positions observed at the ESC (restricted range), which tells us if the evolutionary community is unique within this bracket of niche positions. For the second rule (full range), the range of sampled *µ*_*i*_ covers all niche positions leading to positive intrinsic growth rate *r*(*µ*_*i*_) *>* 0. Such bounds are given by 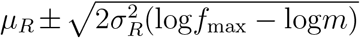 and allow us to compare strategies with respect to the whole trait space.

### 2.7 Choice of parameter values

To reduce the number of free parameters, we first transform the dynamical system into a nondimensional form. That is, we freely chose the time unit, the abundance unit, and the scale of the niche axis. Regarding the abundance and time unit, we can set without loss of generality *α*_max_ = 1, *f*_max_ = 1. The latter defines the ecological timescale, but adaptive dynamics assumes that ecological equilibrium is reached before the next mutation occurs. The combination of both determines the scale of species abundance and can arbitrarily be rescaled. The additional mortality rate *m* remains, and thus cannot be freely chosen. Regarding the niche axis, we can rescale, without loss of generality, its origin to *µ*_*R*_ = 0. Then we can also choose its scale by setting arbitrarily the resource width *σ*_*R*_ = 1 and explore the effect of the niche width *σ*_*α*_, or alternatively, we can also set arbitrarily the niche width *σ*_*α*_ = 1 and explore the effect of the resource width *σ*_*R*_.

For simplicity, in the main text we show the results for one specific parametrization, set to *m* = 0.01 and *σ*_*α*_ = 1, and *σ*_*R*_ = 1.6.

## 3 Results

### 3.1 Co-evolutionary trajectories and branching events

Figure 2A shows a co-evolutionary trajectory, which undergoes diversification until it reaches a stable and convergent community (ESC). After each branching, we observe a divergence in the niche positions, which leads to a decrease in the level of interspecific competition. The evolutionary endpoint for the monomorphic situation (i.e., before the first branching) can be determined analytically. It converges to an evolutionary singular strategy that is always located at the resource optimum *µ*^*∗*^ = *µ*_*R*_. Its invasibility depends on the width of the competition function (*σ*_*α*_), the width of the resources (*σ*_*R*_), the maximum fecundity rate (*f*_max_), and the mortality rate (*m*). We show that branching occurs if and only if 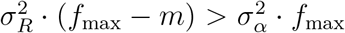 (Supplementary Materials S3). Branching will therefore occur if the resource range is wide enough (as defined by *σ*_*R*_) and/or limiting similarity strong enough (small *σ*_*α*_). This formula is alike the one derived by Dieckmann and Doebeli [35,44] but generalized to an additional mortality term, which has an evolutionary stabilizing effect through reducing the resident abundance at equilibrium. As expected by niche packing theory [30,40], the number of subsequent branching, and therefore, the number of species at the ESC, increases with the resource availability (*σ*_*R*_) and decreases with the competition width (*σ*_*α*_), see Supplementary Material S4.

**Figure 2:**
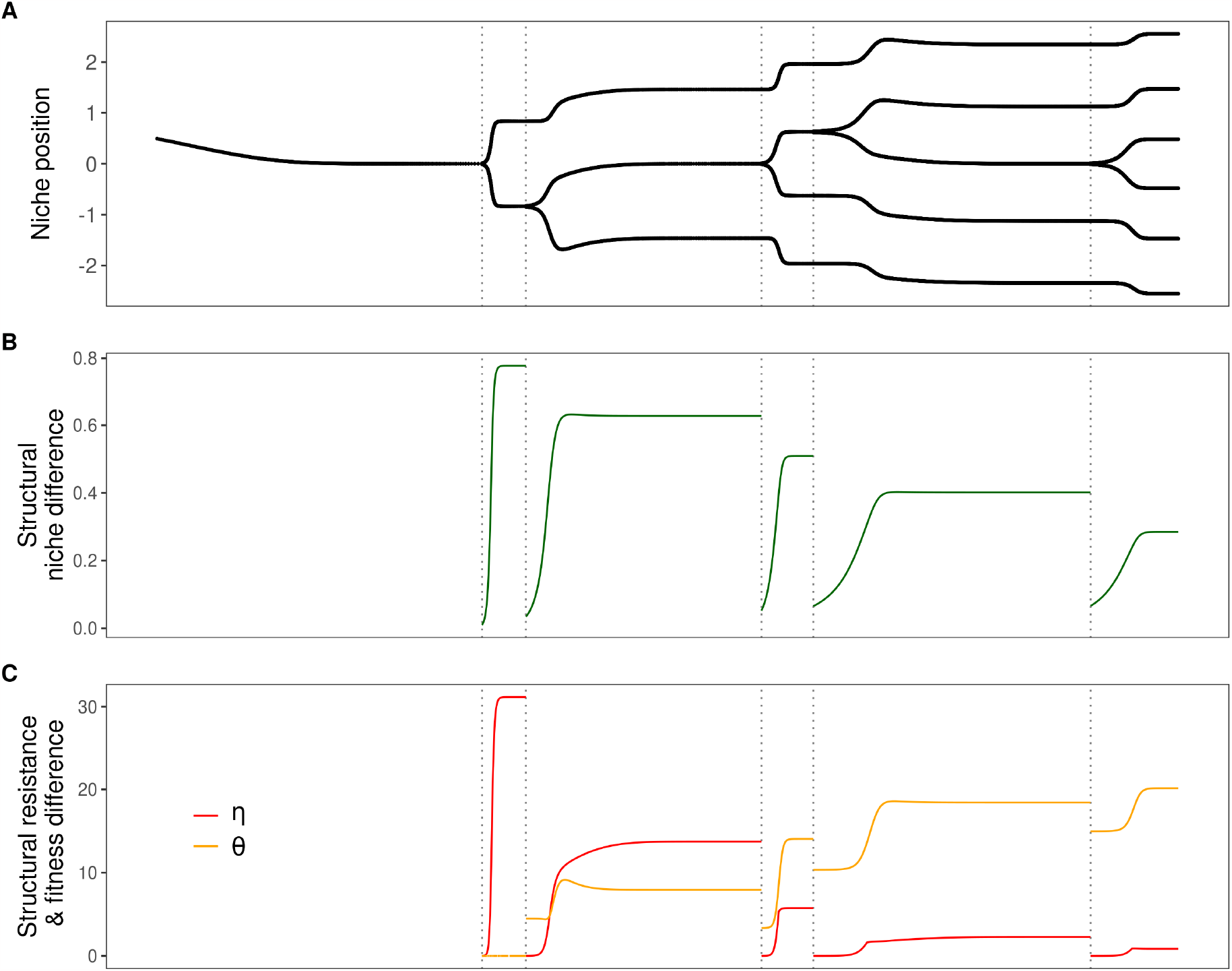
Evolutionary trajectory. Panel **A** shows the evolutionary trajectory of the niche positions *µ*_*i*_. The vertical dotted lines indicate evolutionary branching. Panel **B** shows the evolution of the structural niche difference 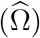. Panel **C** shows the evolution of structural fitness difference (*θ*) and resistance (*η*). For this figure, the resource width has been set to *σ*_*R*_ = 1.6, the niche width to *σ*_*α*_ = 1, and the mortality term to *m* = 0.1, but results are robust to different choices of parameters as shown in the Supplementary Materials.

Figure 2B shows that the structural niche difference 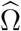 increases between two branching points as a consequence of diverging niche positions. But at the branching points, the addition of a new phenotype very close to an existing one causes the structural niche difference to abruptly decrease. Hence, for a period, coexistence is driven by more neutral (fitness-equalizing) mechanisms, rather than by strong niche separation. More specifically, branching points are the only contiguous trait regions where indefinitely close phenotypes can coexist, creating a niche-neutrality continuum *sensu Song et al*. [39]. Conversely, structural fitness differences *θ* increase along the entire evolutionary trajectory, undermining community robustness (Fig. 2C). The angle *θ* is initially zero when only two types coexist, as their trait positions are symmetric around the resource center. It later tends to increase for *S >* 2 as phenotypes start differing in their intrinsic growth rates, reflecting imbalances between strong and weak competitors. Because evolution here increases both niche differences (stabilizing effect) and fitness differences (unequalizing effect) between branching points, its overall effect on coexistence is not obvious. We therefore rely on the resistance indicator – the angle *η* – to assess the evolution of the distance to the border to the feasibility domain. Figure 2C shows that *η* tends to increase between the branching points, so that the overall effect of a simultaneous increase in both fitness and niche difference ultimately results in more robust communities. As expected, *η* decreases at each branching point due to increased competition and niche overlap in richer communities. This trend is observed even when r values are uniform rather than decreasing towards the resource edges (Supplementary Materials S8), showing that the loss of robustness is due to decrease in niche differences and not just increase in fitness differences. Moreover, as *S* increases and the community becomes saturated, the value of *η* becomes very small. Once the maximum phenotype packing is reached (ESC), small perturbations in **r** suffice to lead to non-feasibility, and the community is structurally more fragile than an undersaturated one (fewer phenotypes than at the ESC).

### 3.2 Projection into coexistence space

Figure 3 illustrates how structural coexistence metrics 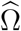 and *θ* change along the evolutionary trajectory of figure 2 when *S ≥* 2. Remember that large 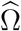 and small *θ* enhance persistence. In the background, we show the metrics for randomized communities that were produced according to the two rules presented in the Methods.

**Figure 3:**
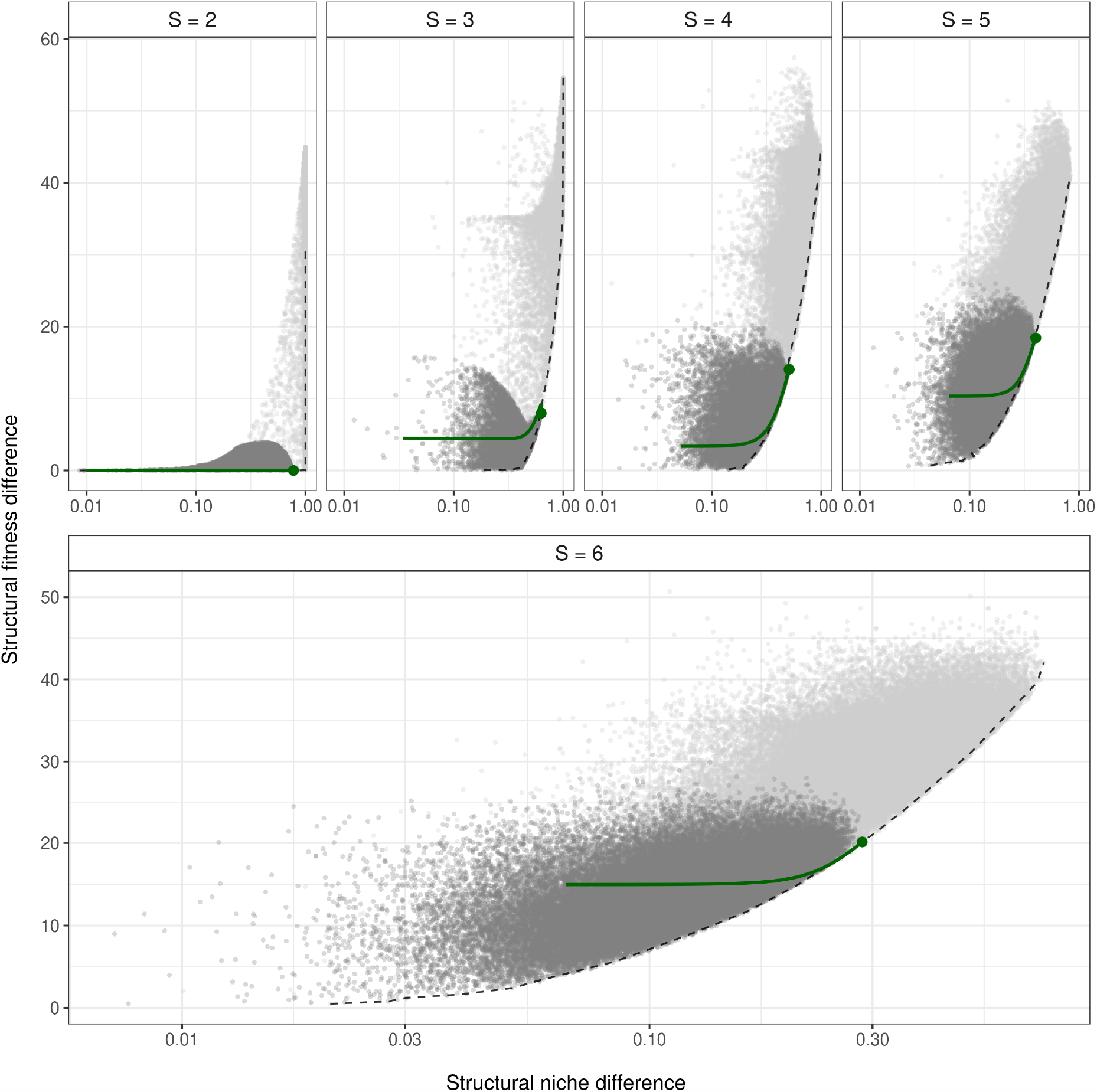
Variations of coexistence metrics along evolutionary trajectories. The figure is split into panels according to the number of phenotypes *S*. Each panel therefore represents a slice (in between two branching points) of the trajectory presented in figure 2. The green line represents the projection of the evolutionary trajectory of figure 2 into the space of structural coexistence metrics for a given *S*, while the larger dot marks the endpoint of the trajectory (singular strategy). Gray points represent randomized feasible communities, with niche position value sampled from the restricted range explored by evolution (1^st^ rule; dark gray) or the full range leading to positive intrinsic growth rates (2^nd^ rule; light gray). Interpolated pareto fronts among randomized communities are shown using dashed lines.

Considering all randomized communities shows that communities that have small fitness differences most often also have small niche differences. This is due to both selecting feasible communities, but also to ecological constraints under this model, which generate a global trade-off at the community level between the two coexistence metrics. Points that optimize one of the two properties relative to the other lie on a Pareto front. Figure 3 shows that evolution leads to this Pareto-optimality, and more specifically to the point on this front that also maximizes the structural niche difference within the restricted range (first randomization rule). More extreme niche positions would also allow larger niche differences, but this decrease in competition would come at the cost of decreasing intrinsic growth rates for the phenotypes further from the resource optimum, thereby increasing structural fitness differences *θ*.

Regarding community resistance (*η*), figure 4 indicates that evolution optimizes it within the range of niche positions explored by evolution (first randomization rule), but greater resistance could (rarely) be reached for the second randomization rules. However, among the niche positions explored by evolution and for a given number of phenotypes, evolution converges to more robust communities, by optimizing the structural niche difference Ω and the resistance *η*.

**Figure 4:**
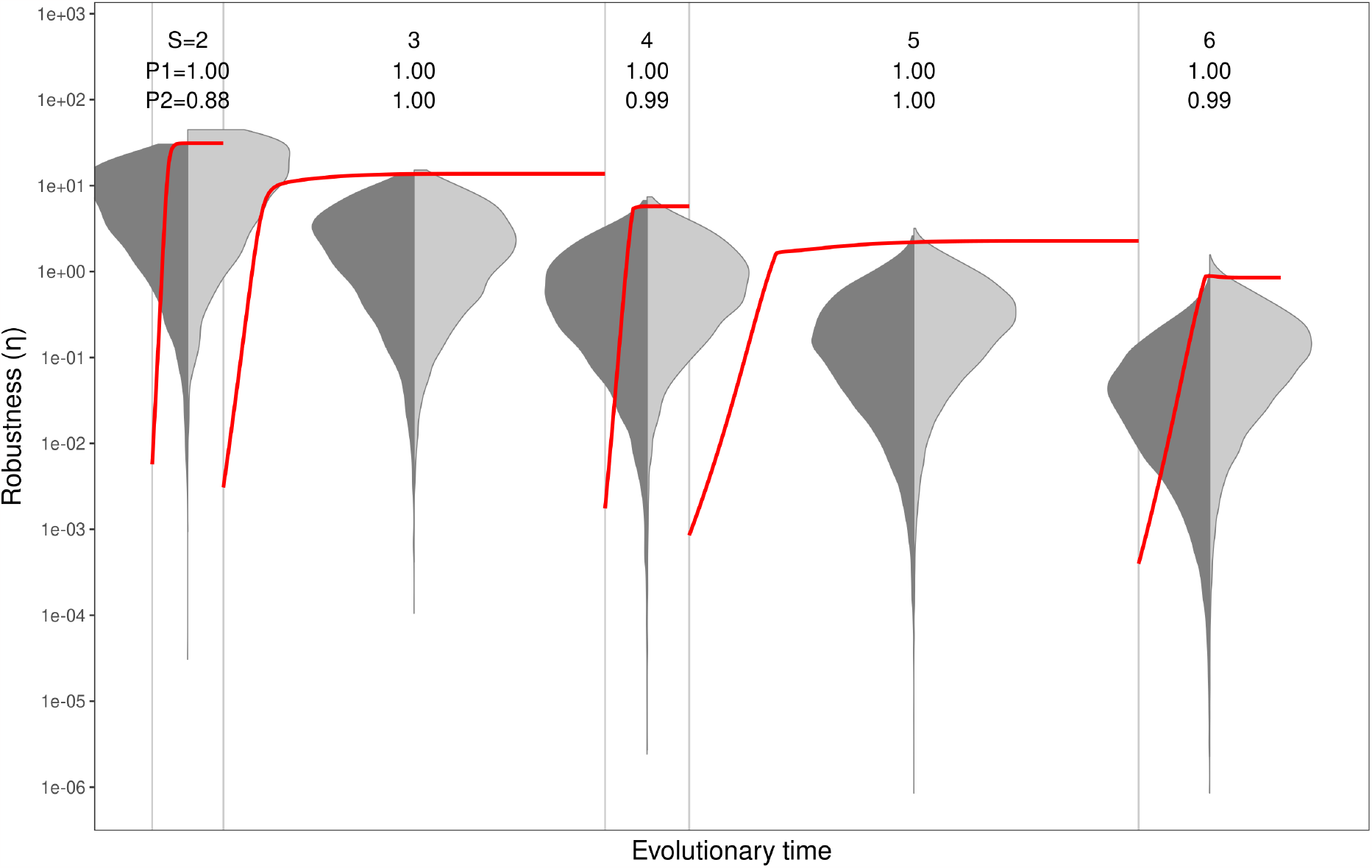
Evolution of community resistance. The red line shows the evolution of *η*. Gray points represent randomized feasible communities, with niche position value sampled from the restricted range explored by evolution (1^st^ rule; dark gray) or the full range leading to positive intrinsic growth rates (2^nd^ rule; light gray). At the singular strategy, the evolved community reaches a value of *η* greater or equal to than a proportion *P*1 and *P*2 of randomized communities according to the first and second randomization rules, respectively.

### 3.3 Evolution of productivity

In general, co-evolutionary dynamics increase productivity at every timescale, due mostly to increased richness but also due to increased niche differentiation and complementarity within a single level of diversity. While strict optimization can be proven in the monomorphic situation (Supplementary Materials S3), rich evolved communities are also among the most productive, as illustrated by figure 5.

**Figure 5:**
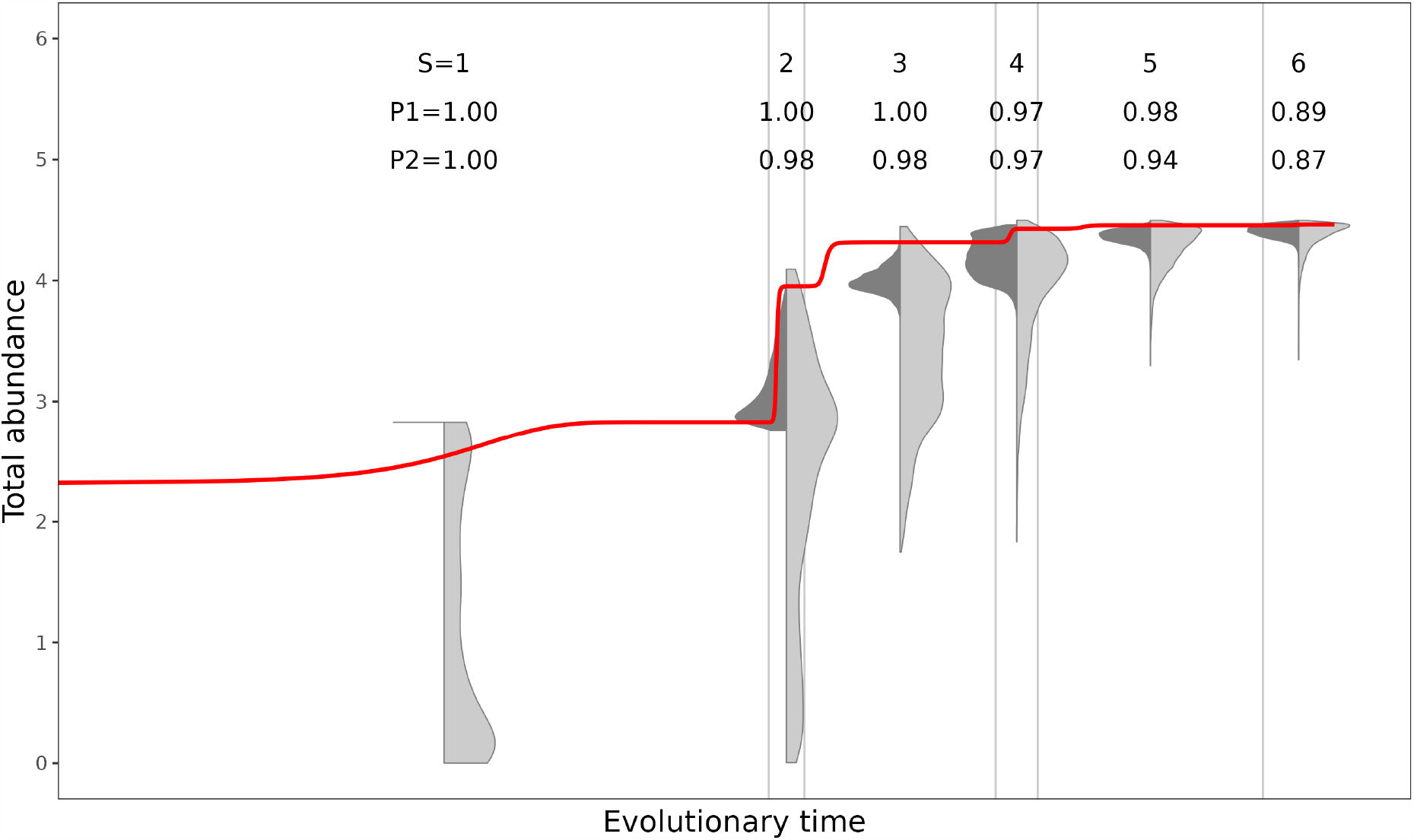
Productivity along evolutionary trajectories. Evolution of productivity. Here the numbers *P*1 and *P*2 in the second row of text indicate the percentage of randomized communities that are less productive than the ESC. For S=1, biomass production is optimized at the ESC. For S=6, the ESC is in the 87th percentile of the most productive communities. Point clouds represent randomization in the narrow (1st rule; dark gray) or the full range (2nd rule; light gray).

## 4 Discussion

Understanding the way evolution shapes community assembly and affects community properties, including robustness, is paramount. This includes untangling how evolutionary communities (ESCs) differ from non-coevolved assemblages. Previous studies showed that mechanisms such as adaptive foraging, which can be evolutionary in nature, can stabilize complex systems [22,59]. Recently a handful of studies, both theoretical and experimental, have explored this question for pairs of coevolving species within the framework of modern coexistence theory [15]. Here we employ recent theoretical advances to investigate the question for species-rich systems (*S >* 2) where diversity arises through subsequent branching events. By using a structural approach to coexistence, we can evaluate strength of coexistence as well as its partitioning structural niche difference and structural fitness differences as well as the resistance, along species-rich evolutionary trajectories. It shall be noted, however, that the structural approach is but one of several proposed frameworks to study coexistence in species-rich systems. Multi-species extensions of Modern Coexistence Theory have been proposed by Spaak and De Laender [60] and Carroll et al. [61]. These approaches, rooted in the concept of invasion analysis, only apply when invasion growth rates (not to be confused with the invasion fitness of adaptive dynamics mentioned earlier) correctly predict coexistence outcomes, as pointed out by Spaak and De Laender [60]. The structural approach, on the other hand, defines alternative multispecies coexistence metrics without relying on invasion growth rates, an assumption shown not to hold in some experimental systems such as the bacterial communities of Chang et al [62].

We show that evolution of coexistence properties follows two distinct trends on two different timescales, while productivity systematically increases along evolutionary trajectories. The short-term trend, in-between branching events, allows for increased efficiency in resource partitioning and decreased competition. On this timescale, evolution promotes structural niche differentiation 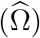, but this effect is counterbalanced by a simultaneous increase in structural fitness difference (*θ*) (Fig. 2). Hence, coexistence is enhanced by favoring niche partitioning, rather than by neutral mechanisms. We observe both an increase in the stabilizing mechanisms (increase in niche differences) and a reduction of the equalizing mechanism (in fitness differences) as in Pastore et al. [14], but we use the measure of resistance *η* introduced earlier to show that between branching communities, evolution selects less fragile communities with respect to environmental perturbations that impact growth rates **r**. The fact that coexistence metrics are usually not independent is already appreciated [17,39], but we here show how they are readily coupled throughout evolutionary dynamics. This link is made obvious in our trait-based model (Fig. 3), where niche position affects both growth rates and competition strength, and in turn Ω and *θ*. This interdependence makes it impossible to minimize simultaneously niche overlap and fitness differences (Fig. 3). Still, evolution leads to communities that are not strict optima of Ω nor minima of *θ* but rather lie on a Pareto front, an optimal compromise that is reached between niche increasing differentiation and restricting fitness imbalances. In a rare direct empirical test of this short term effect, Hart et al. found an increase in fitness difference after evolution, but not one in niche difference, which they explain may have been prevented by competition for a single discrete, non-substitutable resource [18]. Still, evidence of (evolutionary) niche differentiation measured as character displacement abounds in other natural and experimental settings [41,63], hence confirming that we should also expect evolution to increase both metrics in most systems.

Our results suggest that costs in terms of community robustness may happen over a longer timescale, where evolution leads to diversification along with evolutionary niche packing. The contrast between the short-term and long-term variation in community robustness is analogous to a case of Simpson’s paradox: there is an increase in robustness within communities of given size along evolutionary time but an overall decrease over longer timescales due to diversification events. Branching points, by virtue of addition of one more phenotype whose trait value and fitness are initially very close to an existing one, have a simultaneously equalizing and destabilizing effect. As the community size increases, resistance and niche differences peak at increasingly lower levels. This is a consequence of trying to pack a greater number of phenotypes in the same resource width *σ*_*R*_ (Fig. 2). Classical theory shows that the number of phenotypes that can be packed on a resource axis is a function of resource width *σ*_*R*_ relative to the niche width of species [40]. But when evolution is allowed in the community, the richness at the ESC is lower than under strict maximum (non-evolutionary) packing. In fact, ESCs are by definition uninvadable, because the fitness landscape for any possible invading trait value *µ*_*m*_ is zero at the resident trait values *µ*_1_, …, *µ*_*S*_ and negative everywhere else; however, richer, feasible non-evolutionary configurations do exist for a given environment. This result has been shown to be consistent across a range of models, for both continuous and discrete resources (e.g., [31,33,64]). It is worth pointing out that the answer to whether evolution helps or hinders coexistence is context-dependent: when the community is undersaturated (fewer members than at the ESC), evolution drives an increase in resistance *η*. Conversely, by starting with feasible but supersaturated communities (more diversity than allowed at the ESC), *η* will repeatedly drop to zero and we observe a sequence of extinction events until the ESC is reached. For example, Shoresh et al. found that evolution destabilizes communities, which can be explained by the fact that they started with supersaturated communities [64]. This also helps explain why Loeuille found evolution to be usually stabilizing for small communities (likely undersaturated), while its effect is reversed for rich communities (likely oversaturated) [23]. Empirical evidence for the longer-term effects of evolution on community properties is difficult to acquire, but could be found in phylogenetic patterns of niche conservatism [65]. For example, Yguel et al. report patterns consistent with increased productivity over time due to niche diversification and filling [66].

Ultimately, the evolutionary process results in ESCs that are highly non-random with respect to their structure and properties. Coevolved communities also show more evenly-spaced trait distributions than non-evolutionary communities in the model of Barabás and D’Andrea [19] (but see Bennett et al. for contrasting empirical evidence [67]). Our results show that they are also more structurally robust. This has important implications: if coevolved communities are more robust to environmental stresses (changes in temperature, water, nutrients), this could have important consequences in terms of guiding conservation efforts. Indeed, studies on the effects of global change often focus on single-species responses, but many have argued that community-level responses should be given more attention [57,68,69]. Because community composition and interactions are modified by climate change and invasive species, leading to new assemblages that likely depart from coevolved structures [70,71], our results suggest that the robustness of these new communities may be relatively poor. Further experimental testing of community-level responses to environmental stress in co-evolved versus randomized assemblages should prove an exciting and critically needed avenue of research, and the structural approach provides a useful theoretical framework to tackle the question. Experimental testing could for instance be undertaken using systems that rapidly diversify (e.g., the *Pseudomonas* system in [72]).

Regarding productivity, rich randomized communities tend to be more productive on average than poor ones (Fig. 5). By requiring randomized communities to be feasible, we however introduce a selection bias: in this sense, the positive slope between richness and productivity is indeed a byproduct of coexistence [73], since the conditions that promote coexistence (niche differences), are also those that promote complementarity and greater productivity. Thus, we observe the emergence of trade-offs between community properties: higher total biomass values are possible among randomized communities, but these tend to have larger fitness imbalance and/or lower community robustness. This finding echoes results of Rohr et al. showing that very productive communities have low evenness [74], while those with lower deviation from the feasibility domain center (*θ*) have intermediate levels of biomass production. In agreement with Aubree et al. [34], our results also show that evolutionary communities are often more productive on average than randomized ones, especially when species richness is low. Total abundance increases monotonically along evolutionary trajectories (no Simpson’s paradox of production), first because of selection for higher intrinsic growth rates in the monomorphic case (selection effect), then due to a better use of the resource space by the diversifying phenotypes (complementarity effect) (Fig. 2). This is consistent with the expectation that increased diversification within and across species should lead to occupation of vacant niche spaces, leading to increased niche complementarity and utilization, and ultimately increased abundances at the consumer level (a hypothesis supported by the phylogenetic analysis of Yguel et al. [66]). For instance, some of the largest biodiversity ecosystem-functioning experiments have consistently reported an increase in the net effect of biodiversity on biomass production and of niche complementarity across a decade ([75,76]). Similarly, Stefan et al. showed that plant-plant interactions shifted towards increased complementarity and yield over just a few generations of coexistence [77], while van Moorsel et al. report higher productivity in polycultures with an 8-year co-evolution history compared to identical-composition, but evolutionary naive plant communities [78]. This evidence strongly supports the idea that the complementarity effect often observed in biodiversity-ecosystem functioning experiments is expected from an evolutionary point of view, and likely reinforcing along evolutionary trajectories. A consequence of this result is that many BEF studies may have underestimated the productivity gains due to biodiversity, by assembling *de-novo* communities, instead of coevolved ones.

It is, however, possible to find more productive combinations among the random communities. Indeed, strict evolutionary optimization of total biomass at equilibrium only arises for the monomorphic case where the selection gradient coincides with the gradient of *r*(*µ*) (Supplementary Materials S3). In polymorphic systems, maximizing biomass productivity would require further niche differentiation, at the expense of increasing fitness imbalance, which is not achieved under the Pareto-optimality engendered by evolution. While this runs contrary to the adaptationist belief that evolution begets optimality, it is known that (abundance or growth rate) optimization is not the norm once we consider realistic eco-evolutionary feedback loops. This has been shown already by Abrams and Matsuda [12], then more rigorously in a series of papers by Metz and collaborators [79–81], and recently by Rohr and Loeuille [82]. The latter shows that biomass optimization is generally incompatible with niche differentiation and branching, with a few exceptions such as the monomorphic case of our model.

Several questions remain open for investigation. We considered only competitive interactions and a single resource axis, so that packing more than two species with the same amount of niche overlap between all of them is impossible [40]. If phenotypes were arranged in a multidimensional trait space, neutral configurations (in terms of **r**) with multiple species would be possible. In addition, we considered only evolving niche positions in our model, but niche width could also evolve as in the model of Ackermann and Doebeli [46], or in a slightly different form in Barabás and D’Andrea [19], Barabás et al. [83] and Wickman et al. [84]. Nevertheless, the theoretical predictions of our study need to be further experimentally tested. Although there is empirical support in biodiversity ecosystem-functioning studies regarding the increase in niche complementarity and biomass production over time, it remains unclear whether those predictions would hold in co-evolved communities emerging from diversification. Experimentally, comparison of properties of ESCs to non-evolutionary communities is complicated by the fact that ESCs are conceptually useful, but whether they are frequent in nature, and how to go about identifying them, is unclear [32]. Despite these hurdles, experimental tests of our theoretical results could provide timely evidence that would contribute to our understanding of the evolution of community properties in rich systems. Such experiments could be undertaken in microbial systems, where rapidly evolving communities could be compared to control treatments of non-coevolved assemblages [72,85].

## Data and code

Numerical simulations of eco-evolutionary trajectories were performed in Julia 1.5.1, while computation of metrics of coexistence and plotting of results were done in R 4.0.3. Code to reproduce the analyses is available on github.com/vlepori/structuralecoevo.

