## Supplementary Materials for "Robustness vs productivity during evolutionary community assembly: short-term synergies and long-term trade-offs"

### S1: Simulation of evolutionary branching trajectories

**Algorithm 1:** Numerical integration of the Canonical Equation of AD

```
/* Numerically integrate eq. X with ODE solver e.g. ode45()     */
```

**while** $t < \infty$ **do**

    $\frac{d\mu_i}{dt} = c \cdot N_i^* \cdot \frac{\partial \omega_i}{\partial \mu_i}\Big|_{\mu_i' = \mu_i}$

    **if** $\max(\frac{d\vec{\mu}}{dt}) < \epsilon$ **then**

```
        /* When gradients fall under threshold: singular strategy */
```

        **if** *any* $(\frac{d^2\vec{w}_i}{d\mu^2} > 0)$ **then**

```
            /* If any evolutionary unstable, branching occurs      */
```

            for $i$ where $\frac{d^2\vec{w}_i}{d\mu^2} > 0$ :

            $\mu_i \Longrightarrow \mu_i', \mu_i''$

            where $\mu_i' = \mu_i + e,\ \mu_i'' = \mu_i - e$

        **else**

            Terminate.

        **end**

    **end**

**end**

### S2: Coexistence metrics: niche and fitness difference

Coexistence metrics aim at quantifying the opportunity for coexistence. The structural niche difference $\Omega$ quantifies the range in intrinsic growth rates compatible with coexistence (see Fig. 1). The structural niche difference $\Omega$ quantified the size of the so-called feasibility domain (in green), i.e., the domain of intrinsic growth rate vector $\mathbf{r}$ leading to positive abundance of all species. The feasibility domain is given by a convex hull, illustrated, which is generated by the columns of the interaction matrix $\boldsymbol{\alpha}$ (all positive linear combination of the vectors forming the columns of $\boldsymbol{\alpha}$). For more details, see [5] and chapter 7.III of [3]. The size of the feasibility domain can be computed by the surface of the green triangle. Technically speaking, this formula quantifies the fraction of the feasibility domain intersecting the unit simplex. It can also be seen as the solid angle in topology $L_1$; see figure 44 (b), chapter 7.III in [3]. Following equation (7.37) in [3], $\Omega$ is computed as

$$\Omega = \frac{|\det \boldsymbol{\alpha}|}{\sum_i \alpha_{i1} \cdot \ldots \cdot \sum_i \alpha_{iS}}. \tag{S2.1}$$

In turn, the structural fitness difference, angle $\theta$ on figure 1, quantify to what extent the vector of intrinsic growth $r$ rate deviates from the centroid, $r_c$, of the feasibility domain. The centroid of the feasibility domain is given by the following vector

$$\mathbf{r}_c = \frac{1}{\sum_{i=1}^{S} \alpha_{i1}} \begin{pmatrix} \alpha_{11} \\ \alpha_{21} \\ \vdots \\ \alpha_{S1} \end{pmatrix} + \cdots + \frac{1}{\sum_{i=1}^{S} \alpha_{iS}} \begin{pmatrix} \alpha_{1S} \\ \alpha_{2S} \\ \vdots \\ \alpha_{SS}. \end{pmatrix} \tag{S2.2}$$

Then, the angle $\theta$ is simply computed using the scalar product

$$\theta = \arccos\left(\frac{\mathbf{r} \cdot \mathbf{r}_c}{||\mathbf{r}|| \cdot ||\mathbf{r}_c||}\right). \tag{S2.3}$$

Finally, the structural measure of "robustness", angle $\eta$ on figure 1, is defined as the smallest angle between the vector of intrinsic growth rates $\mathbf{r}$ and the border of the feasibility domain [4]. To compute, $\eta$ we first evaluate the smallest angle between $\mathbf{r}$ and each of the $S$ border of the feasibility domain. Then $\eta$ is given by the minimum of those $S$ angles. To do, we project orthogonally $\mathbf{r}$ on each of the $S$ border. This defines a set of $S$ vectors computed as

$$\mathbf{p}_i = \boldsymbol{\alpha}_{-i} \cdot \left(\boldsymbol{\alpha}_{-i}^t \cdot \boldsymbol{\alpha}_{-i}\right)^{-1} \cdot \boldsymbol{\alpha}_{-i}^t \cdot \mathbf{r}, \qquad i = 1, \ldots, S, \tag{S2.4}$$

where $\boldsymbol{\alpha}_{-i}$ is the $\boldsymbol{\alpha}$ form which column $i$ has been removed. Then the set of smallest angles $\eta_i$ between $\mathbf{r}$ and each of the $S$ border is computed using the scalar product

$$\eta_i = \arccos\left(\frac{\mathbf{r} \cdot \mathbf{p}_i}{\|\mathbf{r}\| \cdot \|\mathbf{p}_i\|}\right). \tag{S2.5}$$

Finally, $\eta$ is the minimum of those $\eta_i$, i.e.,

$$\eta = \min(\eta_1, \ldots, \eta_S). \tag{S2.6}$$

### S3: Analytic derivation of the branching conditions for 1 morph

#### *Location of the singular strategy for $S = 1$*

An evolutionarily singular strategy is a point where the selection gradient vanishes. We can find the location of the monomorphic singular strategy for our model analytically. Recall that

$$r(\mu) = f(\mu) - m = f_{\max} e^{-\frac{(\mu_R - \mu)^2}{2\sigma_R^2}} - m$$

$$\alpha(\mu_1, \mu_2) = \alpha_{\max} e^{-\frac{(\mu_1 - \mu_2)^2}{2\sigma_\alpha^2}}$$

The invasion fitness (Equ. 4) in the monomorphic case simplifies to :

$$w(\mu_m, \mu_r) = r(\mu_m) - \alpha(\mu_m, \mu_r) \frac{r(\mu_r)}{\alpha(\mu_r, \mu_r)}$$

and the selection gradient is given by its derivative:

$$\frac{\partial w}{\partial \mu_m} = \frac{dr(\mu_m)}{d\mu_m} - \frac{\partial \alpha(\mu_m, \mu_r)}{\partial \mu_m} \frac{r(\mu_r)}{\alpha(\mu_r, \mu_r)}$$

Now substituting the above expressions for $r$ and $\alpha$ and evaluating at $\mu = \mu_m = \mu_r$ we get

$$\frac{\partial w}{\partial \mu_m}\Big|_{\mu_m = \mu_r} = \frac{f_{\max}(\mu_R - \mu) e^{-\frac{(\mu_R - \mu)^2}{2\sigma_R^2}}}{\sigma_R^2} = 0$$

Since $f_{\max}$ and $\sigma_R$ are strictly positive, we find that the one-morph singular strategy is located at $\mu^* = \mu_R$. Note that in this monomorphic case $\frac{\partial \alpha(\mu_m, \mu_r)}{\partial \mu_m}\Big|_{\mu_m = \mu_r} = 0$, so we have $\frac{\partial w}{\partial \mu_m} = \frac{dr(\mu_m)}{d\mu_m}$ and $\mu^* = \operatorname{argmax}_\mu r(\mu) = \operatorname{argmax}_\mu K(\mu)$, where $K(\mu) = \frac{r(\mu)}{\alpha(\mu, \mu)}$ the carrying capacity is strictly proportional to $r(\mu)$ since $\alpha(\mu, \mu) = \alpha_{\max}$. Thus, the intrinsic growth rate and the biomass is optimized at the singular strategy. This identity does not hold in higher dimensions, where $\boldsymbol{\mu}_r$ is a vector of resident morphs (see also Fig. 5) and the competition component of the gradient is non-null.

#### *Derivation of the branching condition for $S = 1$*

The condition for switching between a branching point (convergence stable, evolutionary unstable strategy) to a Continuously Stable Strategy (convergence and evolutionary stable strategy) can be derived analytically for our model in the monomorphic case. Both branching points and CSSs are points where the selection gradient becomes null. The difference is that a CSS is an uninvadable fitness maxima, while a branching point is a fitness minimum that can be invaded by adjacent strategies. Calculus tells us that the difference lies in the sign of the derivative of the fitness gradient (i.e., the second derivative of the invasion fitness). A singular strategy is evolutionarily stable if

$$\left.\frac{\partial^2 w}{\partial \mu_m^2}\right|_{\mu_m=\mu^*} < 0 \ [1]$$

The second derivative of the invasion fitness in our model is:

$$\frac{\partial^2 w}{\partial \mu_m^2} = \frac{d^2 r(\mu_m)}{d\mu_m^2} - \frac{\partial^2 \alpha(\mu_m, \mu_r)}{\partial \mu_m^2} \frac{r(\mu_r)}{\alpha(\mu_r, \mu_r)}$$

Plugging in our expressions for $r$ and $\alpha$ and evaluating at $\mu = \mu^*$:

$$\left.\frac{\partial^2 w}{\partial \mu_m^2}\right|_{\mu_m=\mu_r=\mu^*} = \frac{f_{\max} e^{-\frac{(\mu_R-\mu)^2}{2\sigma_R^2}} - m}{\sigma_\alpha^2} + f_{\max}\left(\frac{(\mu_R-\mu)^2 e^{-\frac{(\mu_R-\mu)^2}{2\sigma_R^2}}}{\sigma_R^4} - \frac{e^{-\frac{(\mu_R-\mu)^2}{2\sigma_R^2}}}{\sigma_R^2}\right)$$

$$= \frac{\frac{f_{\max} e^{-\frac{(\mu_R-\mu)^2}{2\sigma_R^2}}\left(-\sigma^2\sigma_R^2+\sigma_R^4+\sigma_\alpha^2(\mu_R-\mu)^2\right)}{\sigma_R^4} - m}{\sigma_\alpha^2}$$

We have shown earlier that at the singular strategy $\mu^* = \mu_R$, so we can simplify further:

$$\frac{\frac{f_{\max}\left(\sigma_R^4-\sigma_\alpha^2\sigma_R^2\right)}{\sigma_R^4} - m}{\sigma_\alpha^2} = \frac{f_{\max} - m}{\sigma_\alpha^2} - \frac{a}{\sigma_R^2}$$

$$= \frac{\sigma_R^2(f_{\max} - m) - \sigma_\alpha^2 f_{\max}}{\sigma_\alpha^2\sigma_R^2} < 0$$

Since $f_{\max}$, $\sigma_R$, and $\sigma_\alpha$ all must be greater than zero, the condition for evolutionary stability is

$$\sigma_R^2(f_{\max} - m) < \sigma_\alpha^2 f_{\max}$$

and for $m << f_{\max}$ we retrieve the condition of Doebeli and Dieckmann [2], equation 13, i.e., the monomorphic singular strategy is a branching point for $\sigma_R > \sigma$.

#### *Branching points and niche neutrality continuum*

In our model, because of the non-independence of coexistence metrics, there is no niche-neutrality continuum (*sensu* Song et al. [6]). That is, zones of neutral coexistence and niche-driven coexistence are separated by an unfeasible region: two species indefinitely close cannot generally coexist, unless they are identical. If not, they must have a certain niche difference given by a minimum trait difference $\epsilon = \mu_1 - \mu_2 > \epsilon_{min}$ function of $\delta := \mu_R - \mu_1$ , $\sigma_\alpha$, and $\sigma_R$, to stably coexist. For the two species Lotka-Volterra system, the equilibrium abundances can be found by solving the following system of equations for the non-trivial solution $N_1^*, N_2^* > 0$

$$\begin{cases} \frac{dN_1}{dt} = N_1 \cdot (r_1 - \alpha_{11}N_1 - \alpha_{12}N_2) \\ \frac{dN_2}{dt} = N_2 \cdot (r_2 - \alpha_{21}N_1 - \alpha_{22}N_2) \end{cases}$$

which gives:

$$N_1^* = \frac{\alpha_{22}r_1 - \alpha_{12}r_2}{\alpha_{11}\alpha_{22} - \alpha_{12}\alpha_{21}},\ N_2^* = \frac{\alpha_{11}r_2 - \alpha_{21}r_1}{\alpha_{11}\alpha_{22} - \alpha_{12}\alpha_{21}}$$

Since under our model competition is always greater than zero for $\Delta\mu < \infty$, the denominator is always positive and the signs of the abundances are given by the numerator. Now, let's assume two morphs characterized by close trait values $\mu_1$ and $\mu_2 = \mu_1 + \epsilon$. Assuming morph 1 has the larger growth rate, its abundance will always be positive since $r_1 > r_2$ and $\alpha_{22} > \alpha_{12}$. For the sign of $N_2^*$ on the other hand, we have (assuming no density-independent mortality):

$$N_2^* \propto \alpha_{11} r_2 - \alpha_{21} r_1 = \alpha_{\max} e^{-\frac{(\mu_1-\mu_1)^2}{2\sigma_\alpha^2}} f_{\max} e^{-\frac{(\mu_R-\mu_2)^2}{2\sigma_R^2}} - \alpha_{\max} e^{-\frac{(\mu_1-\mu_2)^2}{2\sigma_\alpha^2}} f_{\max} e^{-\frac{(\mu_R-\mu_1)^2}{2\sigma_R^2}} \quad \text{(S3.1)}$$

Setting $\delta := \mu_R - \mu_1$

$$\begin{aligned} N_2^* &\propto f_{\max}\, \alpha_{\max}\, e^{-\frac{(\delta+\epsilon)^2}{2\sigma_R^2}} - \alpha_{\max} e^{-\frac{\epsilon^2}{2\sigma_\alpha^2}} f_{\max} e^{-\frac{\delta^2}{2\sigma_R^2}} \\ &= f_{\max}\, \alpha_{\max} \left( e^{-\frac{(\delta+\epsilon)^2}{2\sigma_R^2}} - e^{-\frac{\epsilon^2}{2\sigma_\alpha^2} - \frac{\delta^2}{2\sigma_R^2}} \right) \end{aligned}$$

$N_2^*$ is then positive if

$$-\frac{(\delta+\epsilon)^2}{2\sigma_R^2} > -\frac{\epsilon^2}{2\sigma_\alpha^2} - \frac{\delta^2}{2\sigma_R^2}$$

$$\epsilon > \frac{\delta\, \sigma_\alpha^2}{\sigma_R^2 - \sigma_\alpha^2}$$

So, morphs 1 and 2 need a minimum spacing for both to have positive abundances. But note the special case where $\mu_R = \mu_1$ (first singular strategy), equation S3.1 reduces to:

$$N_2^* \propto f_{\max}\, \alpha_{\max}\, e^{-\frac{\epsilon^2}{2\sigma_R^2}} - \alpha_{\max}\, e^{-\frac{\epsilon^2}{2\sigma^2}} f_{\max} \propto \frac{1}{\sigma_\alpha^2} - \frac{1}{\sigma_R^2}$$

so that for any $\epsilon$, morphs can coexist if $\sigma_R > \sigma_\alpha$. This is the branching condition derived above for $m = 0$. In fact, we have shown that branching points are the only point where two arbitrarily close species can coexist for $\epsilon \to 0$, i.e., the only place where there is a niche-neutrality continuum, which permits the splitting of one identical morph (neutral coexistence) into two polymorphic lineages (niche-driven coexistence). Which makes sense since branching points are by definition strategies that are invadable from either side.

### S4: Effect of resource width on ESC richness

As is expected, a larger resource space allows the emergence of a greater number of phenotypes (Figure S4.1A). Moreover, for a given number of phenotypes $S$, the standardized niche difference at the evolutionary singular strategy increases with resource width, as there is then more room for niche differentiation. Vice-versa, Panel B illustrates the effect of varying the strength of similarity dependence in competition by changing niche widths $\sigma_\alpha$ of phenotypes (for a fixed resource width). For smaller niches, more phenotypes can be packed at the ESC, while for a given number of phenotypes, narrow niches allow a larger domain of feasibility $\hat{\Omega}$.

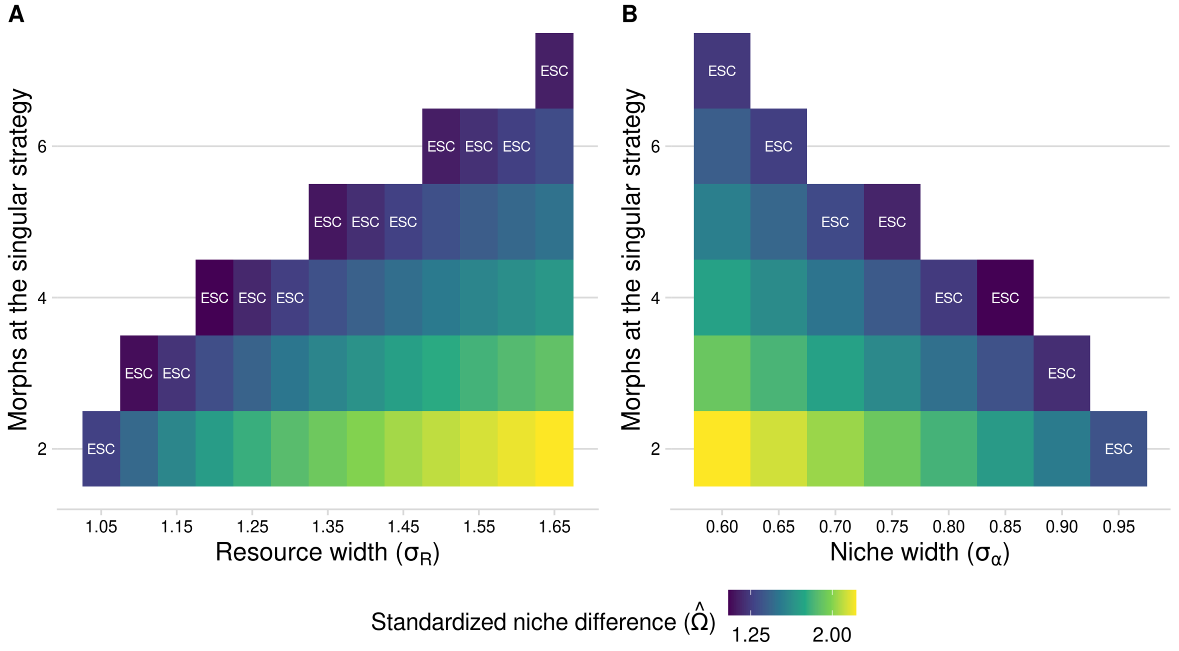


Figure S4.1: **Number of phenotypes at the ESC and at evolutionary branching points.** Panel **A** shows, for a fixed niche width $\sigma_\alpha = 1$, how varying the resource with $\sigma_R$ modulates the number of consecutive branching points and, therefore, the number of phenotypes at the ESC. Cases marked with "ESC" represent evolutionary endpoints, while all the others are invasible singular strategies (branching points). The color represents the level of standardized niche difference at the evolutionary singular strategy. Making the resource space larger, for a fixed niche width, results in greater diversity at the ESC. Moreover, for a given number of phenotypes at the evolutionary singular strategy, the structural niche difference increases with resource width. Conversely, panel **B** shows the effect of changing the niche width $\sigma_\alpha$ for a fixed resource width $\sigma_R = 1$: the smaller the niche width, the more phenotypes can fit on a given resource axis.

### S5: Relative yield measures along evolutionary time

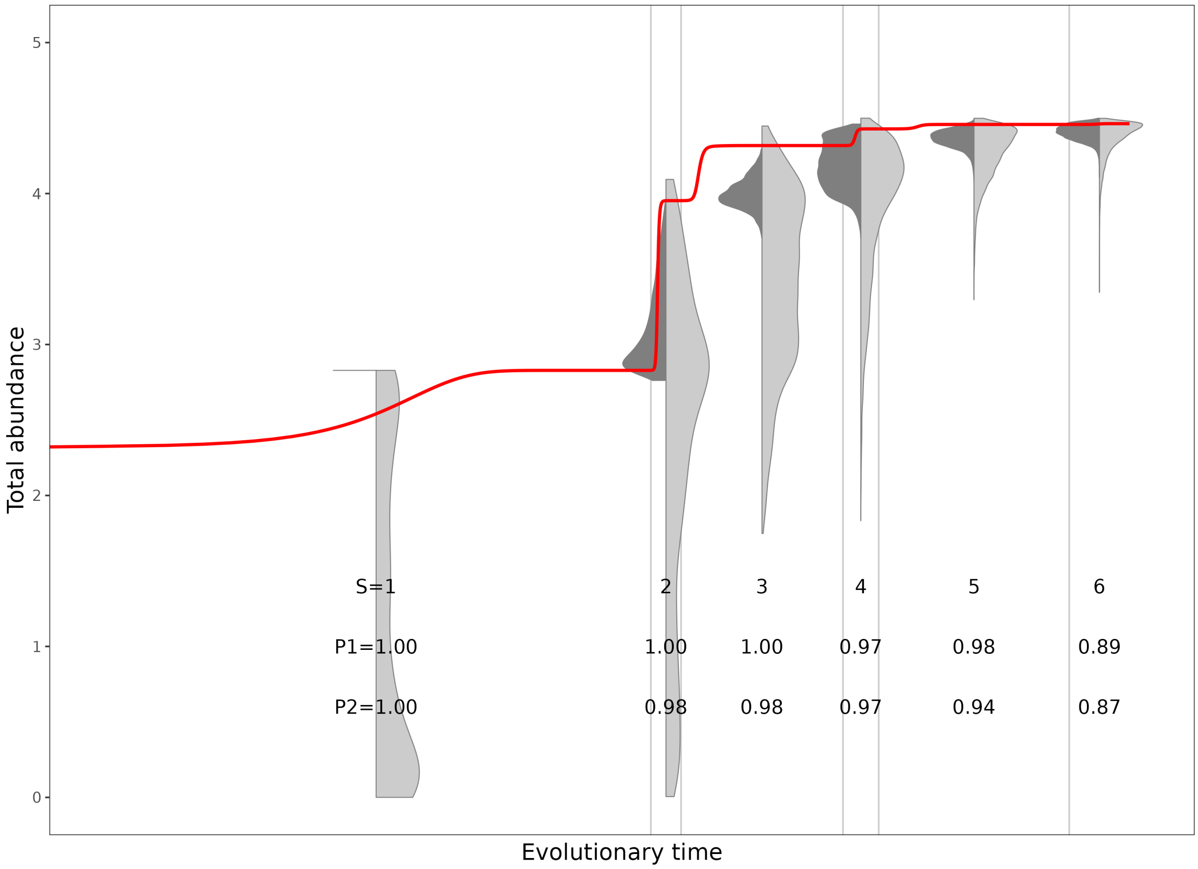


Figure S5.1: **Productivity along evolutionary trajectories.** Evolution of both productivity metrics. Here the numbers $P1$ and $P2$ in the second row of text indicate the percentage of randomized communities that are less productive than the ESC. For S=1, biomass production is optimized at the ESC. For S=6, the ESC is in the 80th percentile of the most productive communities. Point clouds represent randomization in the narrow (1st rule; dark gray) or the full range (2nd rule; light gray).

#### S6: Supplementary figures for different resource width ($\sigma_R$)

*Resource width* $\sigma_R = 1.1$

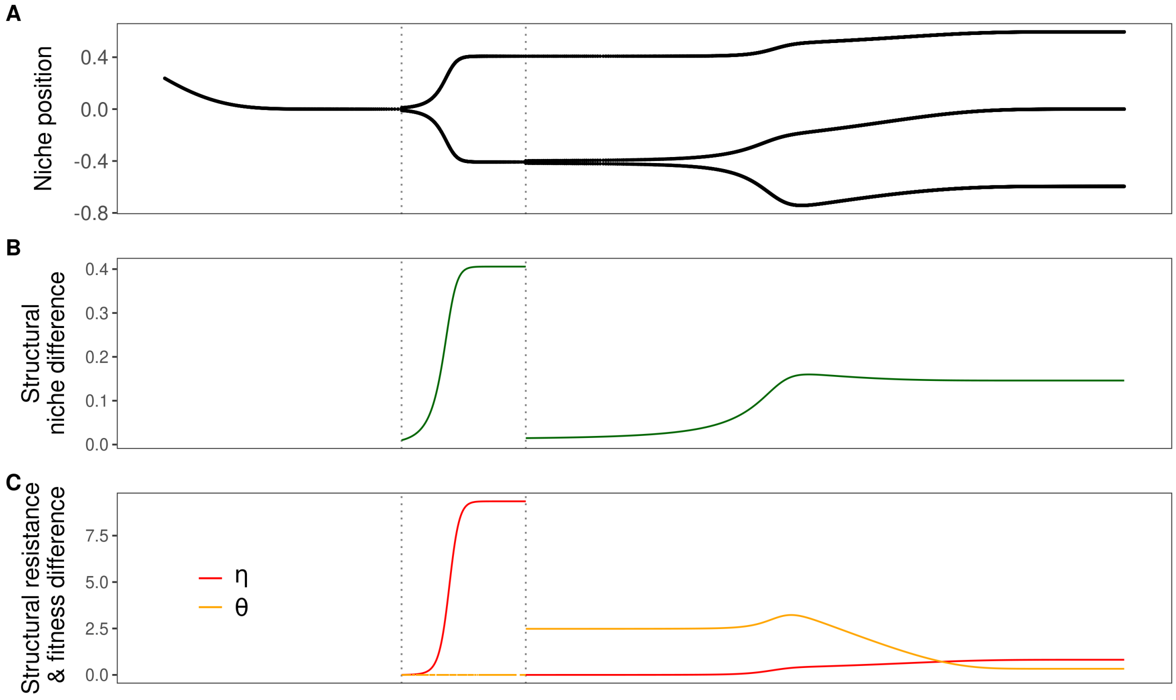


Figure S6.1: **Figure 2 for $\sigma_R$ = 1.1 (m = 0.01)**.

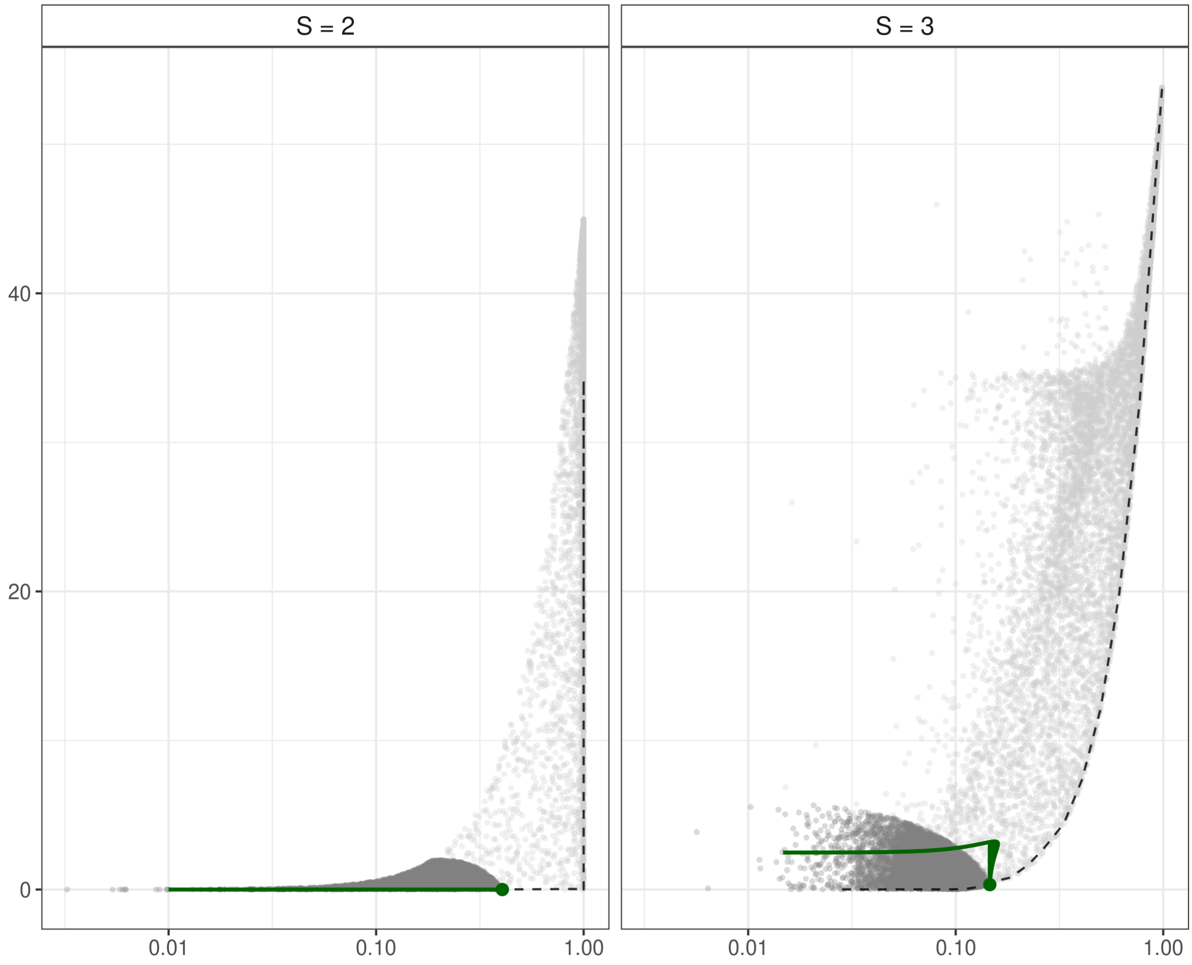


Figure S6.2: **Figure 4 for $\sigma_R$ = 1.1 (m = 0.01)**.

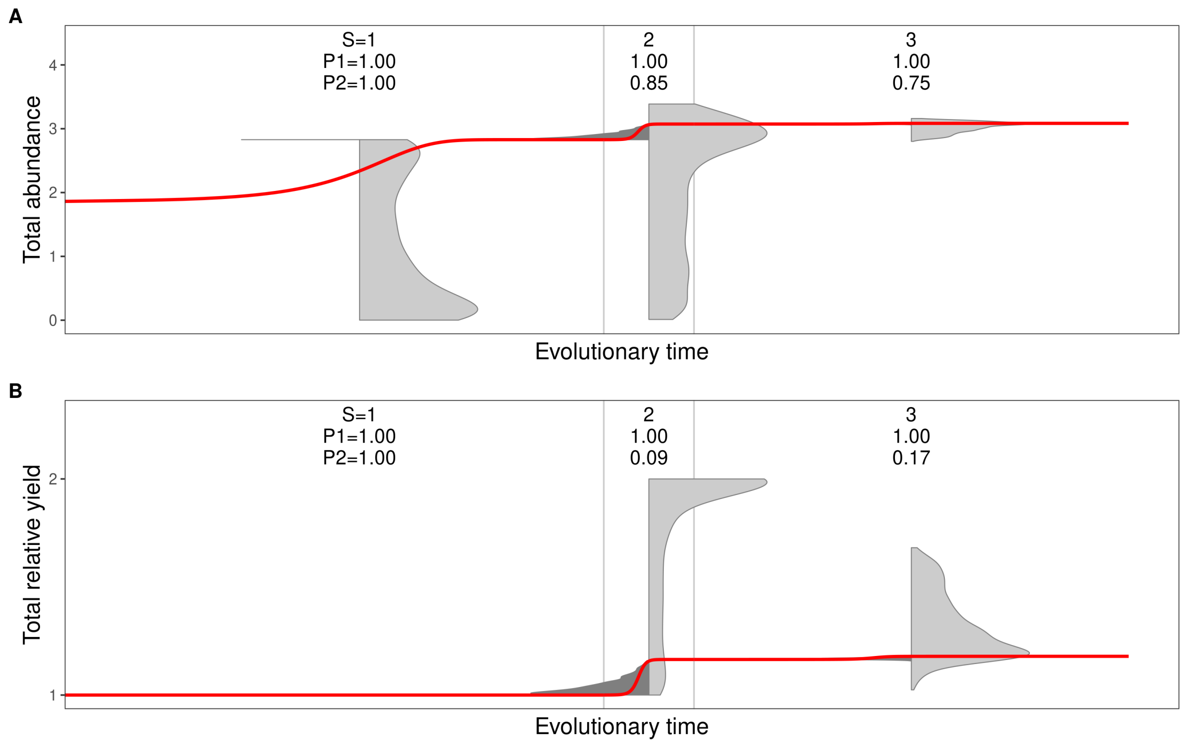


Figure S6.3: **Figure 5 for $\sigma_R = 1.1$ (m = 0.01)**.

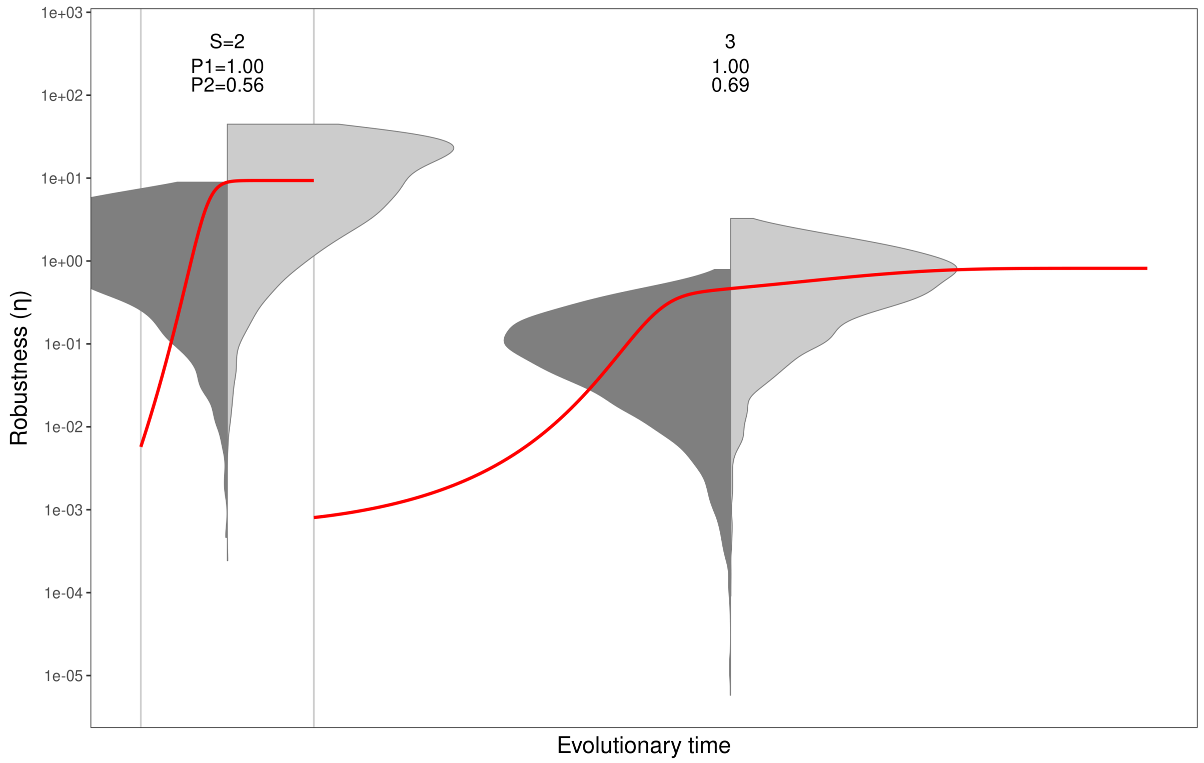


Figure S6.4: **Figure 6 for $\sigma_R = 1.1$ (m = 0.01)**.

80

#### *Resource width* $\sigma_R = 1.2$

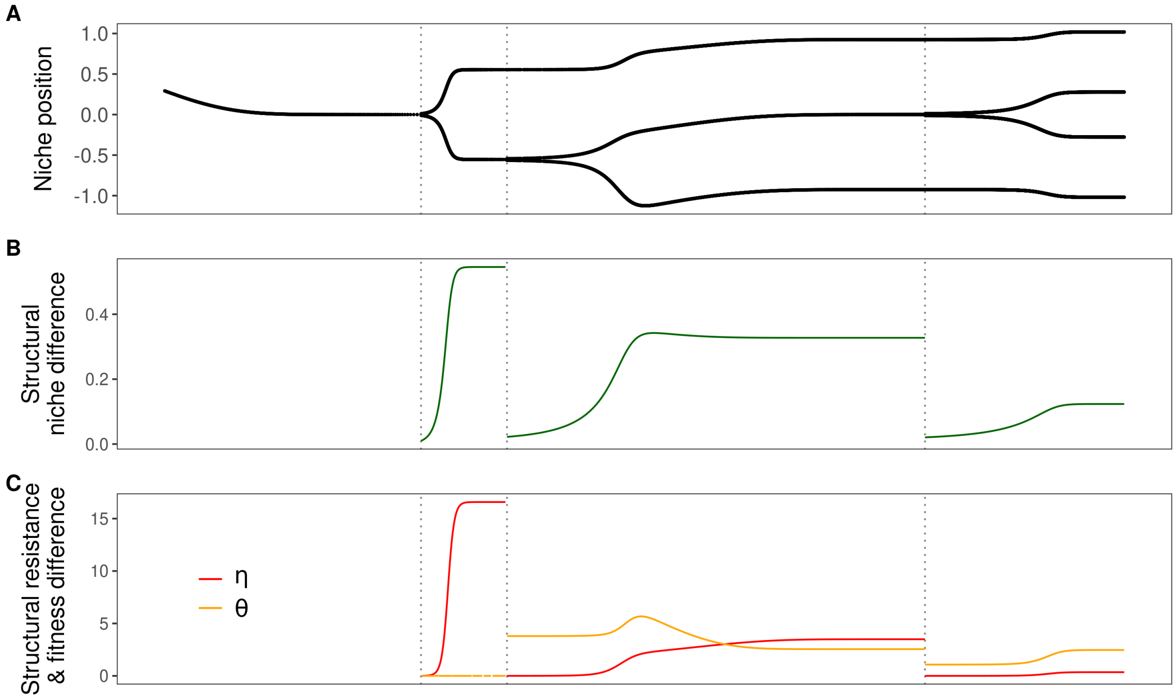


Figure S6.5: **Figure 2 for** $\sigma_R$ **= 1.2 (m = 0.01)**.

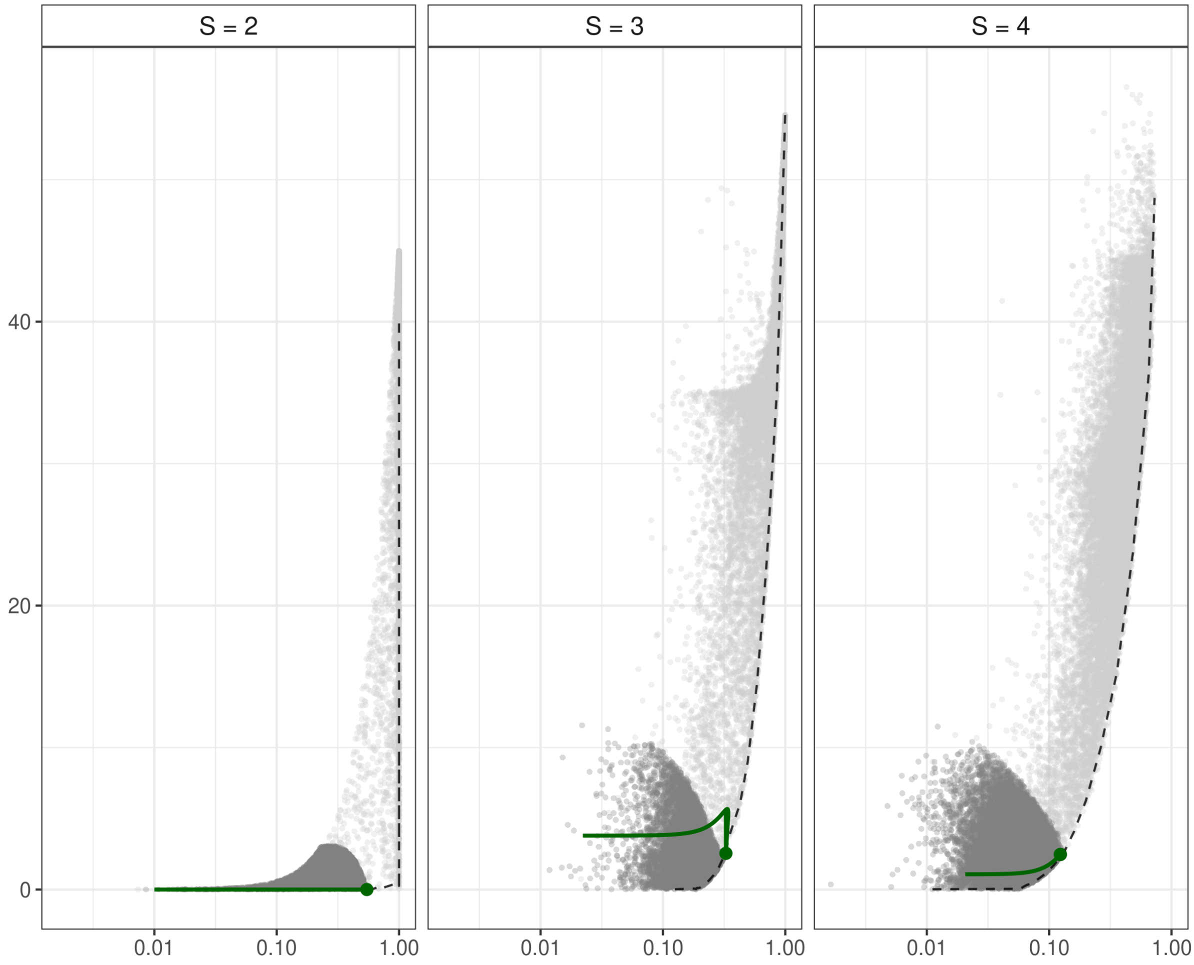


Figure S6.6: **Figure 4 for $\sigma_R$ = 1.2 (m = 0.01)**.

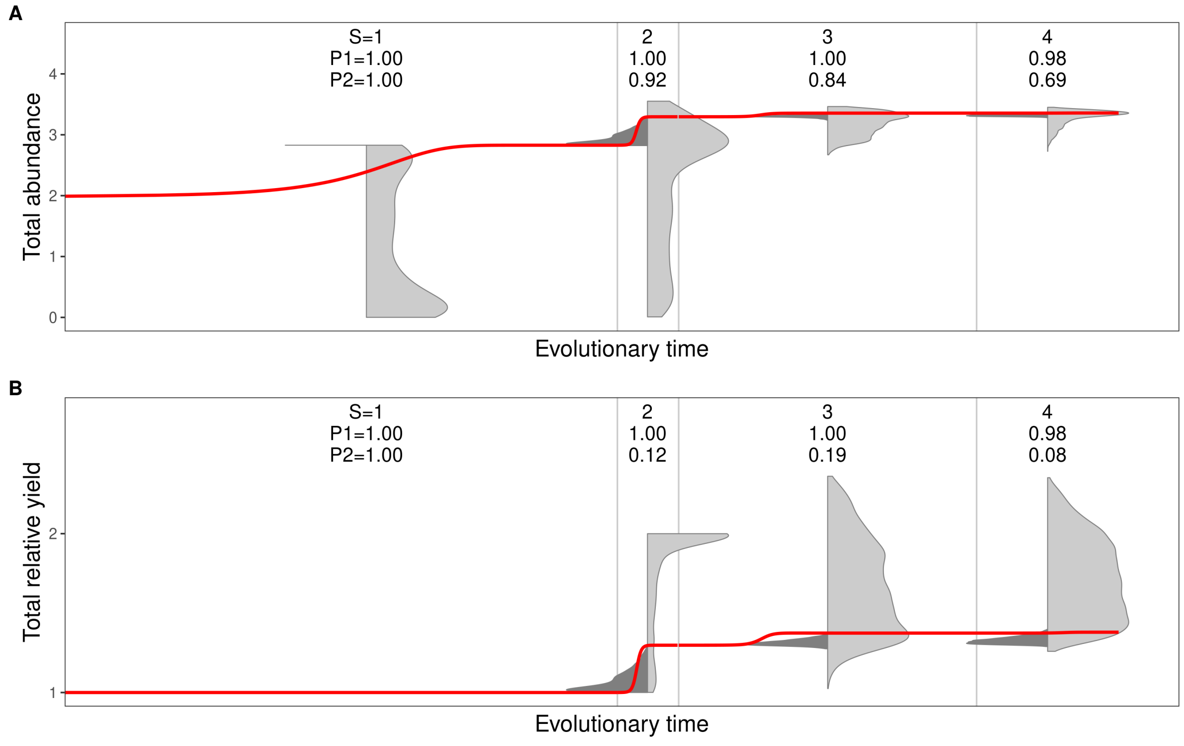


Figure S6.7: **Figure 5 for $\sigma_R = 1.2$ (m = 0.01)**.

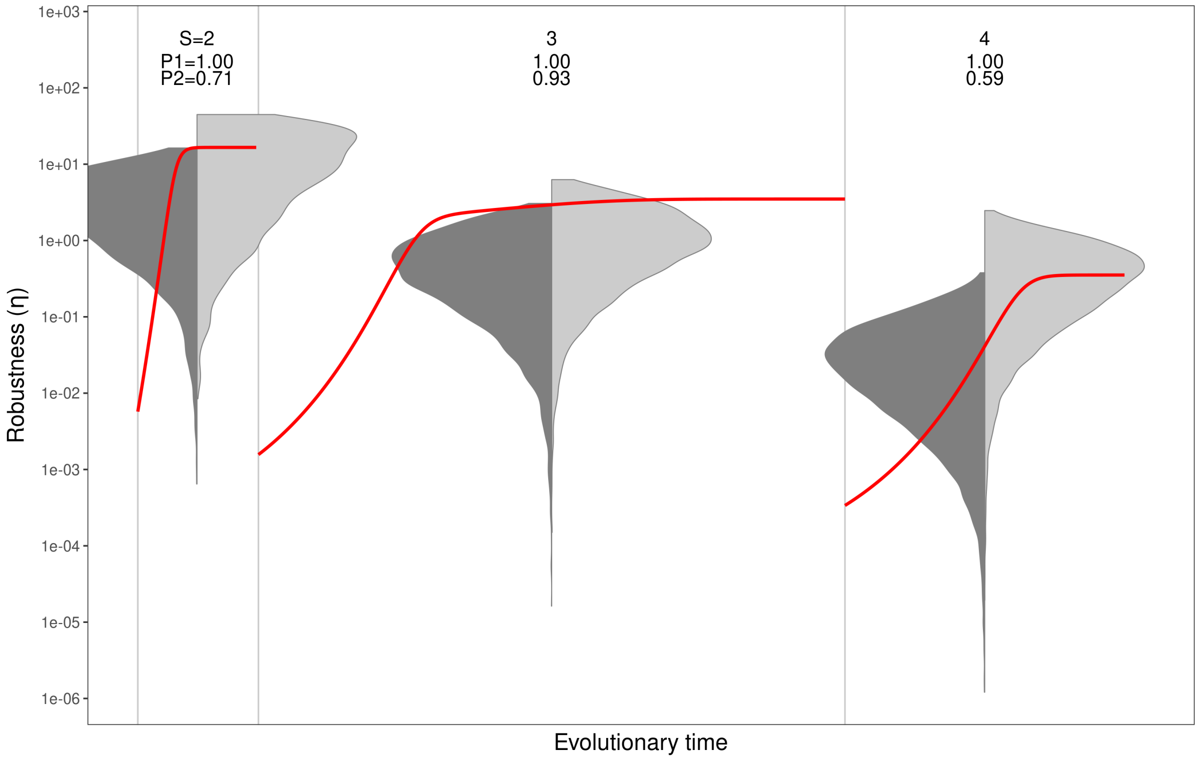


Figure S6.8: **Figure 6 for $\sigma_R = 1.2$ (m = 0.01)**.

81 *Resource width* $\sigma_R = 1.3$

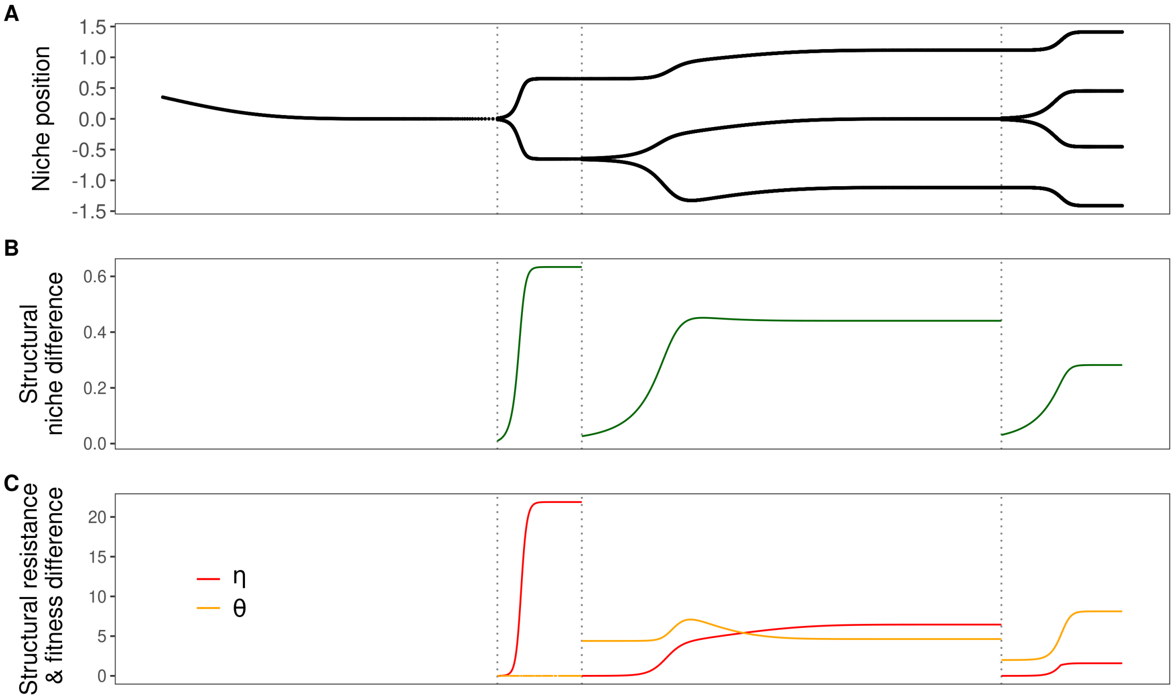


Figure S6.9: **Figure 2 for** $\boldsymbol{\sigma_R = 1.3}$ **(m = 0.01)**.

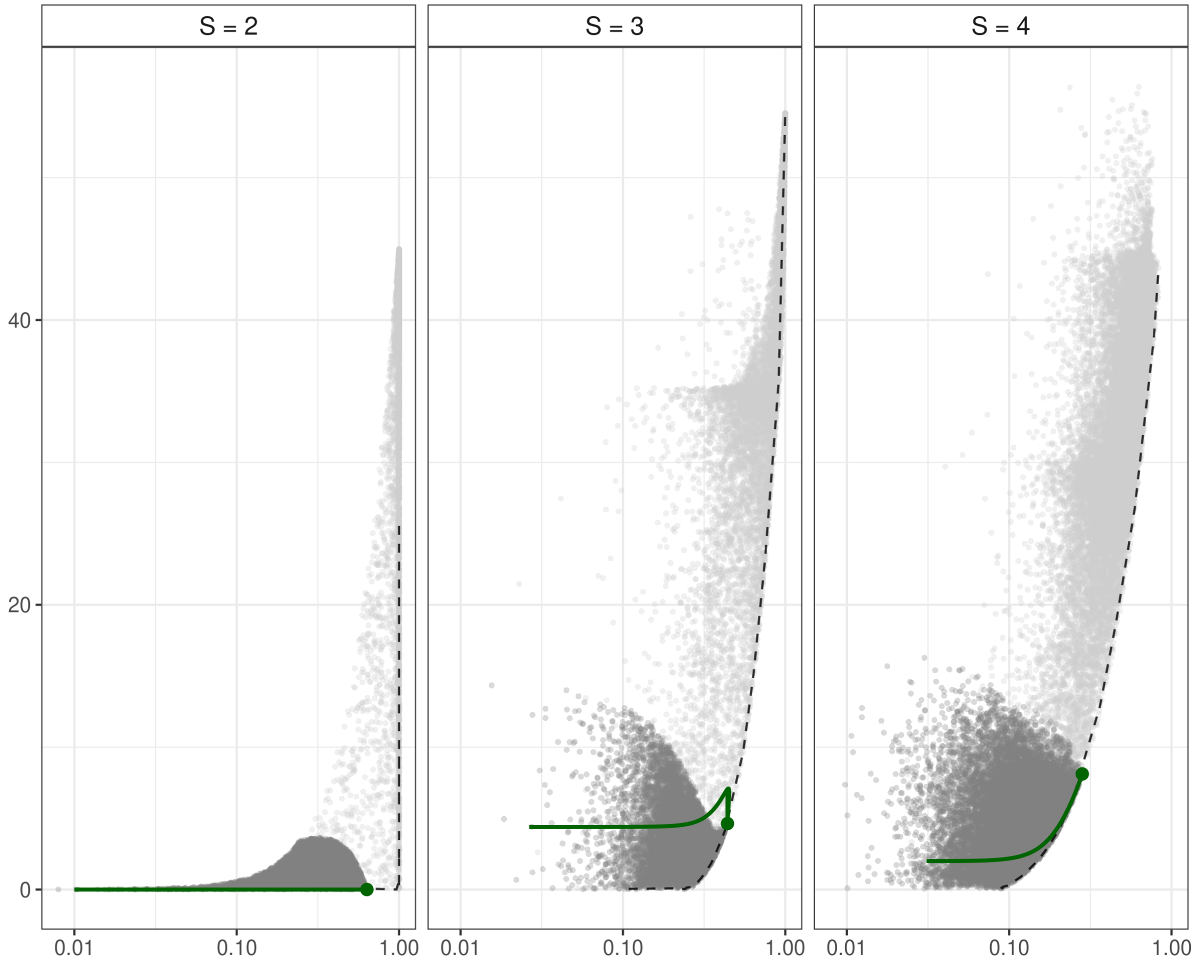


Figure S6.10: **Figure 4 for $\sigma_R$ = 1.3 (m = 0.01)**.

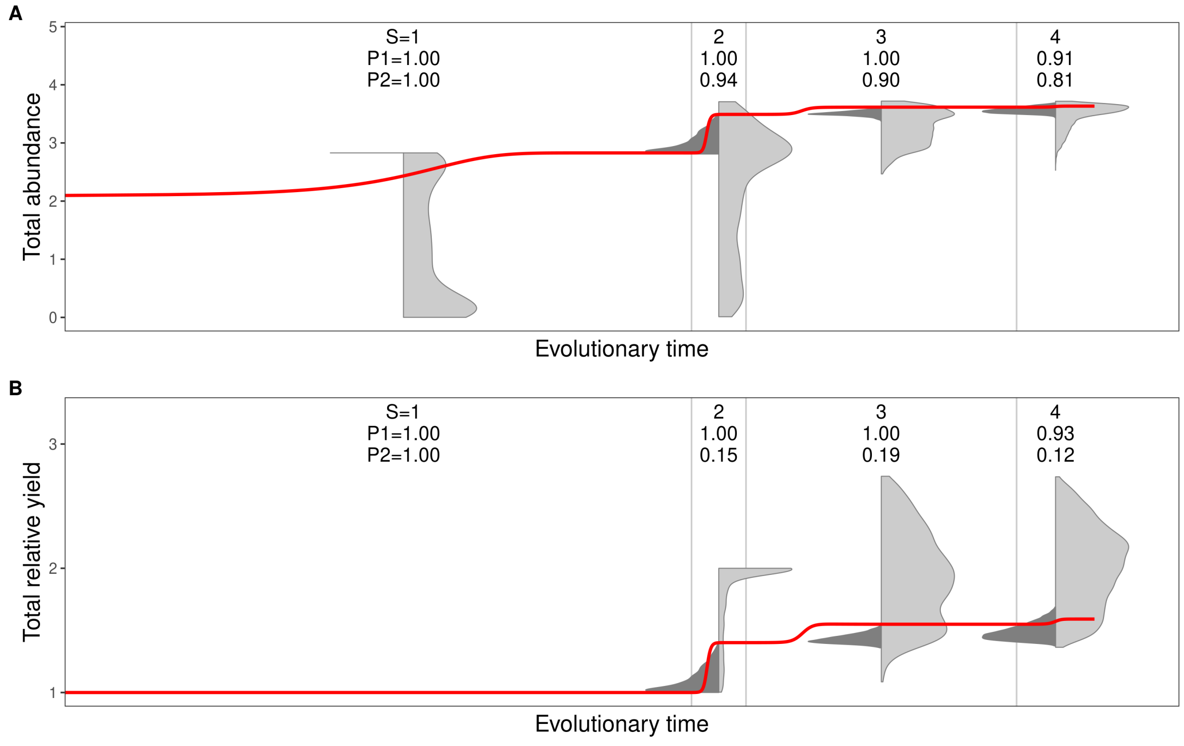


Figure S6.11: **Figure 5 for $\sigma_R = 1.3$ (m = 0.01)**.

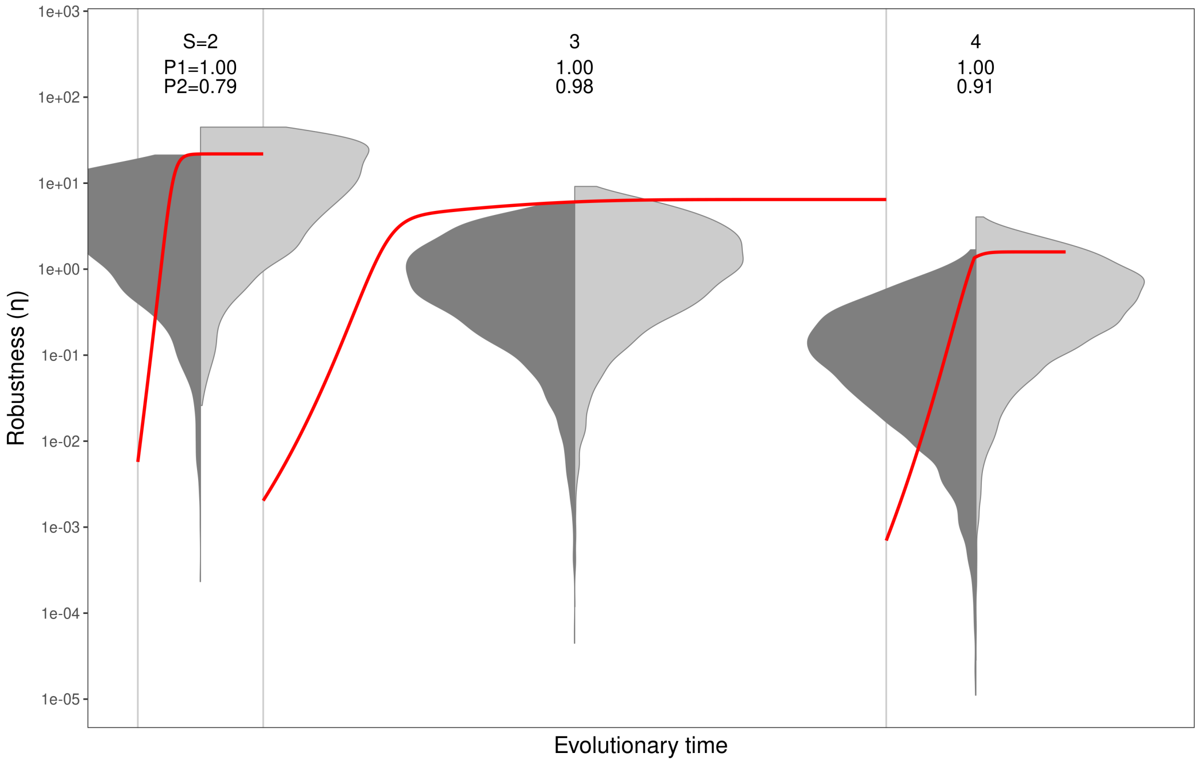


Figure S6.12: **Figure 6 for $\sigma_R = 1.3$ (m = 0.01)**.

### *Resource width* $\sigma_R = 1.4$

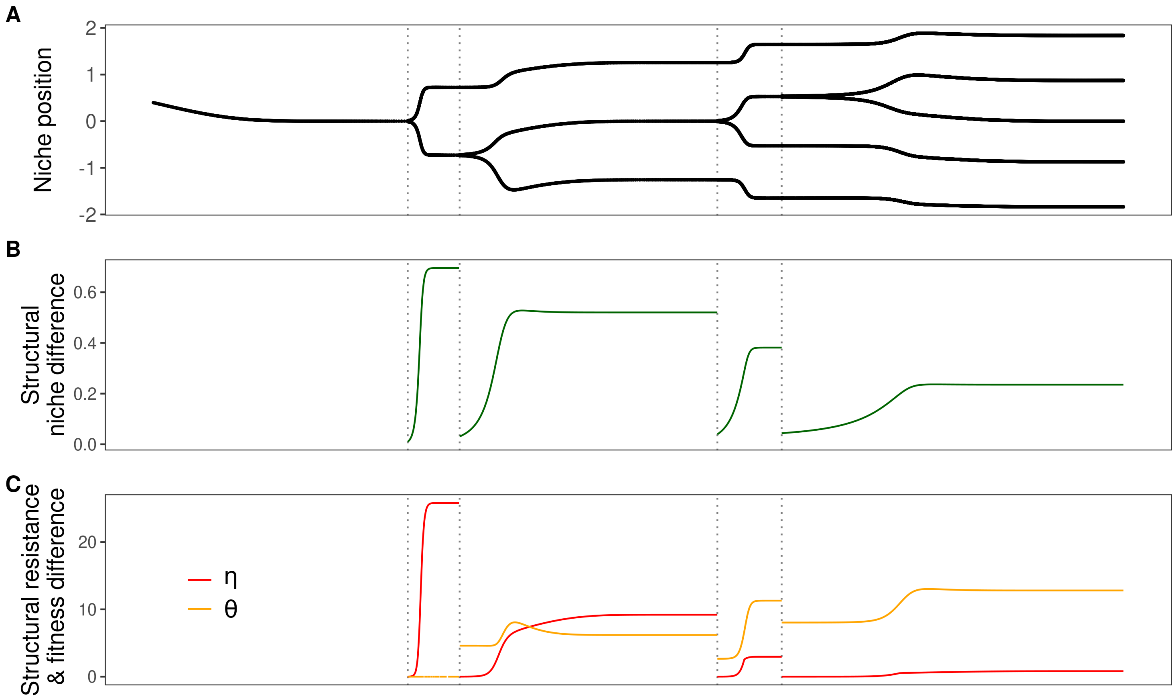


Figure S6.13: **Figure 2 for** $\sigma_R$ **= 1.4 (m = 0.01)**.

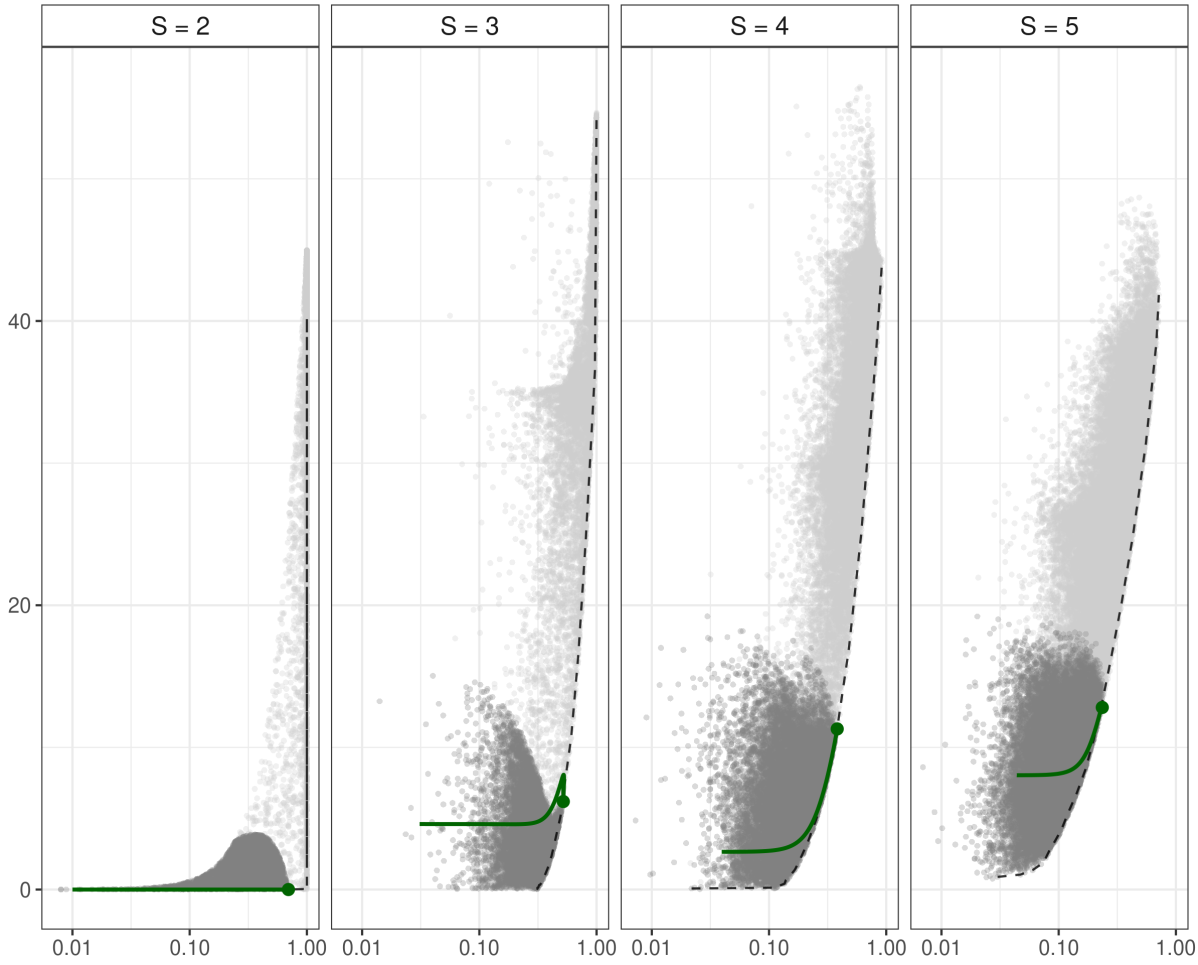


Figure S6.14: **Figure 4 for $\sigma_R$ = 1.4 (m = 0.01)**.

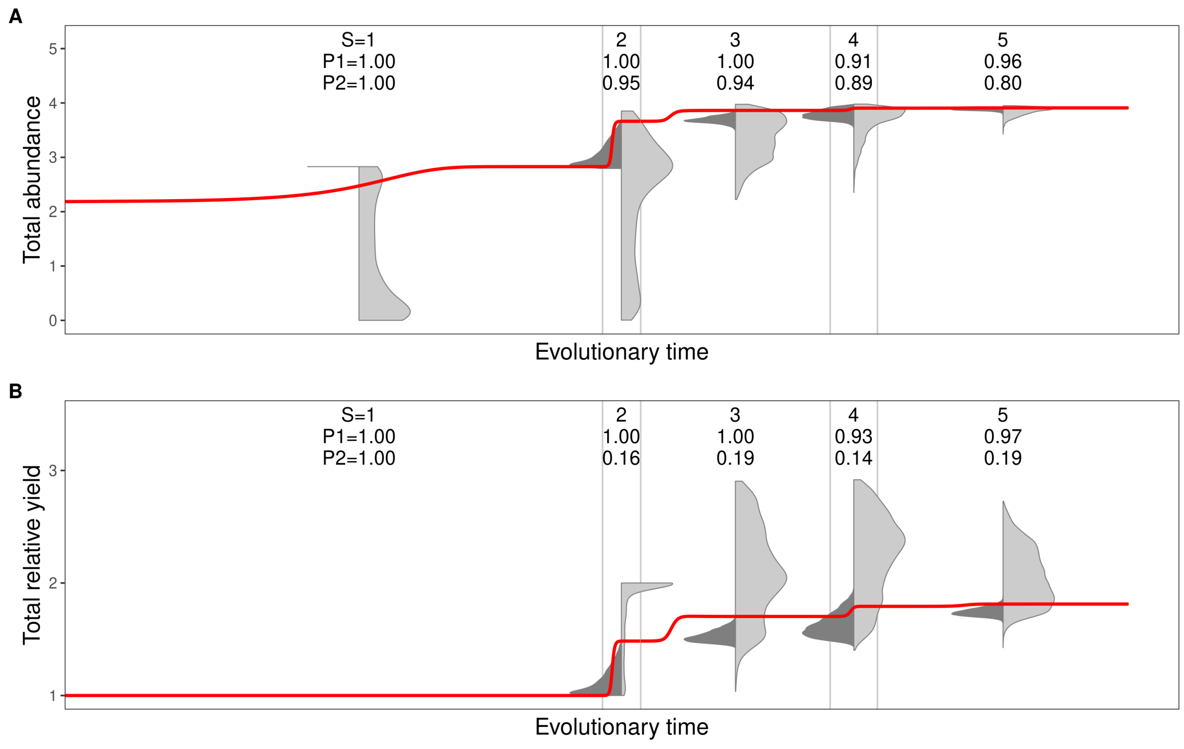


Figure S6.15: **Figure 5 for $\sigma_R$ = 1.4 (m = 0.01).**

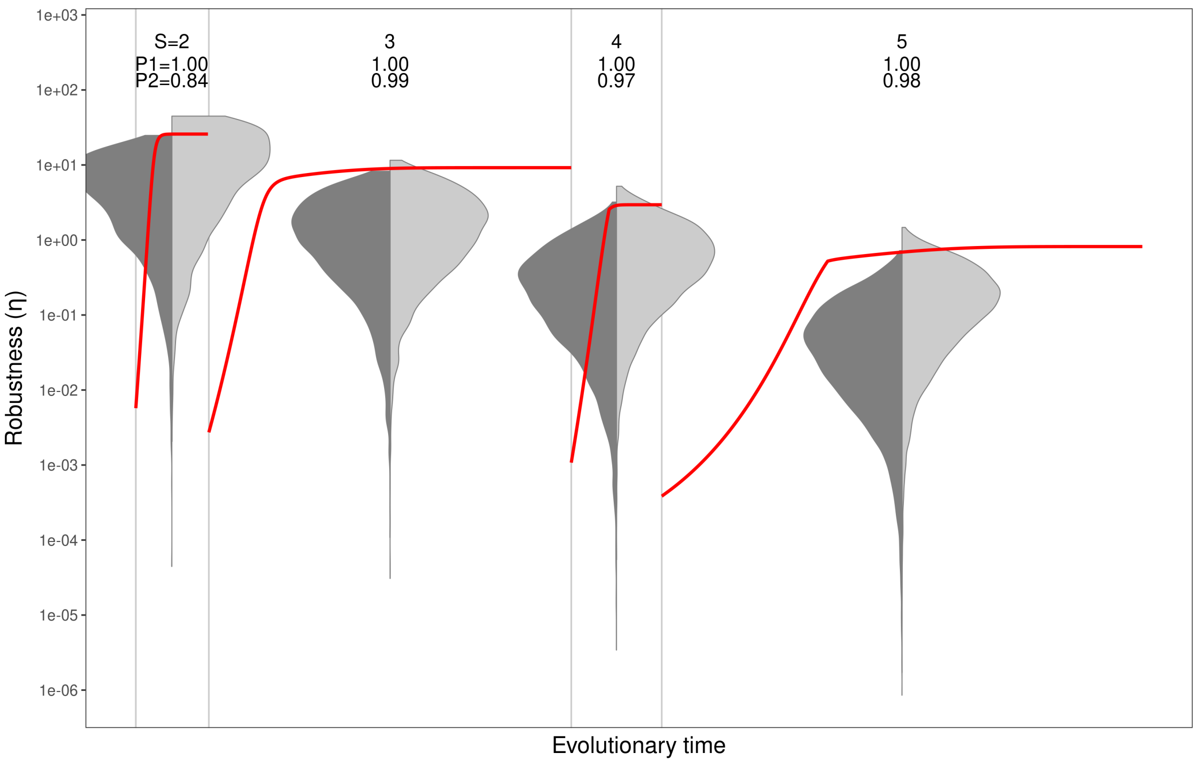


Figure S6.16: **Figure 6 for $\sigma_R$ = 1.4 (m = 0.01).**

### *Resource width* $\sigma_R = 1.5$

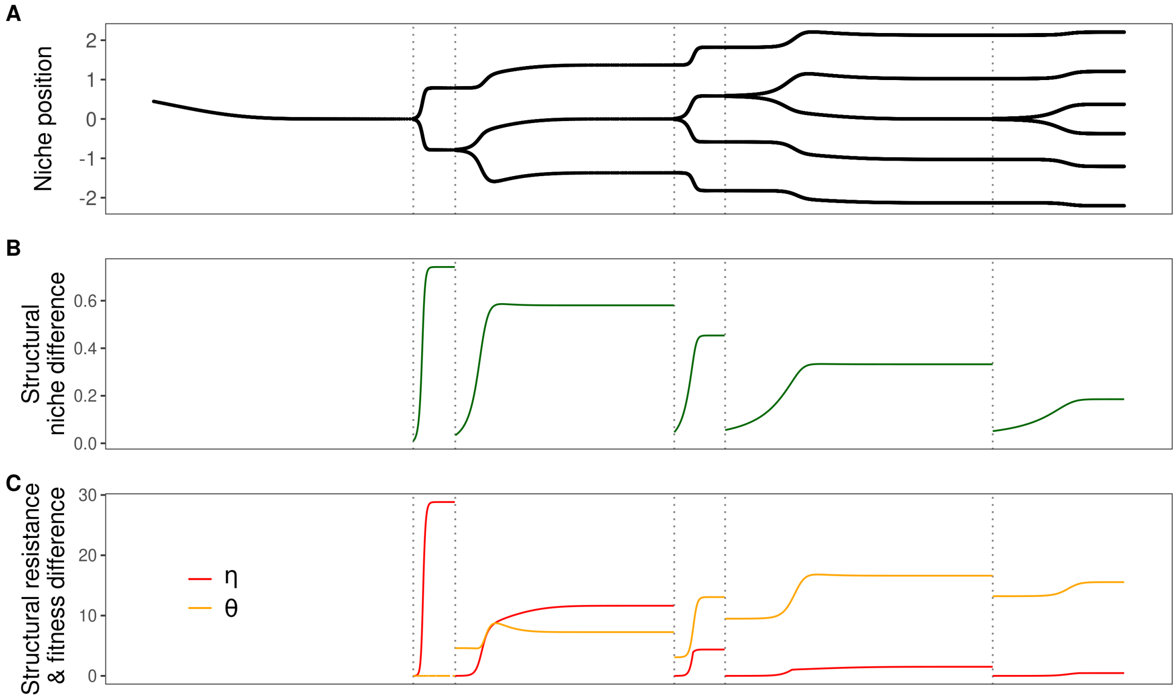


Figure S6.17: **Figure 2 for $\sigma_R$ = 1.5 (m = 0.01)**.

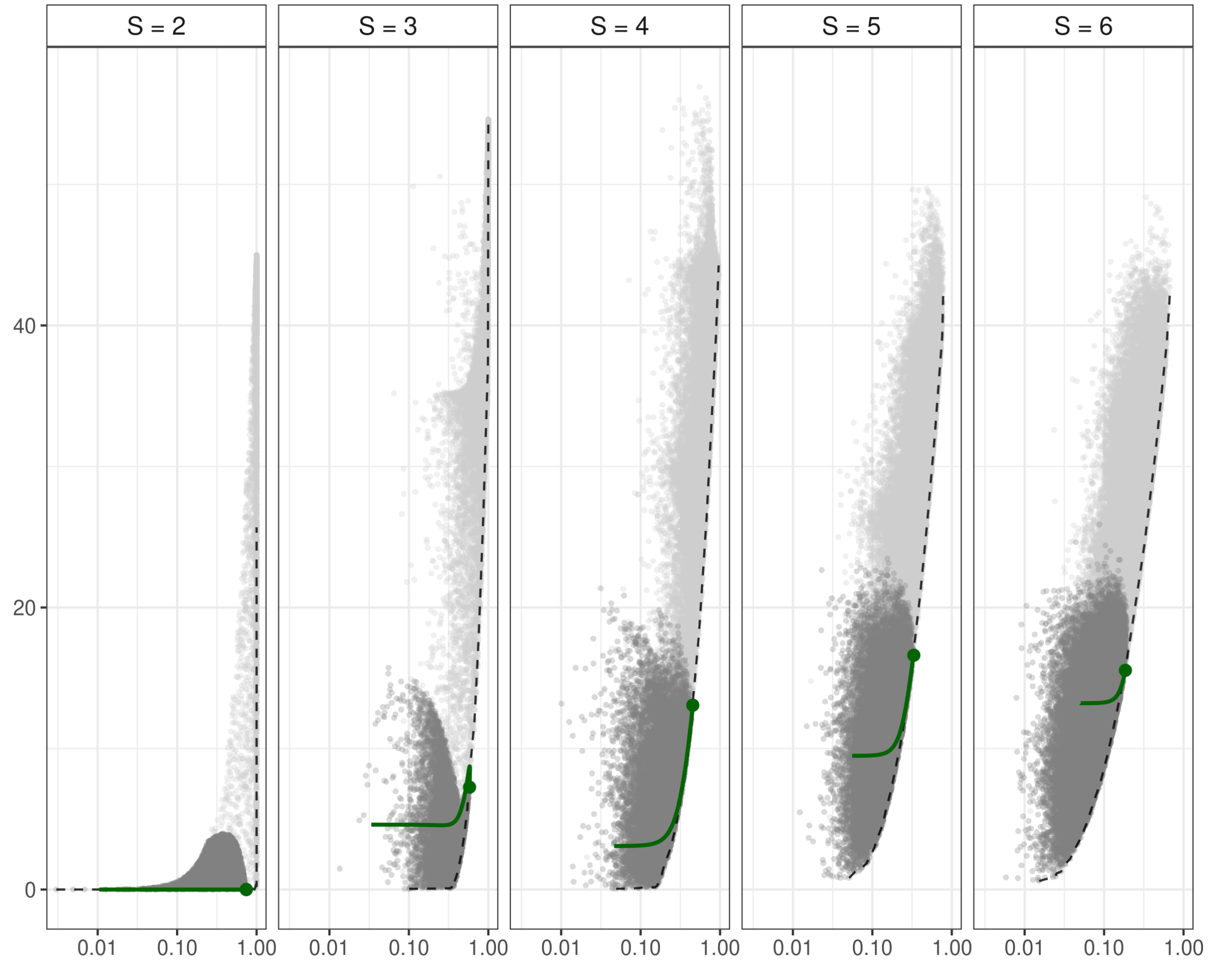


Figure S6.18: **Figure 4 for $\sigma_R = 1.5$ (m = 0.01)**.

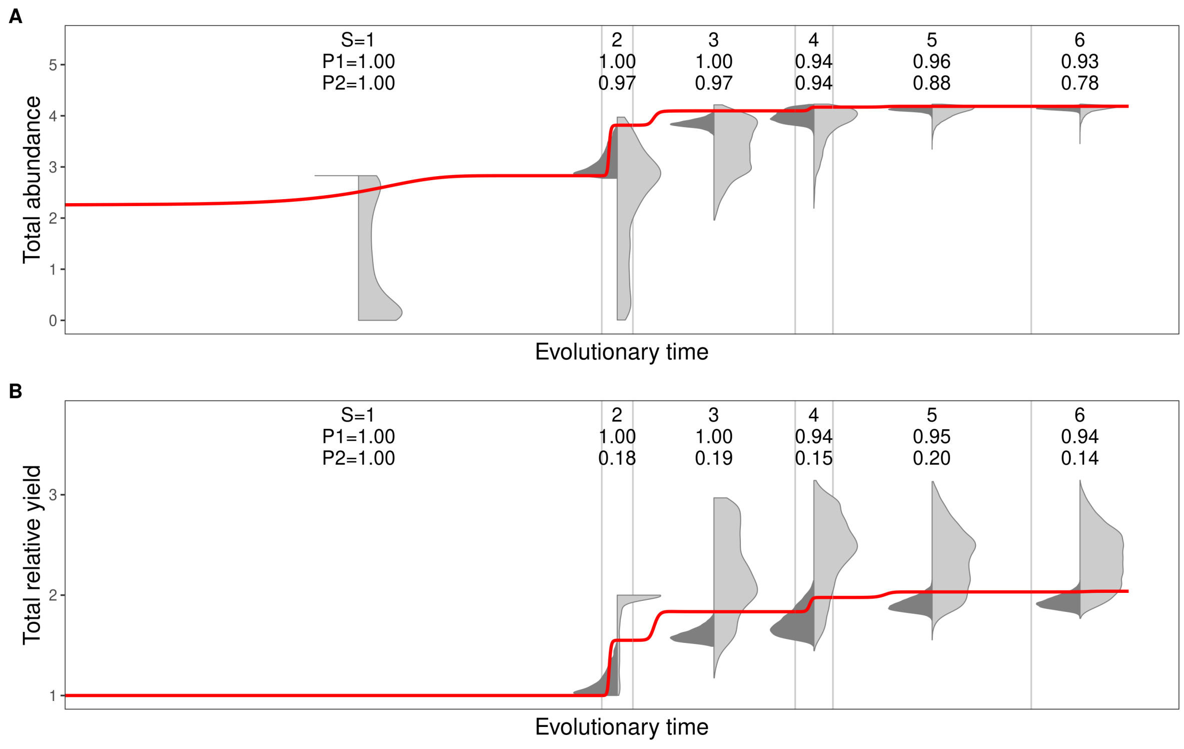


Figure S6.19: **Figure 5 for $\sigma_R = 1.5$ (m = 0.01)**.

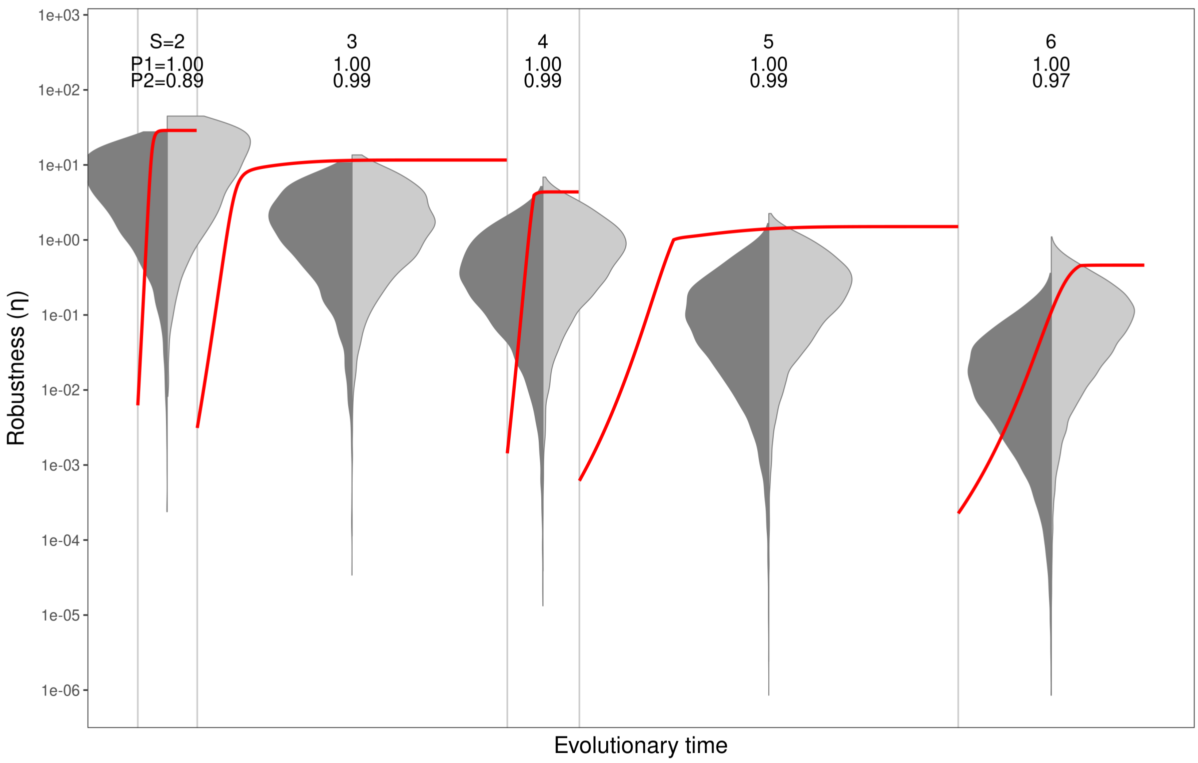


Figure S6.20: **Figure 6 for $\sigma_R = 1.5$ (m = 0.01)**.

### *Resource width* $\sigma_R = 1.6$

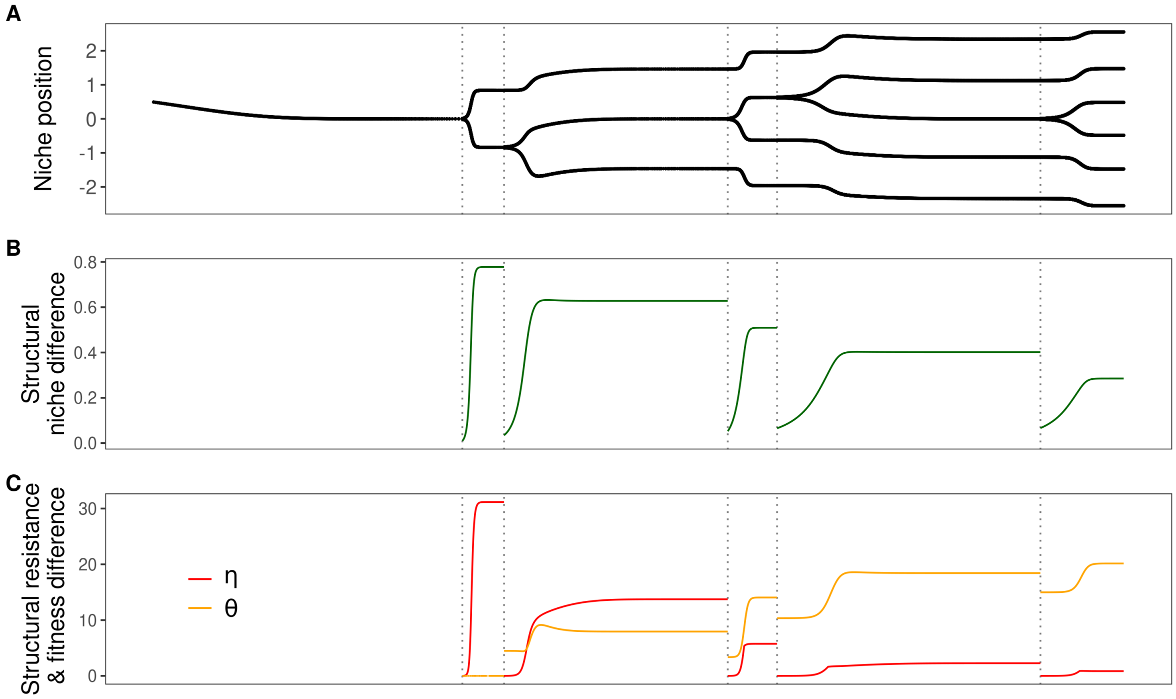


Figure S6.21: **Figure 2 for $\sigma_R$ = 1.6 (m = 0.01)**.

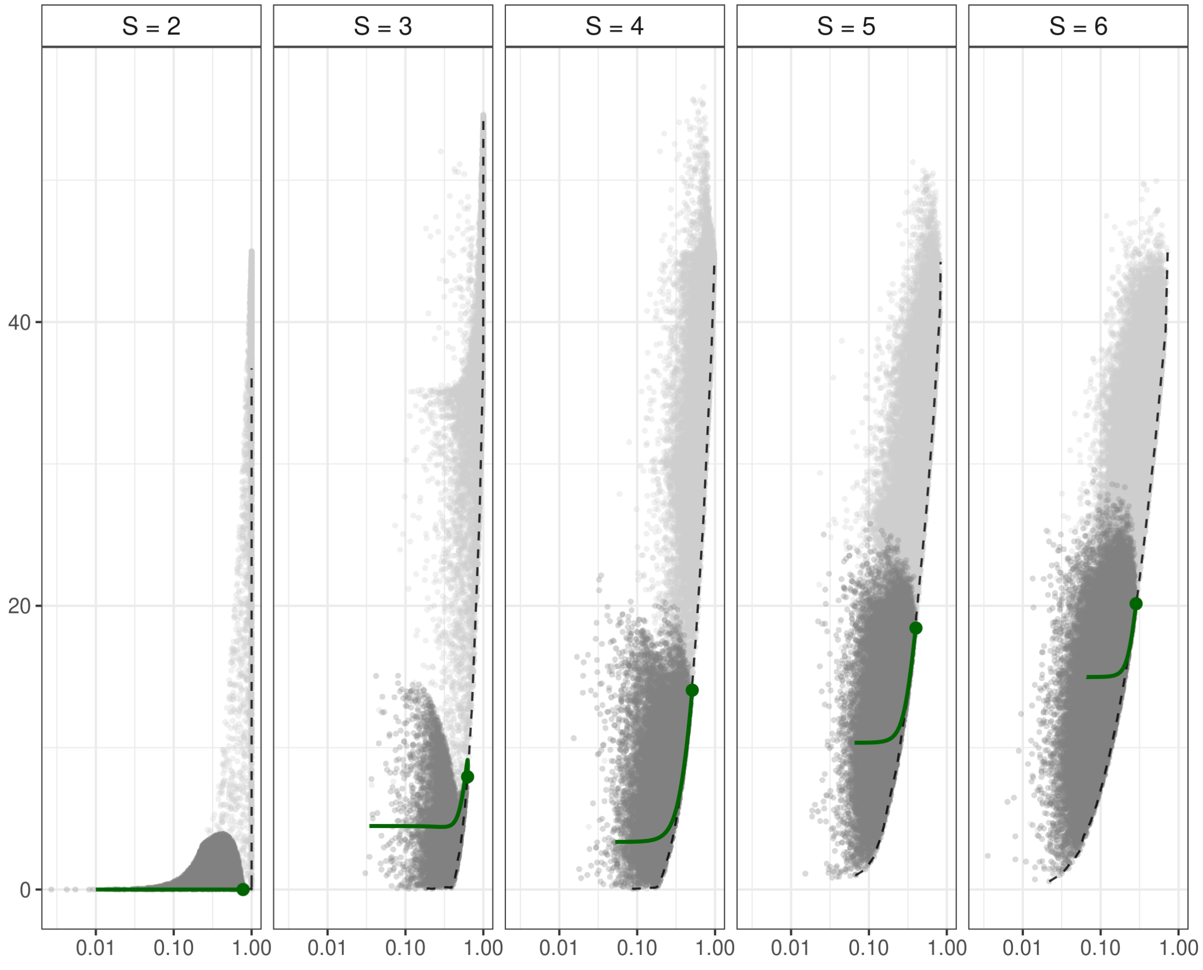


Figure S6.22: **Figure 4 for $\sigma_R$ = 1.6 (m = 0.01)**.

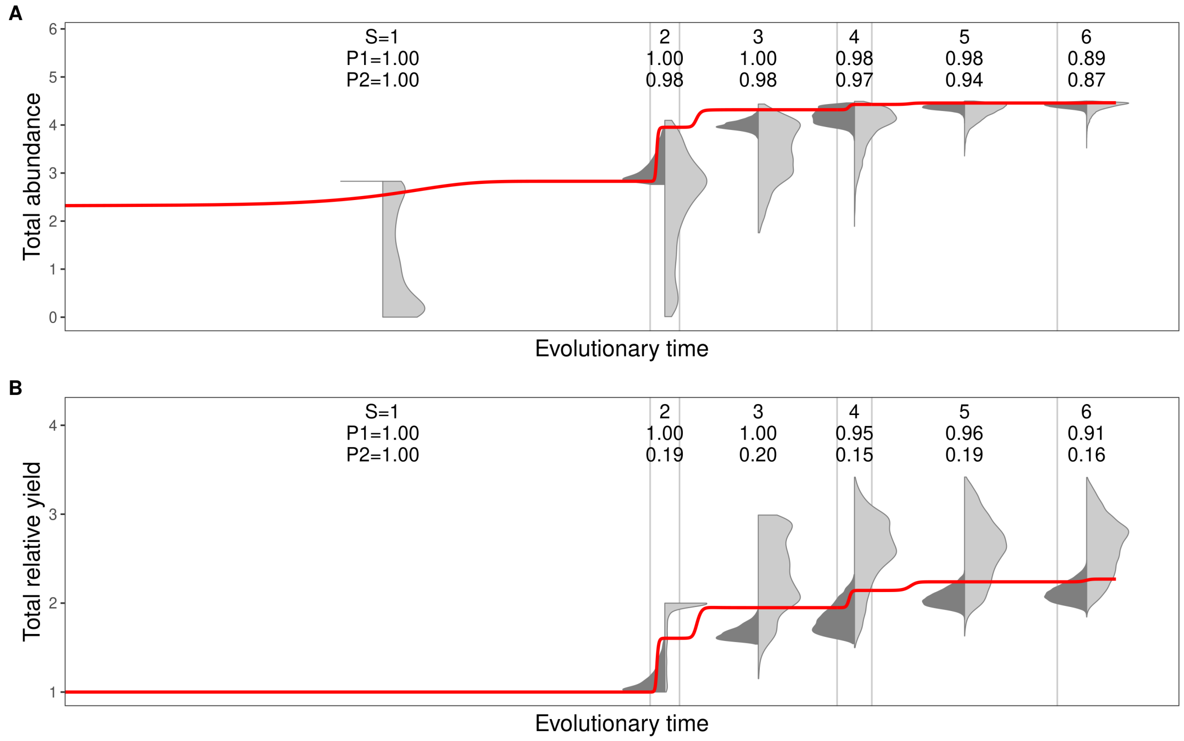


Figure S6.23: **Figure 5 for $\sigma_R$ = 1.6 (m = 0.01)**.

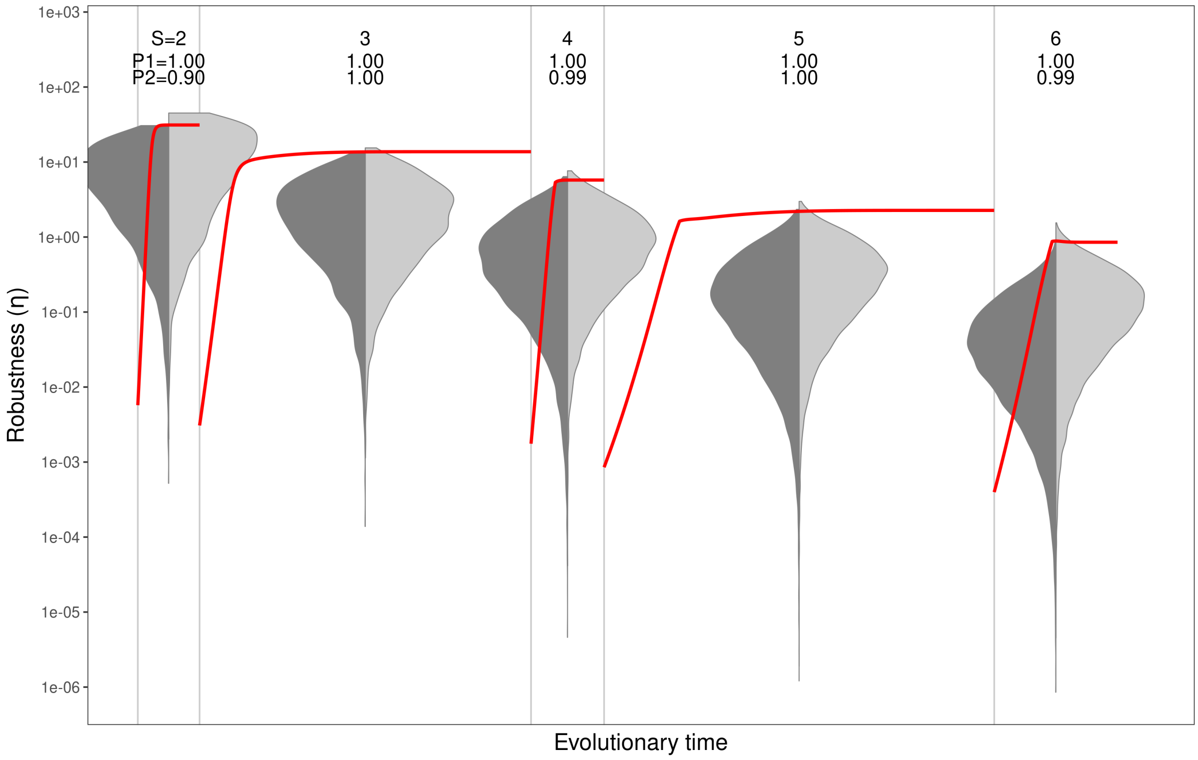


Figure S6.24: **Figure 6 for $\sigma_R$ = 1.6 (m = 0.01)**.

#### *Resource width* $\sigma_R = 1.7$

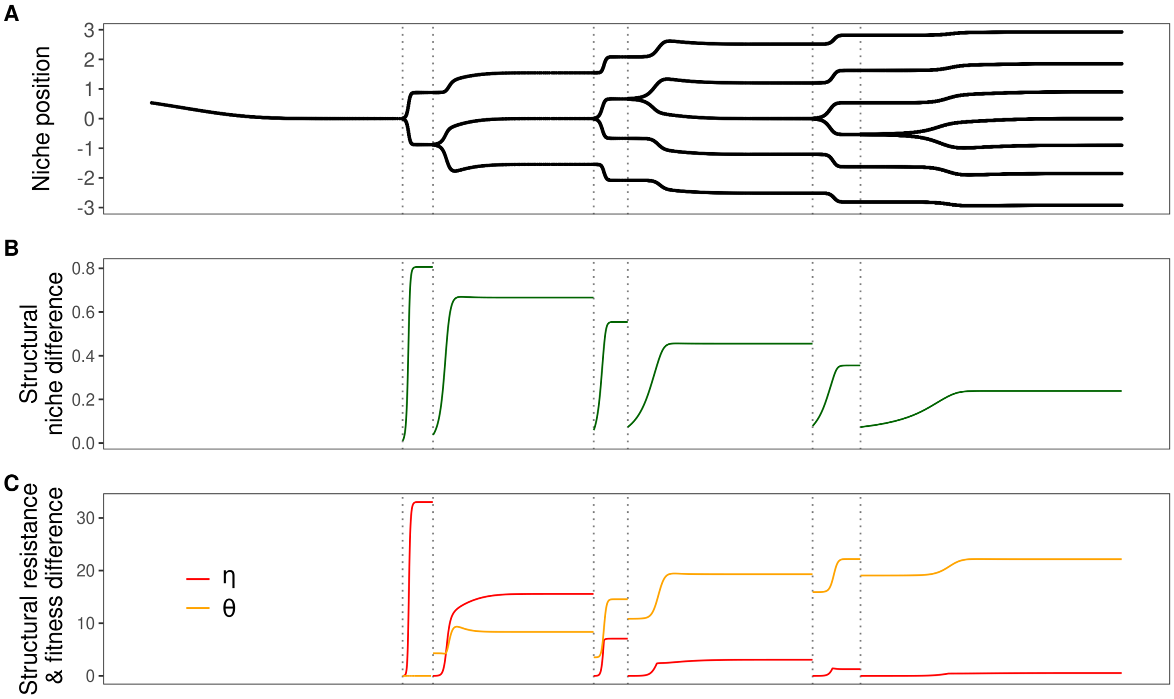


Figure S6.25: **Figure 2 for $\sigma_R = 1.7$ (m = 0.01)**.

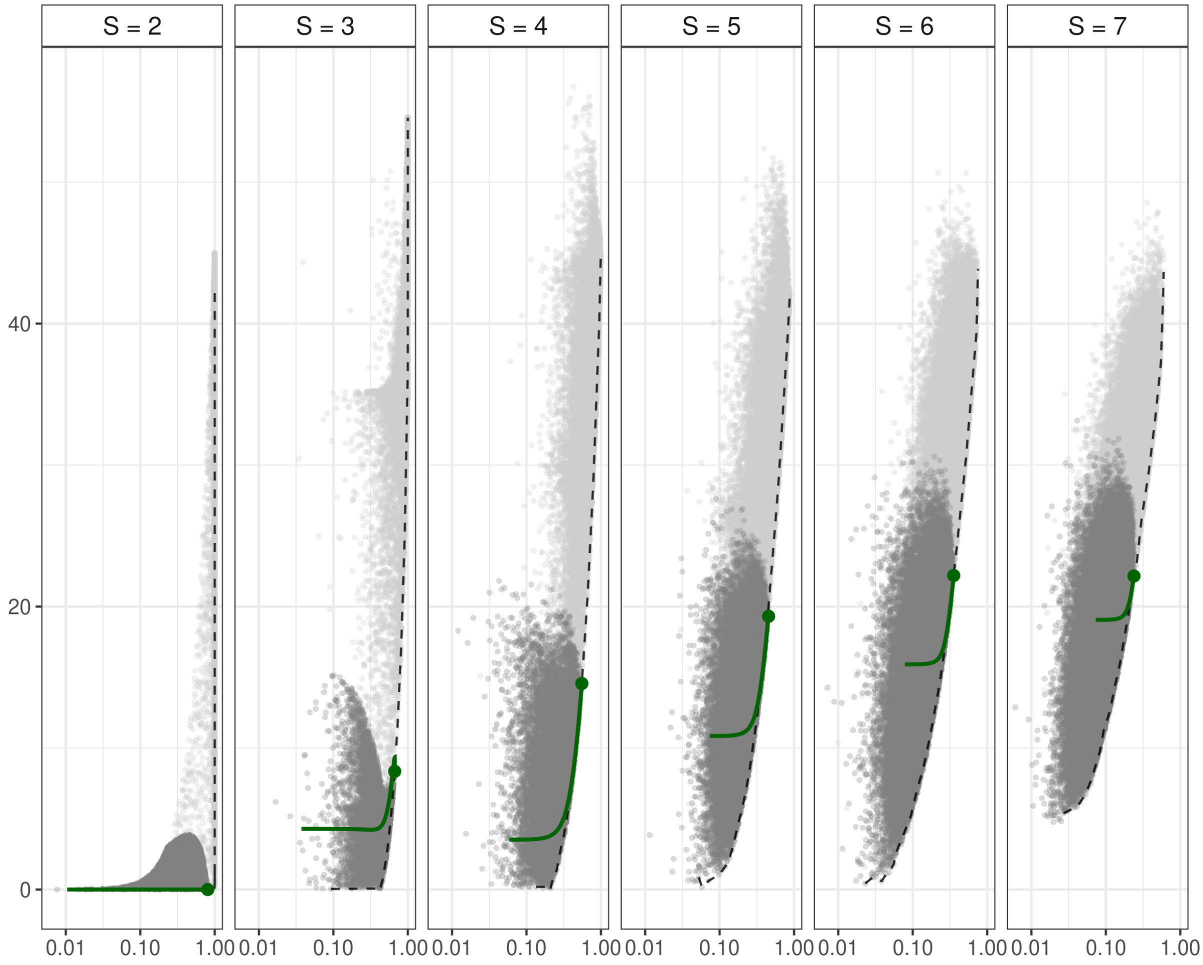


Figure S6.26: **Figure 4 for $\sigma_R = 1.7$ (m = 0.01)**.

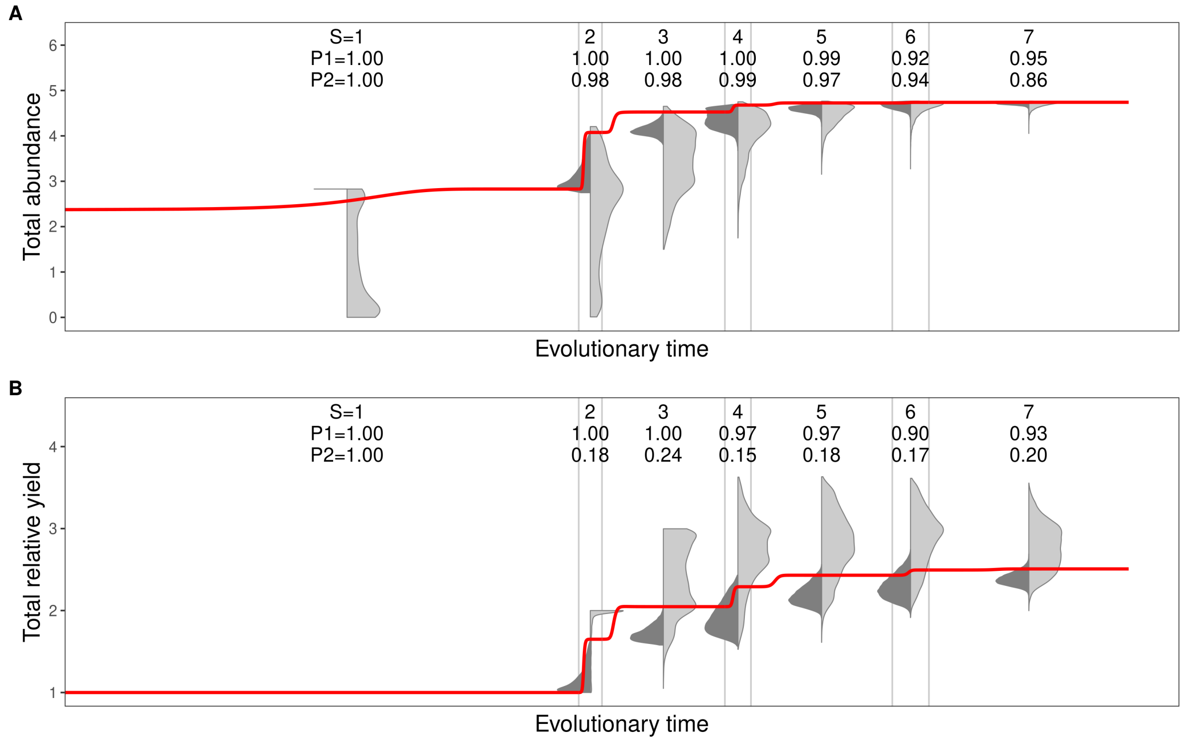


Figure S6.27: **Figure 5 for $\sigma_R$ = 1.7 (m = 0.01)**.

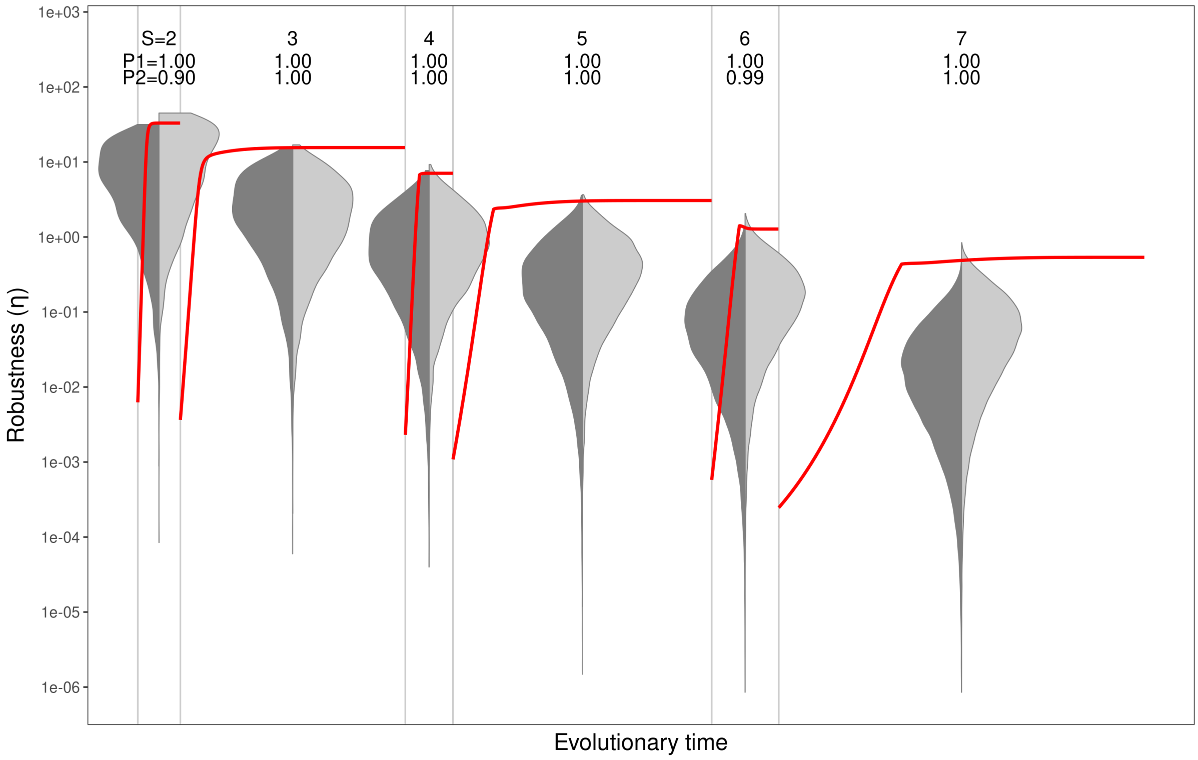


Figure S6.28: **Figure 6 for $\sigma_R$ = 1.7 (m = 0.01)**.

### *Resource width* $\sigma_R = 1.8$

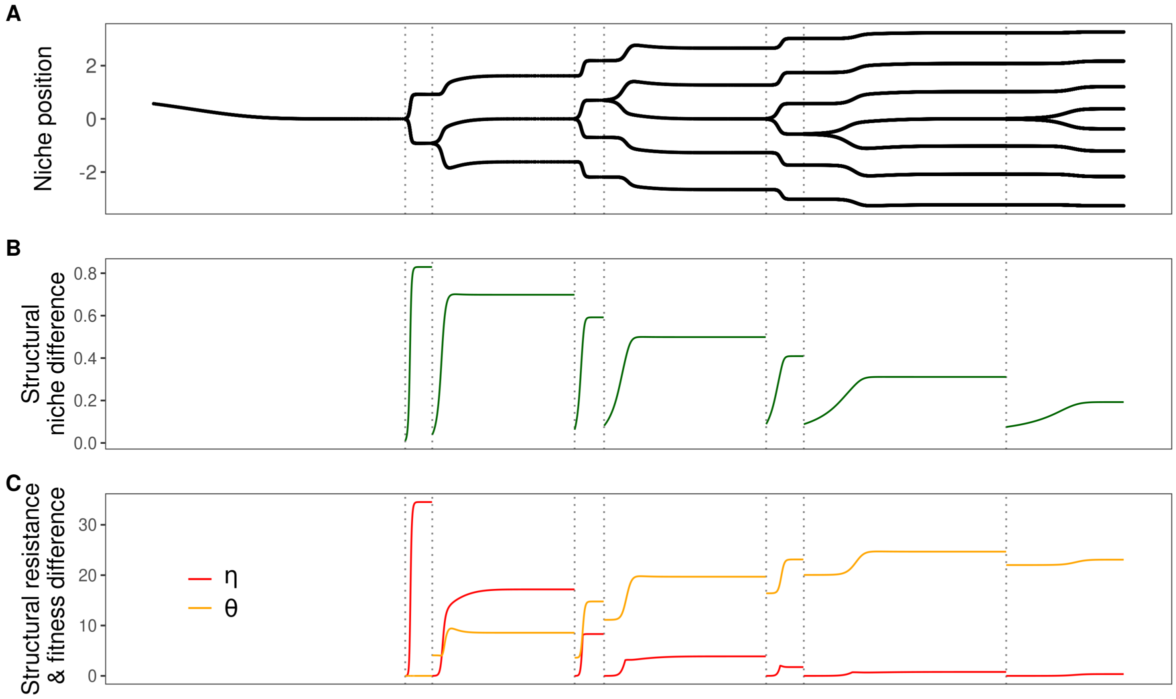


Figure S6.29: **Figure 2 for** $\sigma_R$ **= 1.8 (m = 0.01)**.

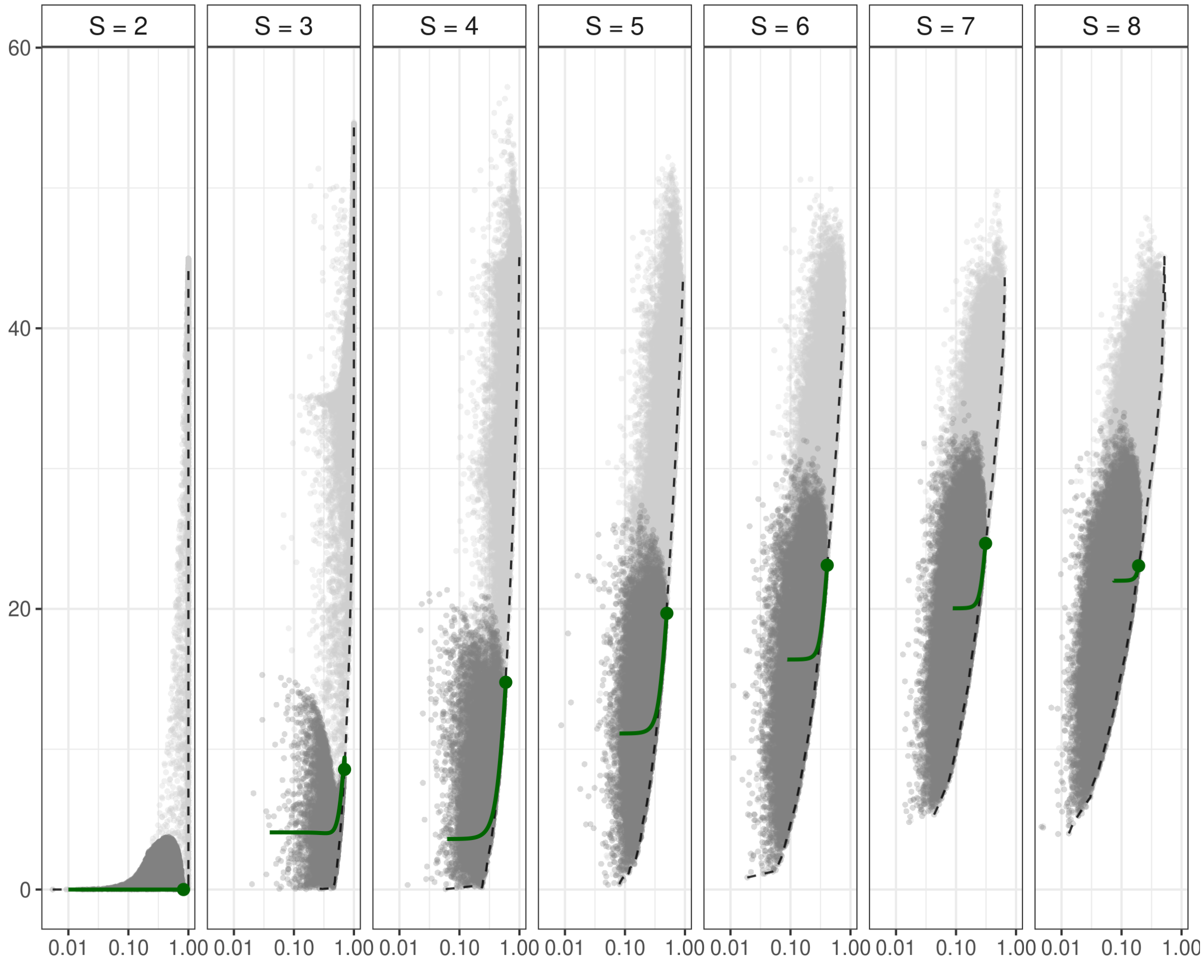


Figure S6.30: **Figure 4 for $\sigma_R$ = 1.8 (m = 0.01).**

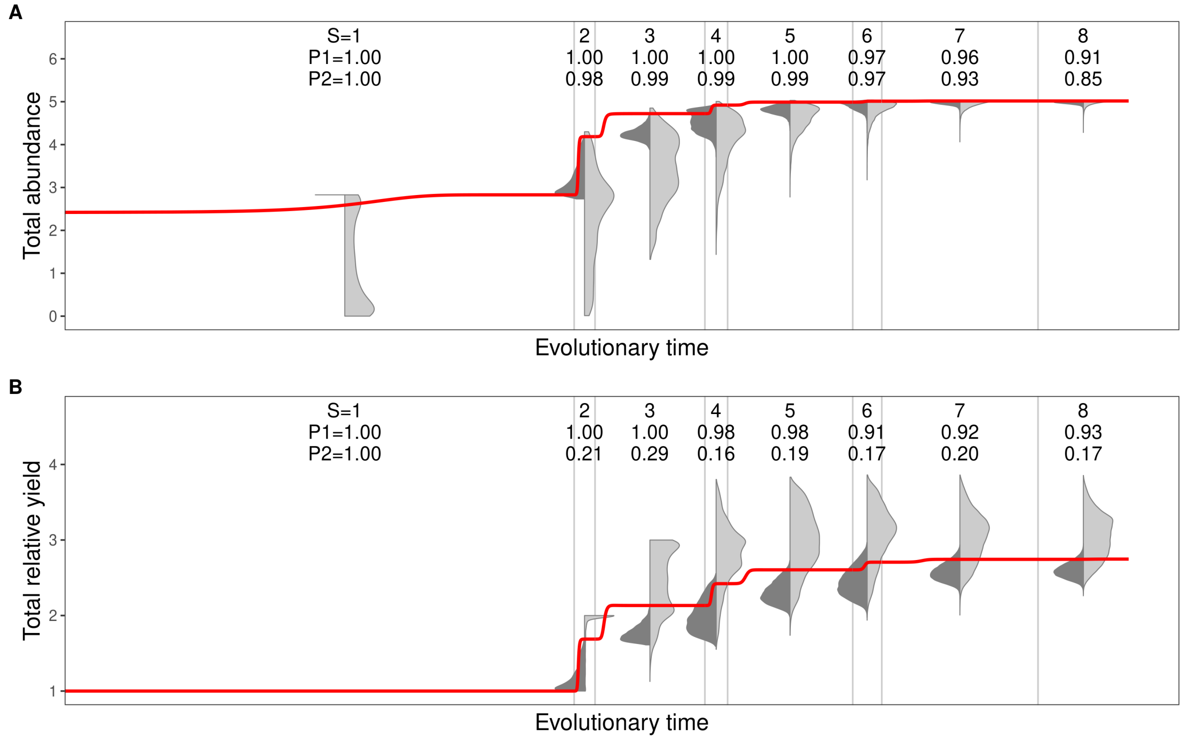


Figure S6.31: **Figure 5 for $\sigma_R = 1.8$ (m = 0.01)**.

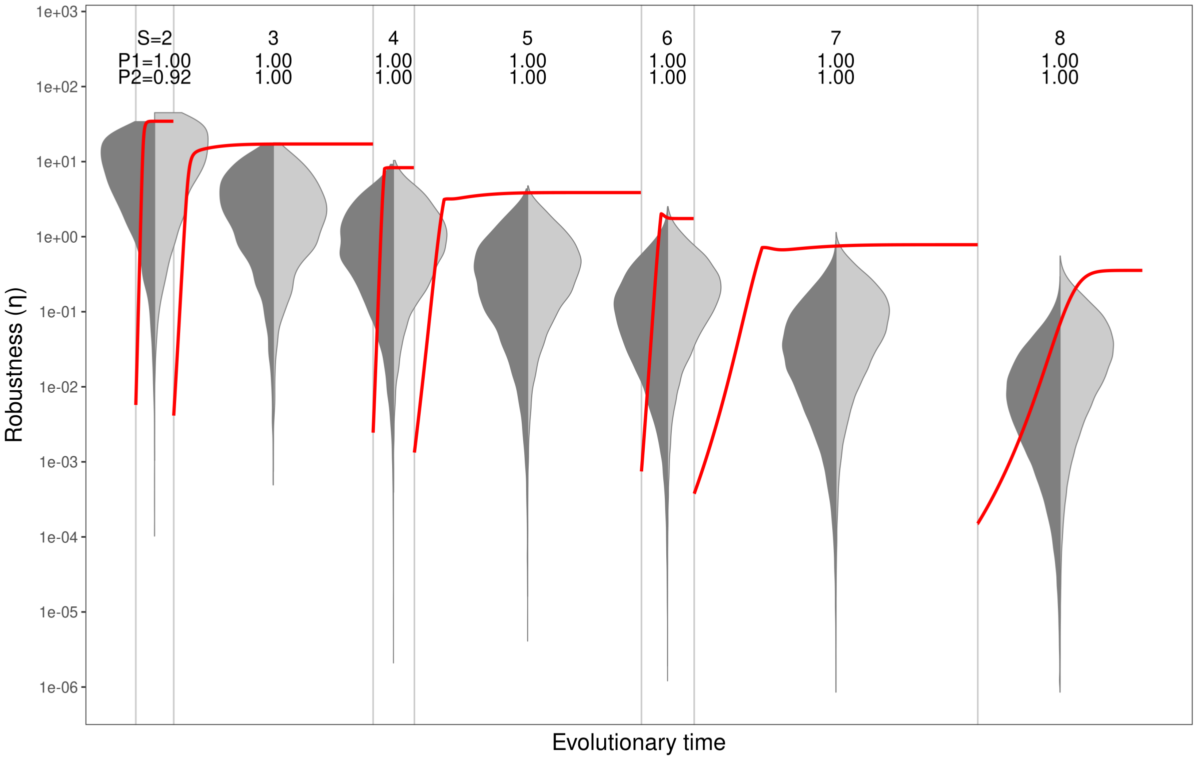


Figure S6.32: **Figure 6 for $\sigma_R = 1.8$ (m = 0.01)**.

87

#### *Resource width* $\sigma_R = 1.9$

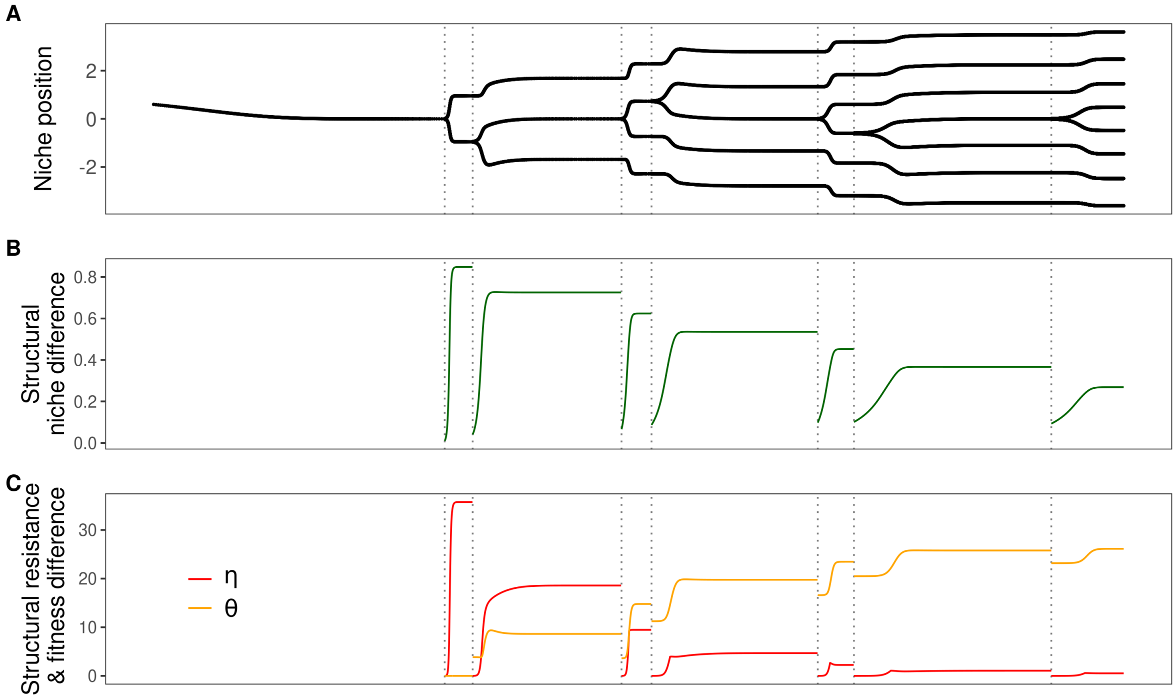


Figure S6.33: **Figure 2 for** $\boldsymbol{\sigma_R = 1.9}$ **(m = 0.01)**.

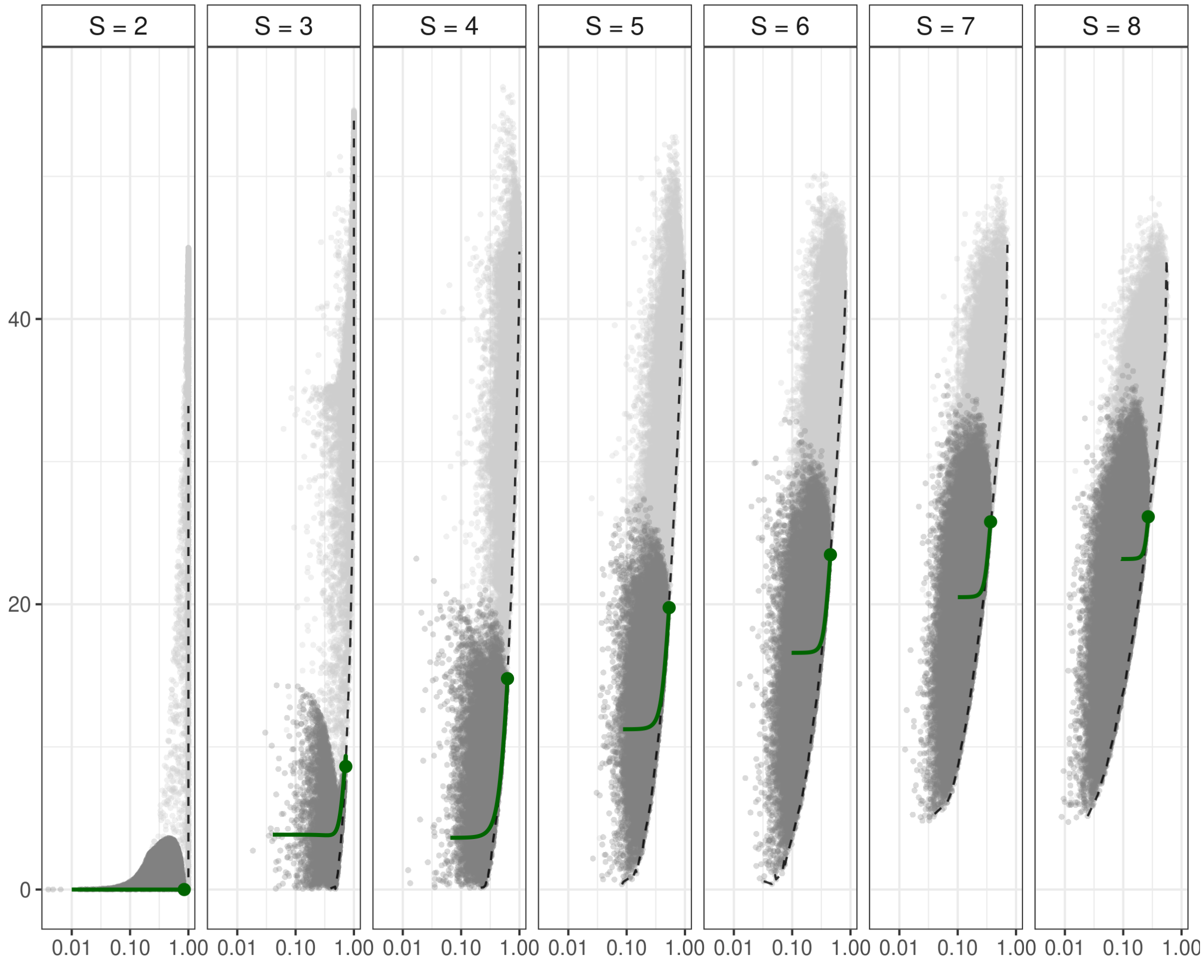


Figure S6.34: **Figure 4 for $\sigma_R = 1.9$ (m = 0.01)**.

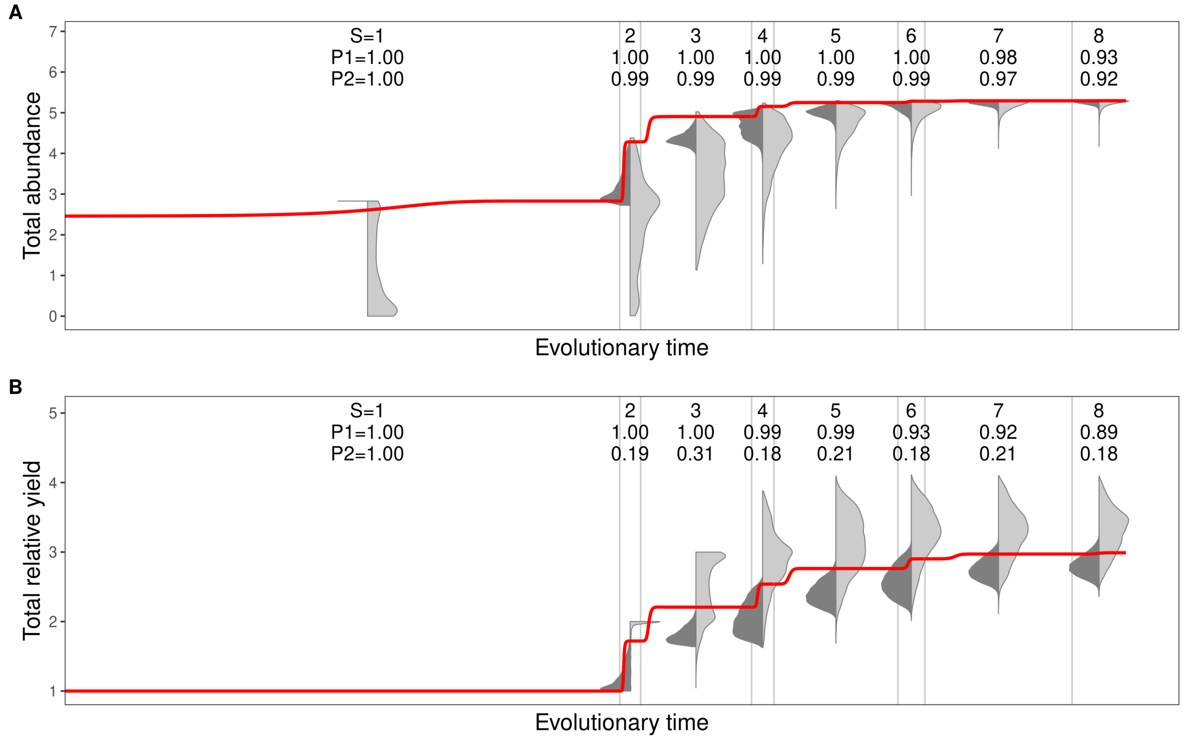


Figure S6.35: **Figure 5 for $\sigma_R$ = 1.9 (m = 0.01).**

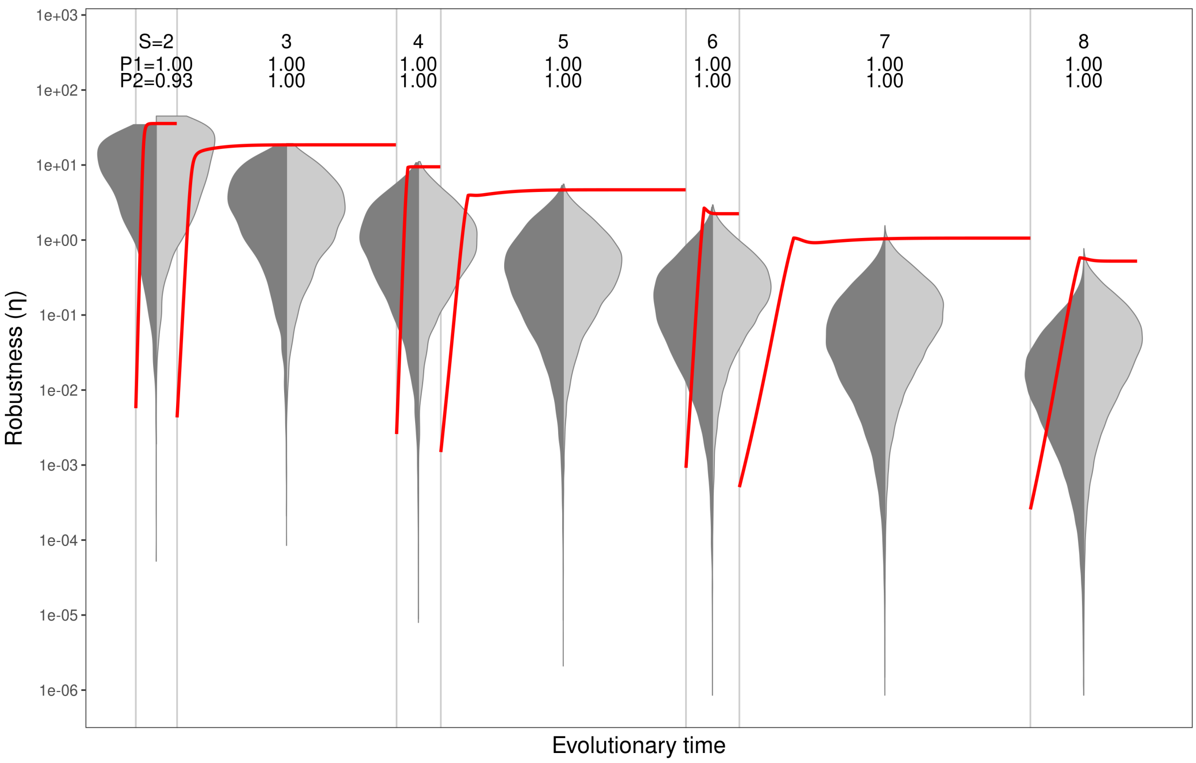


Figure S6.36: **Figure 6 for $\sigma_R$ = 1.9 (m = 0.01).**

### *Resource width* $\sigma_R = 2.0$

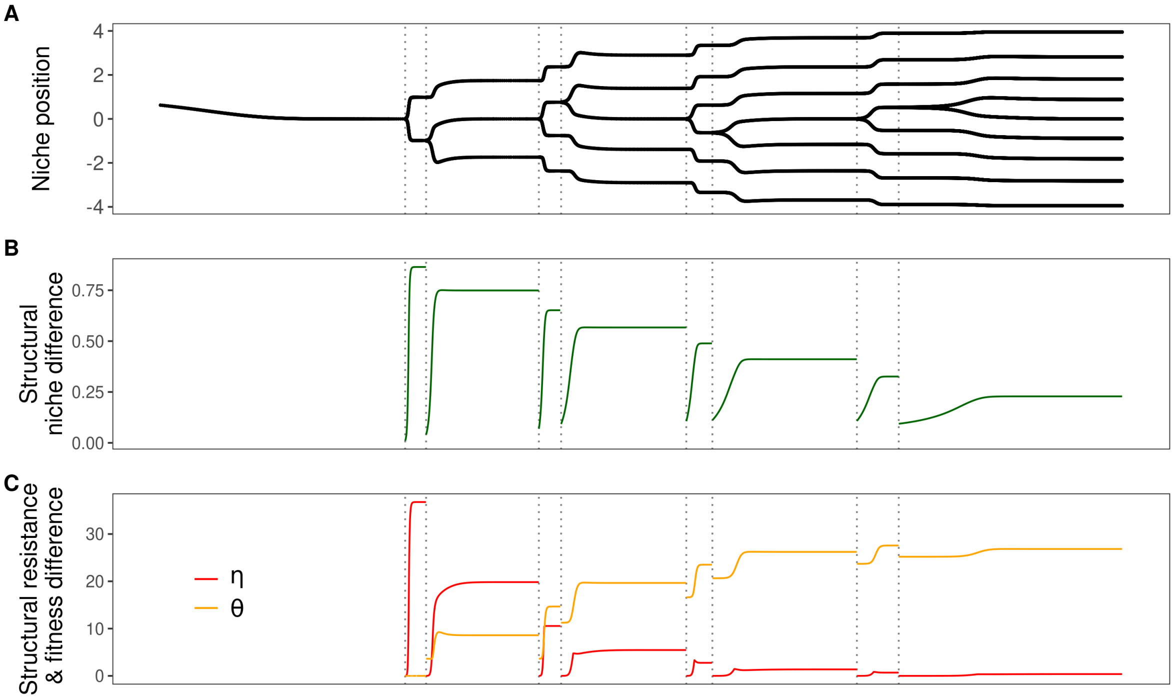


Figure S6.37: **Figure 2 for** $\sigma_R$ **= 2.0 (m = 0.01)**.

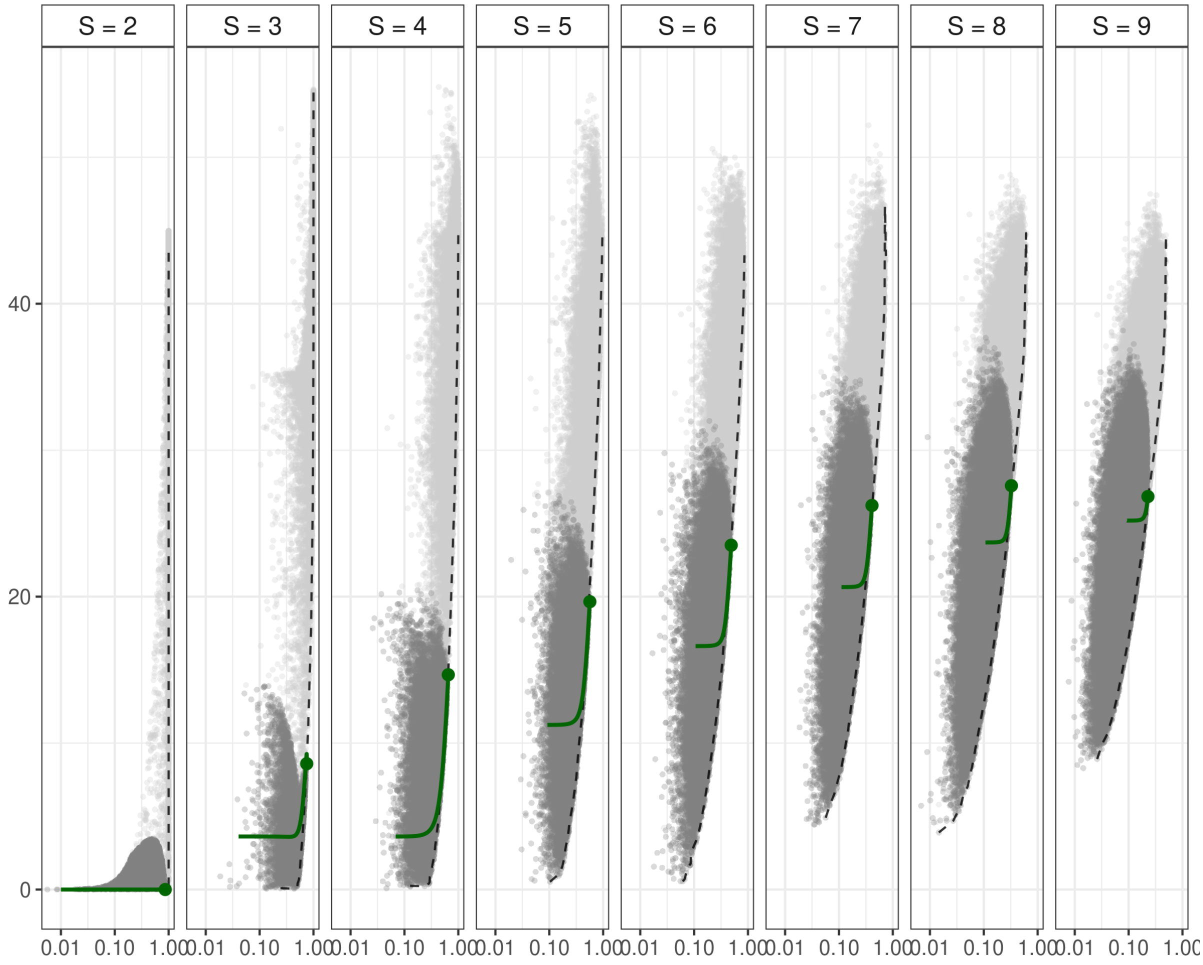


Figure S6.38: **Figure 4 for $\sigma_R$ = 2.0 (m = 0.01)**.

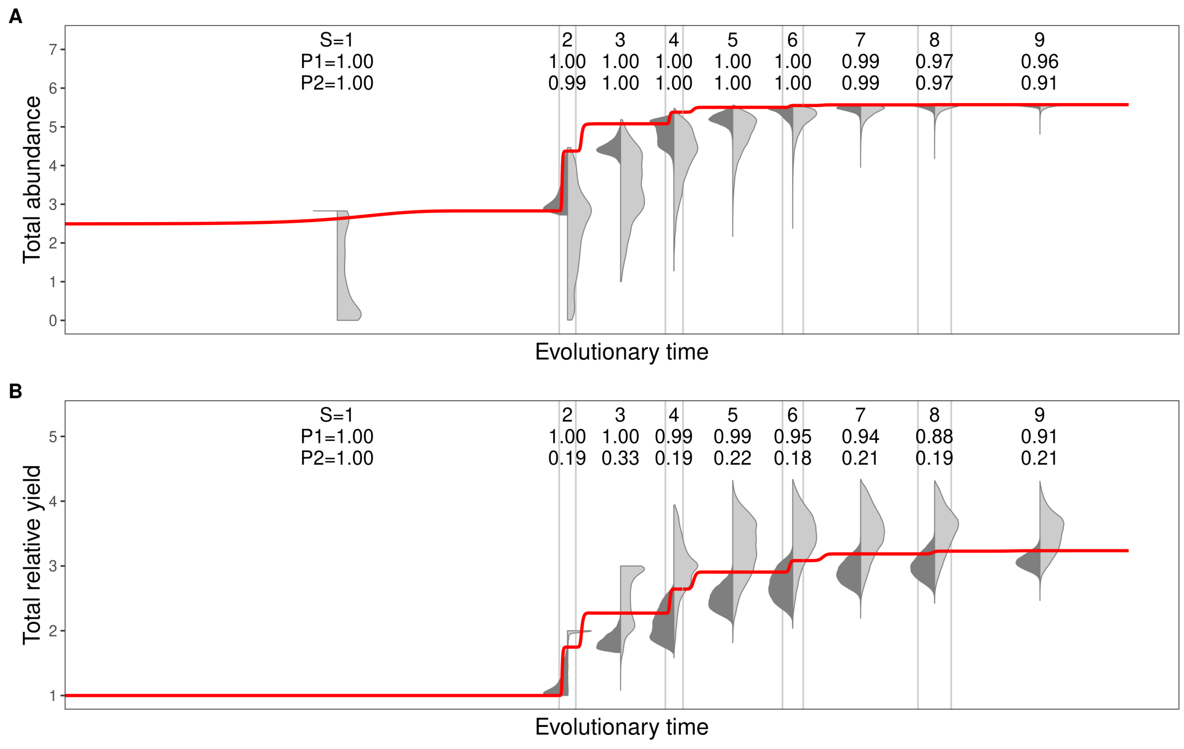


Figure S6.39: **Figure 5 for $\sigma_R = 2.0$ (m = 0.01)**.

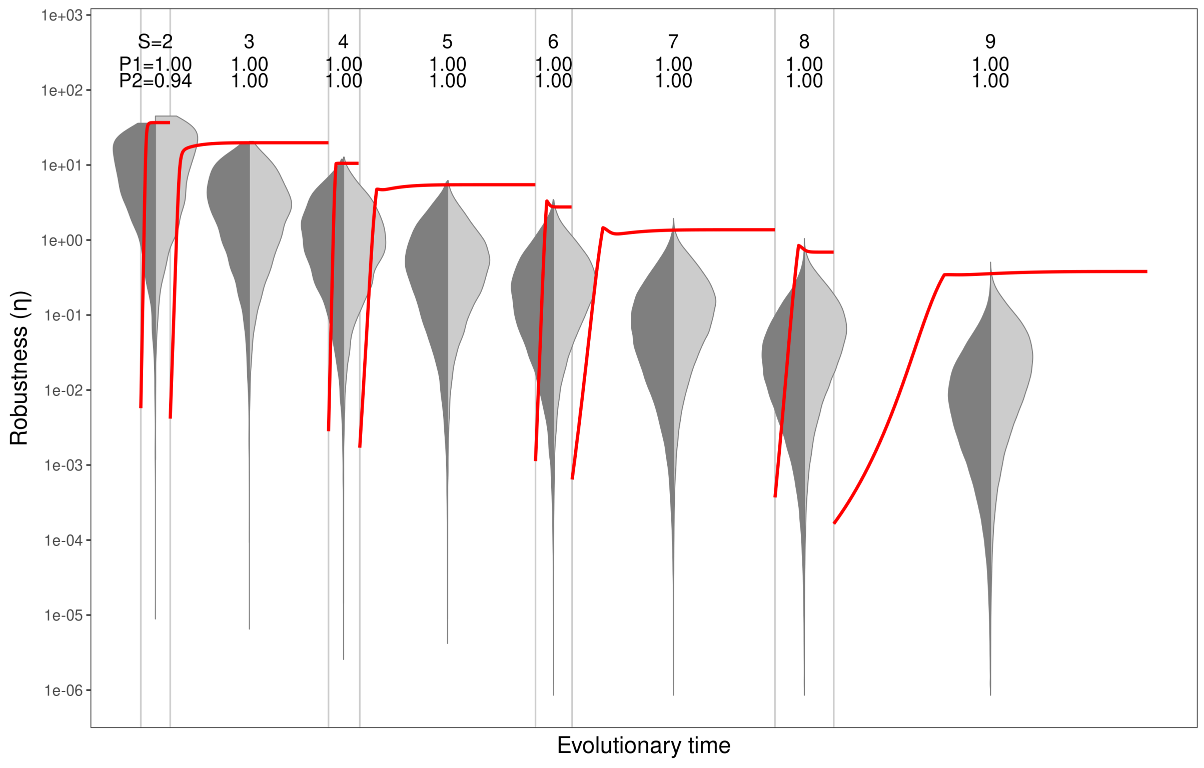


Figure S6.40: **Figure 6 for $\sigma_R = 2.0$ (m = 0.01)**.

### S7: Supplementary figures for different mortality rates ($m$)

$$\textit{Mortality } m = 0.1$$

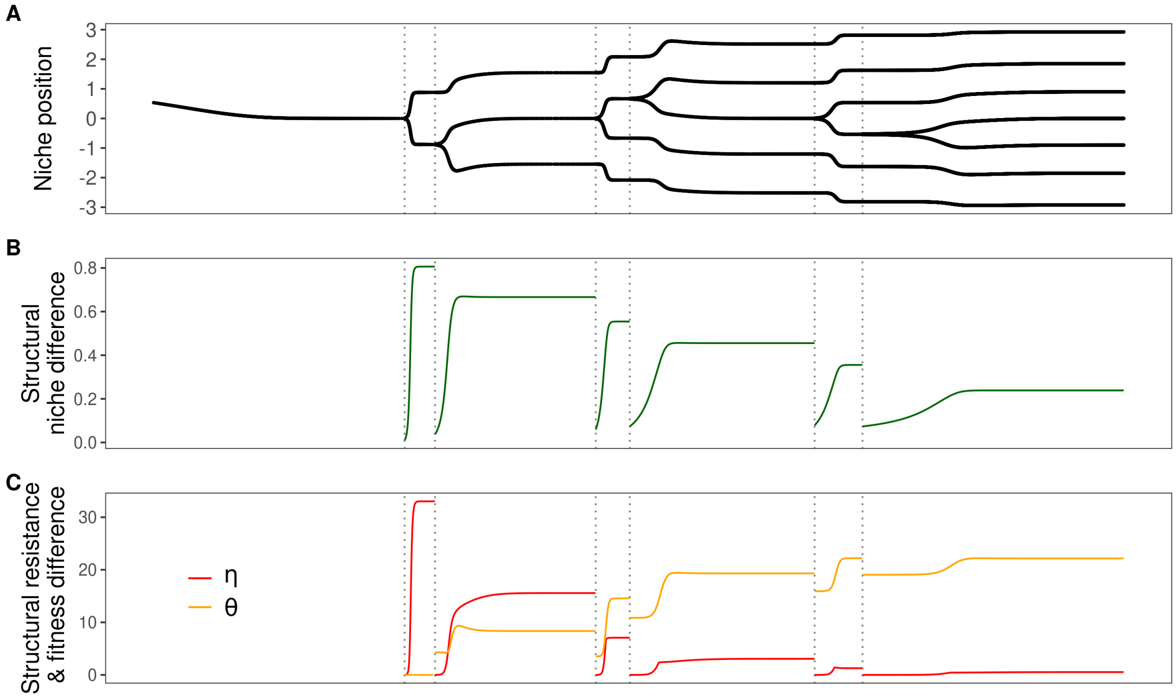


Figure S7.1: **Figure 2 for mortality = 0.1** .

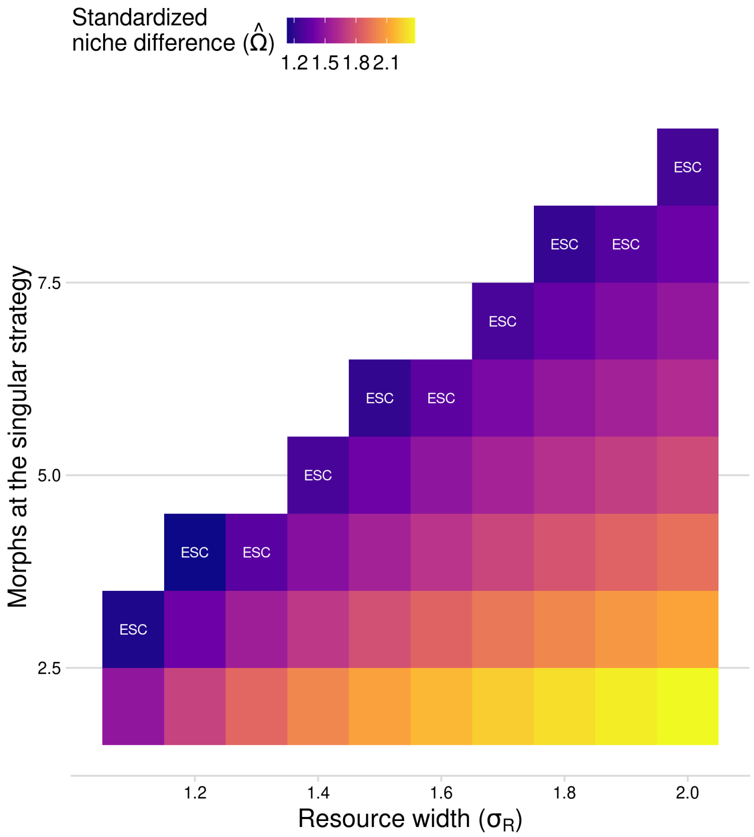


Figure S7.2: **Figure 3 for mortality = 0.1** .

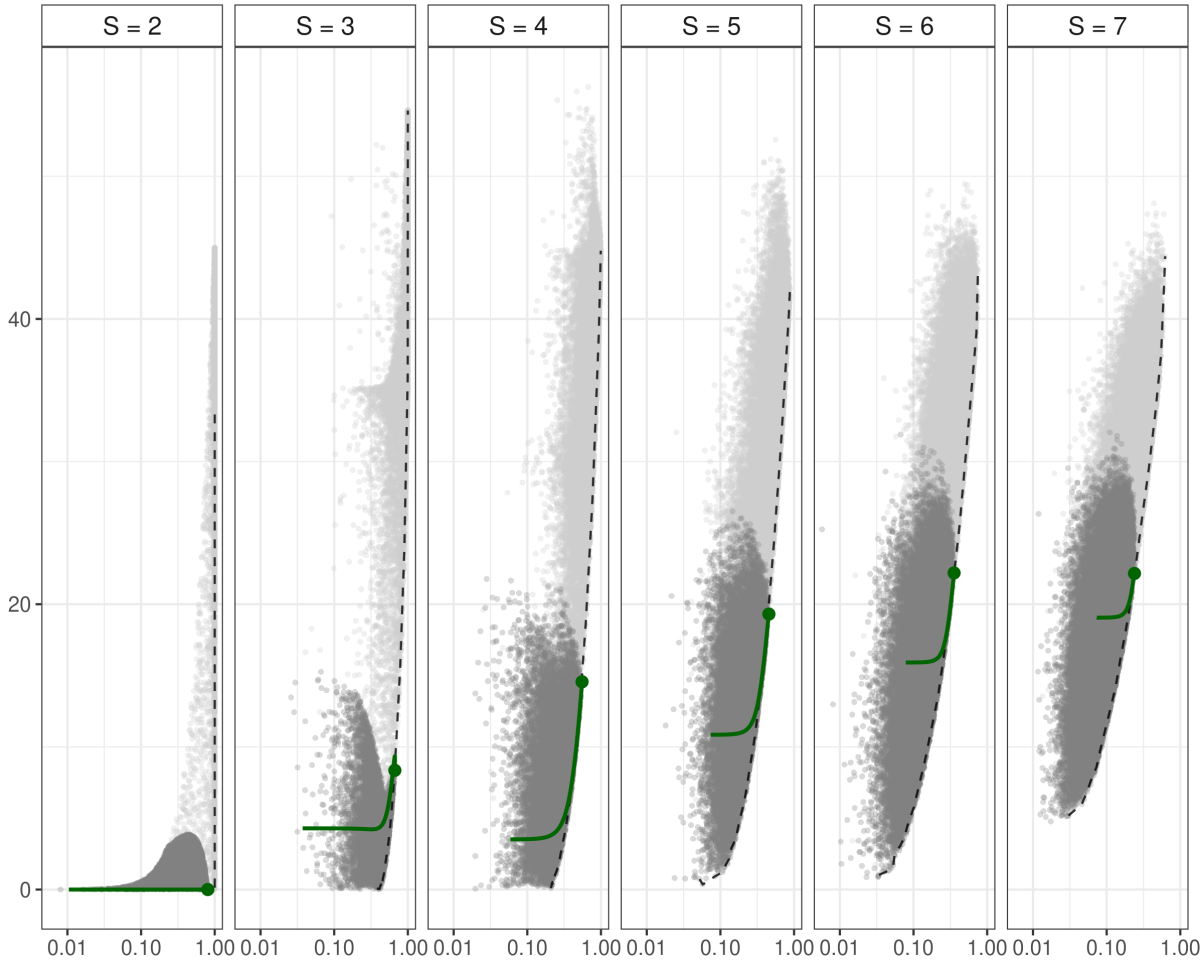


Figure S7.3: **Figure 4 for mortality = 0.1** .

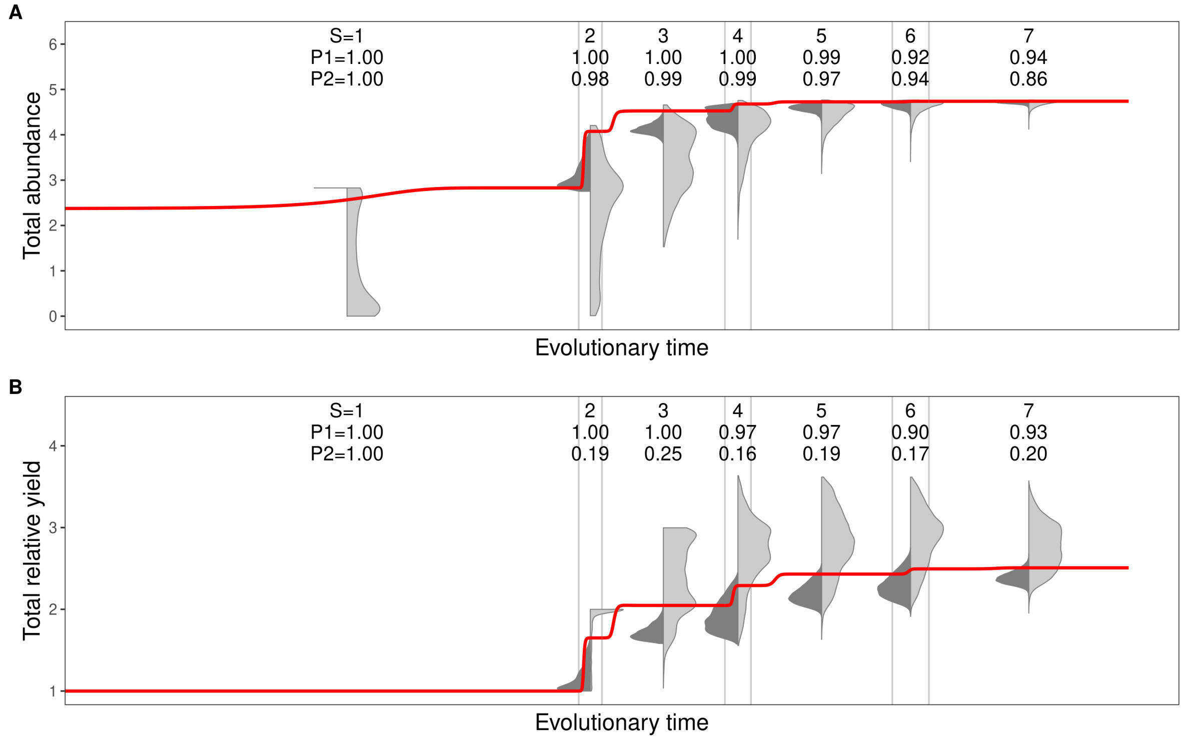


Figure S7.4: **Figure 5 for mortality = 0.1** .

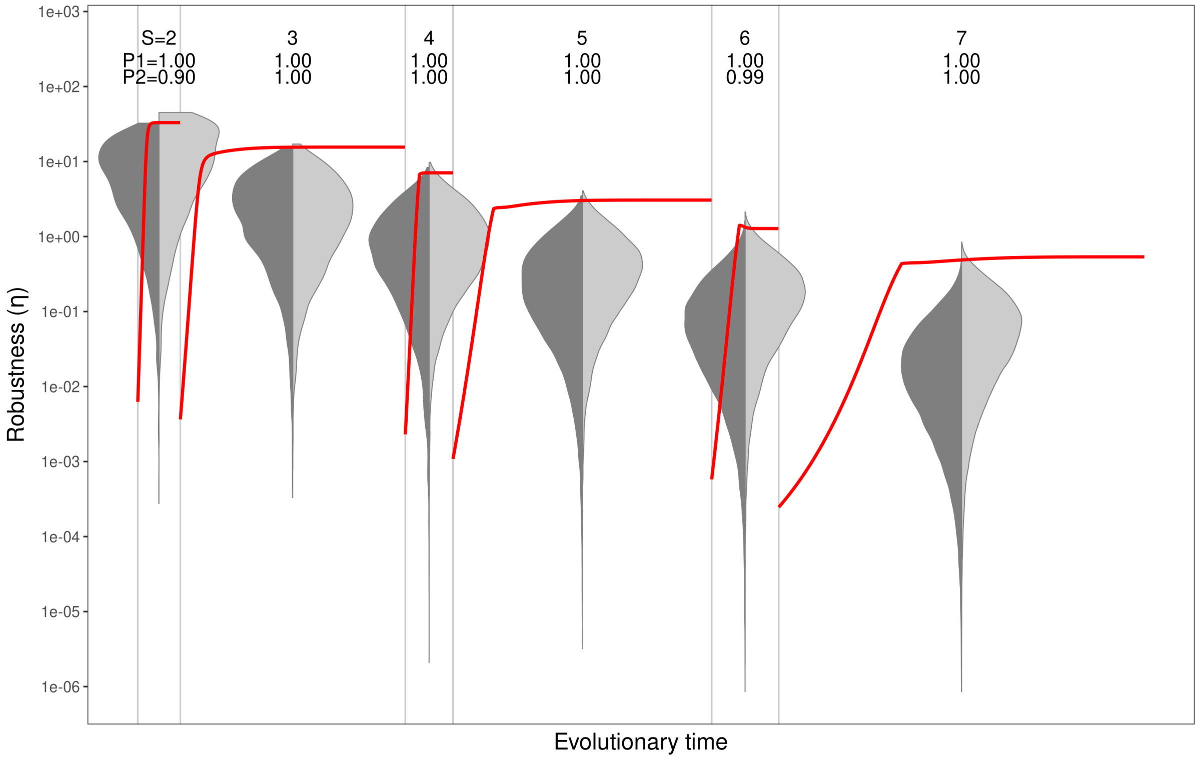


Figure S7.5: **Figure 6 for mortality = 0.1** .

#### *Mortality* $m = 0.5$

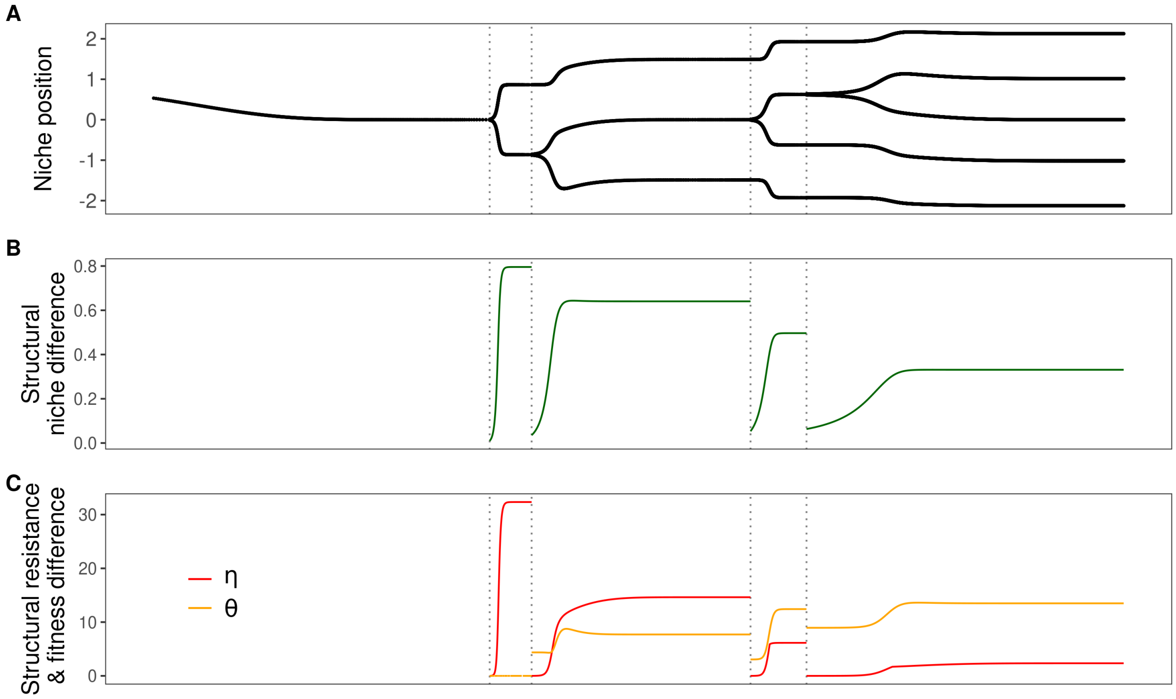


Figure S7.6: **Figure 2 for mortality = 0.5** .

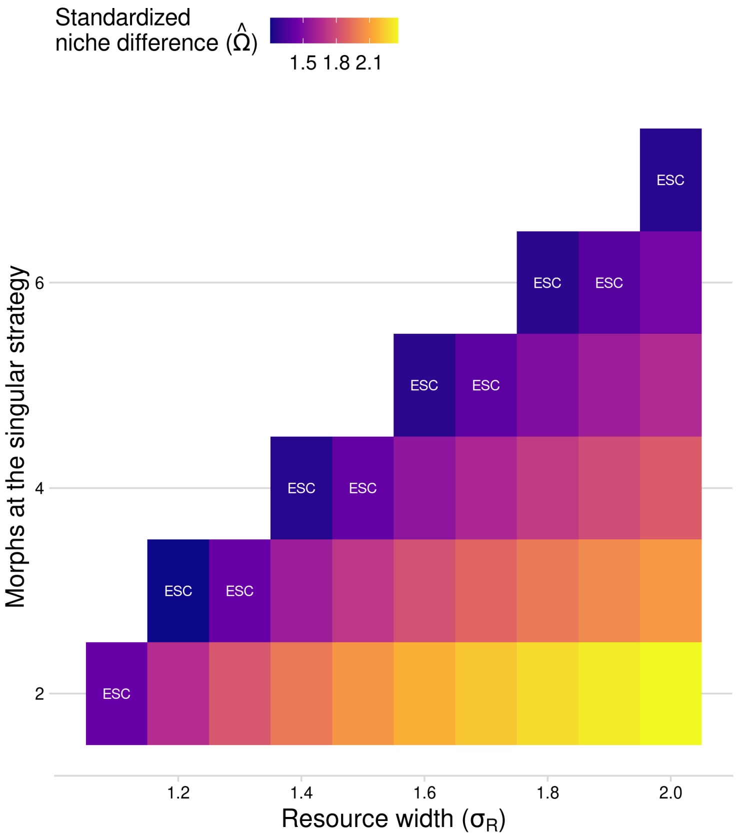


Figure S7.7: **Figure 3 for mortality = 0.5 .**

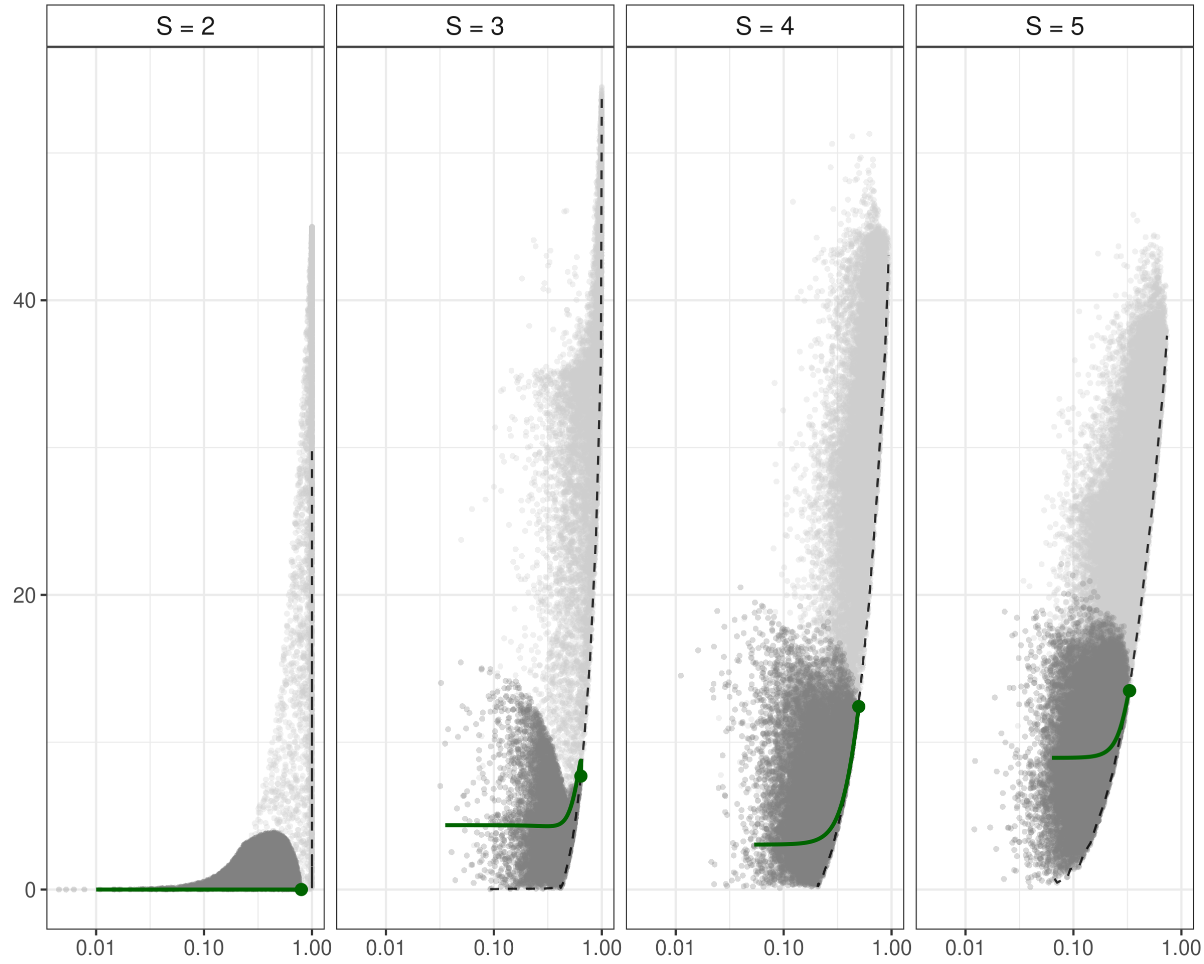


Figure S7.8: **Figure 4 for mortality = 0.5** .

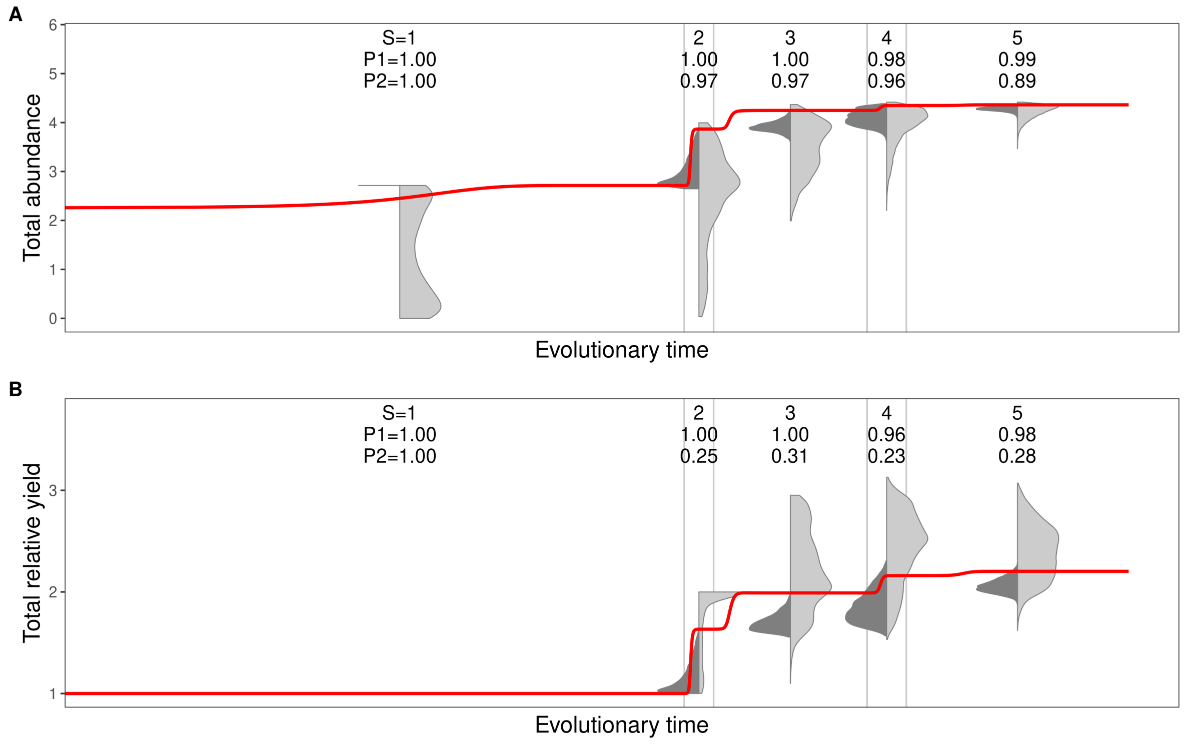


Figure S7.9: **Figure 5 for mortality = 0.5** .

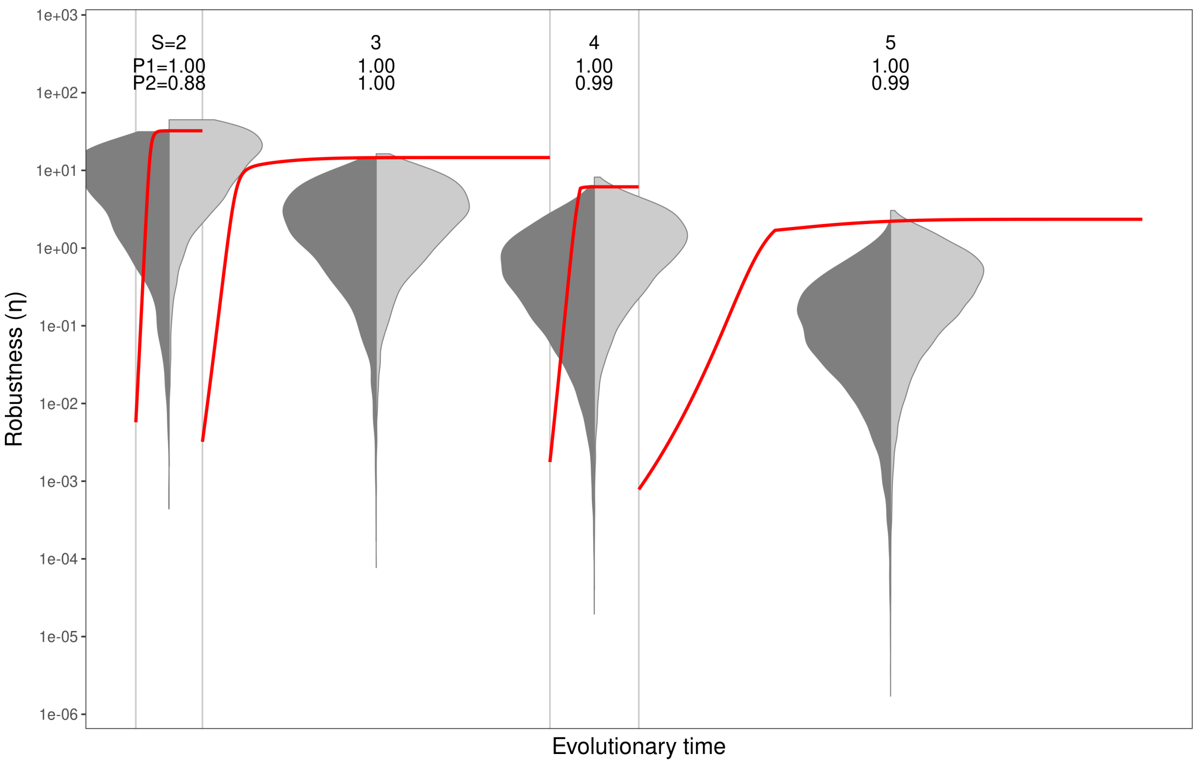


Figure S7.10: **Figure 6 for mortality = 0.5** .

92

#### *Mortality* $m = 1.0$

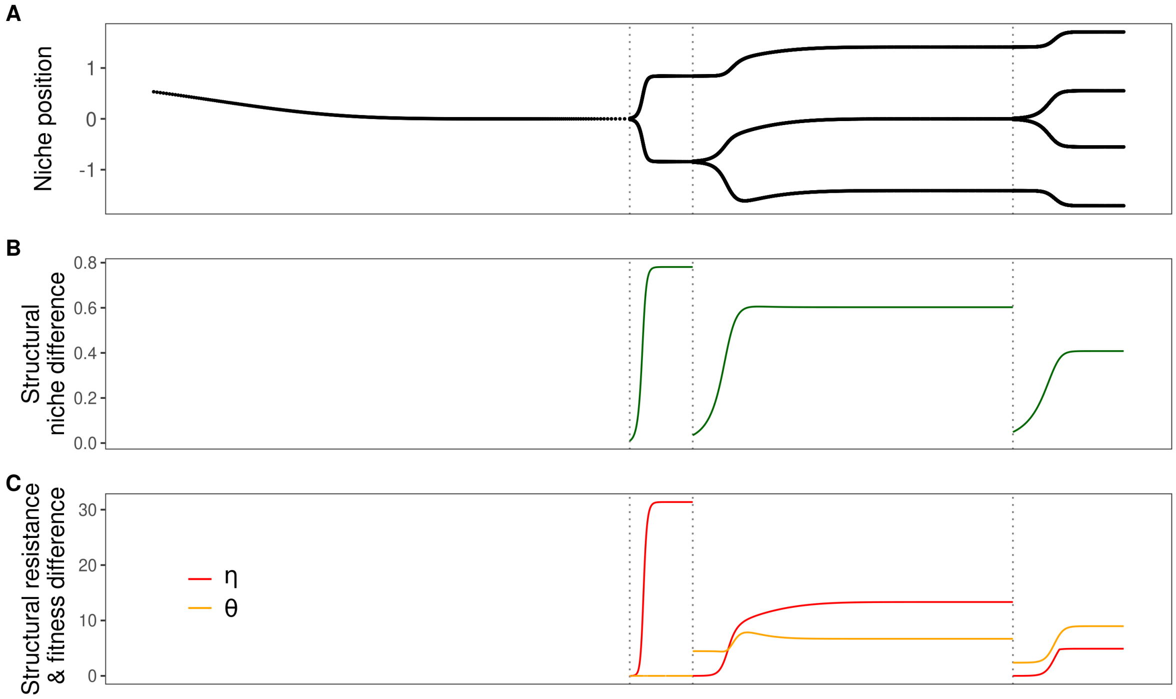


Figure S7.11: **Figure 2 for mortality = 1.0** .

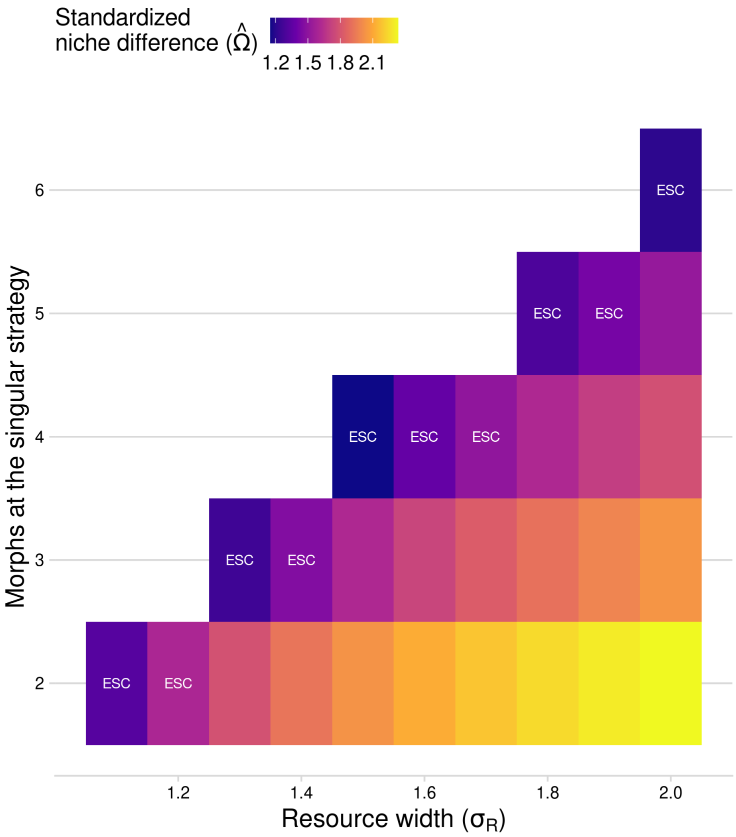


Figure S7.12: **Figure 3 for mortality = 1.0 .**

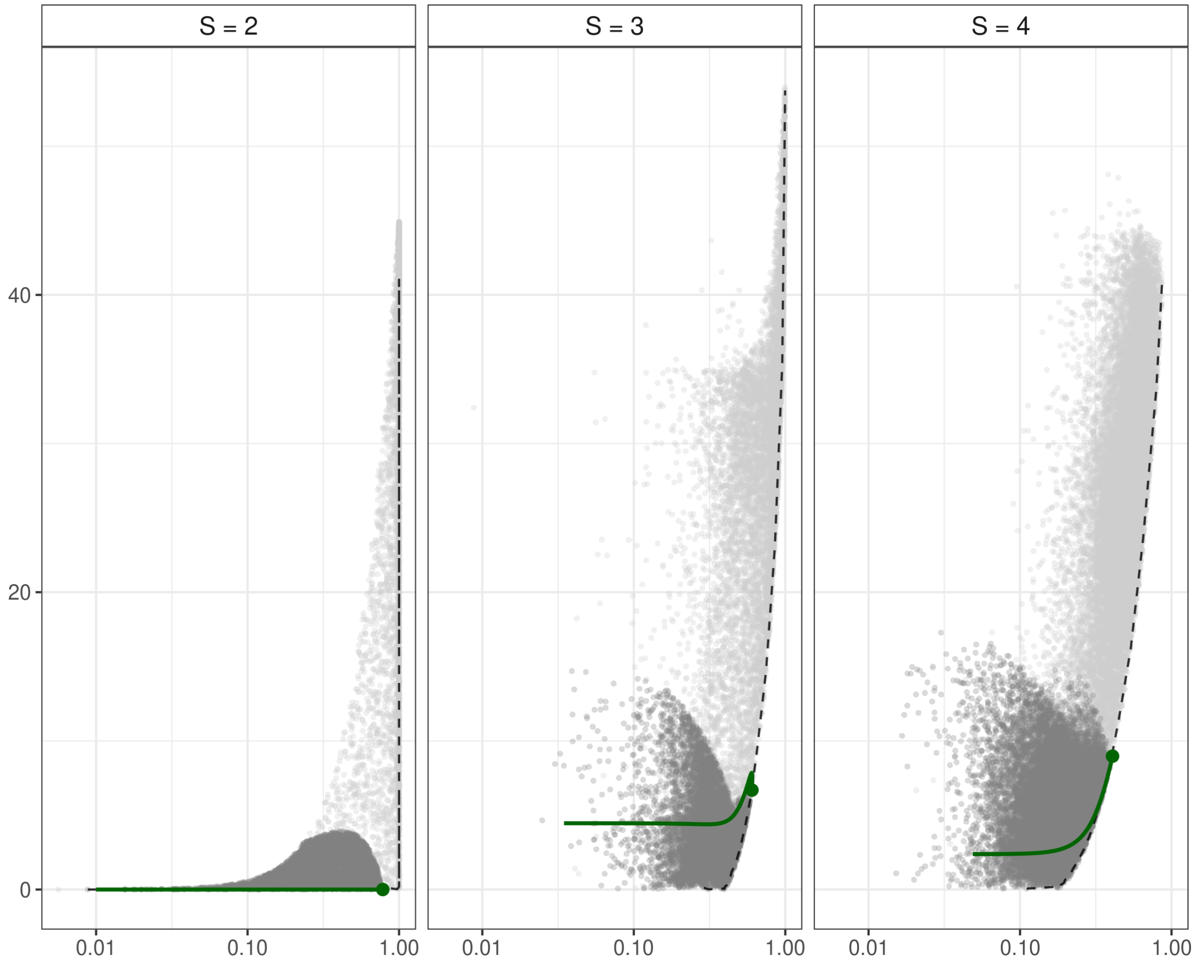


Figure S7.13: **Figure 4 for mortality = 1.0** .

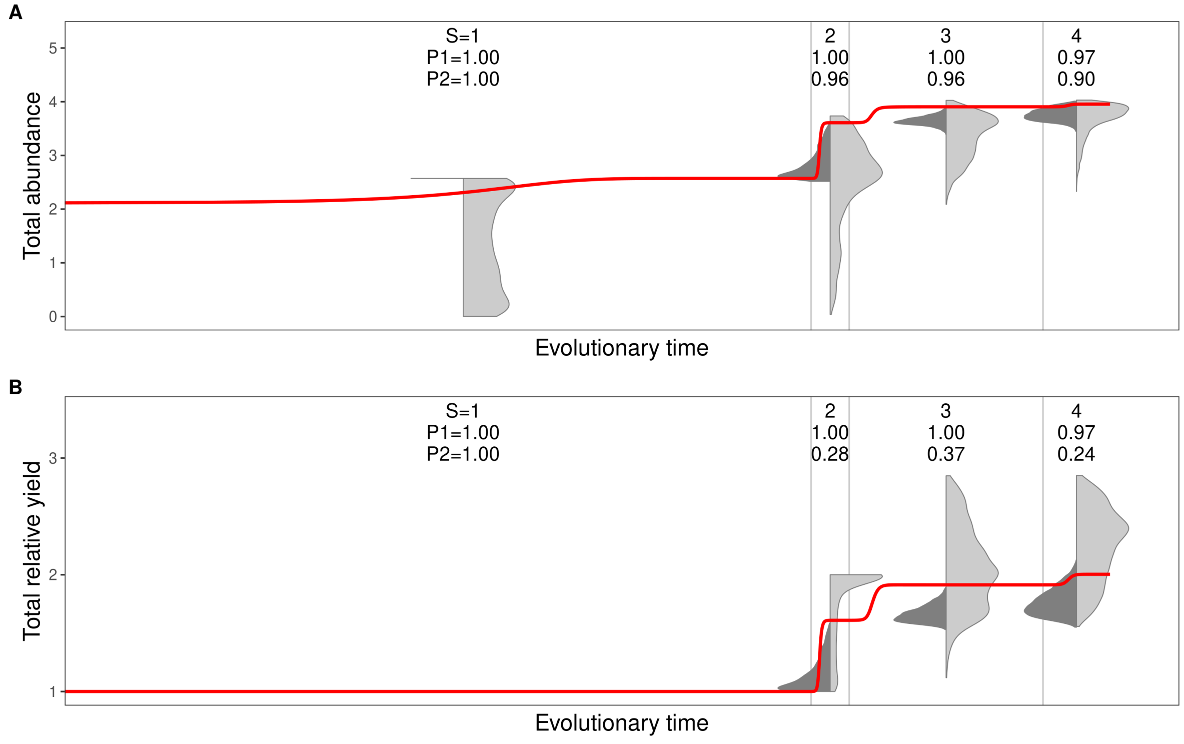


Figure S7.14: **Figure 5 for mortality = 1.0** .

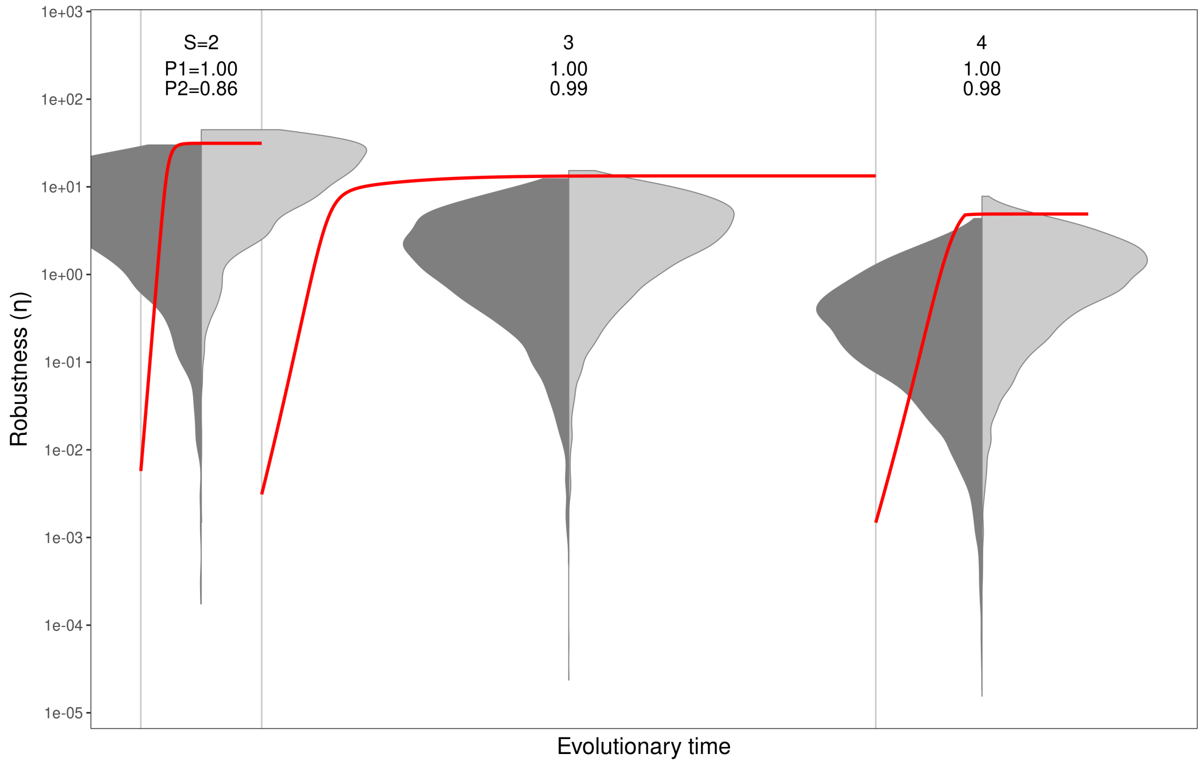


Figure S7.15: **Figure 6 for mortality = 1.0** .

### S8: Box-like intrinsic growth functions

Figure 1 in the case where $r(x)$ is a box-car function. To obviate the problem of a non-differentiable discontinuity at the edges, we use a piecewise function which decreases polinomially and is hence everywhere twice differentiable.

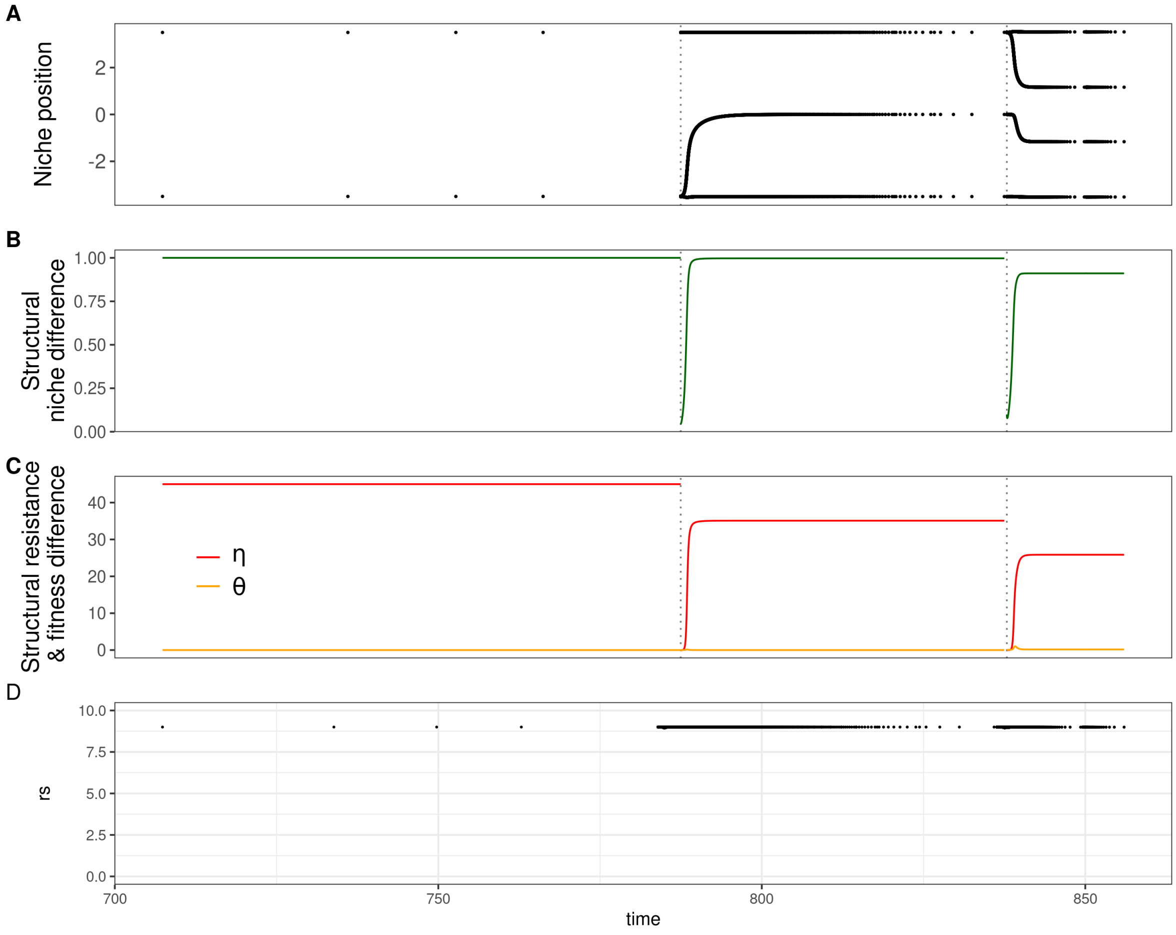


Figure S8.1: **Figure 1 for a box-like intrinsic growth rate .**
